# Single Nucleus MultiOmics Links Novel Transcription Factor Motifs to Murine Hepatic Sex Differences in Chromatin Accessibility and Metabolic Dysfunction-Associated Steatotic Liver Disease

**DOI:** 10.1101/2025.10.31.685903

**Authors:** Kritika Karri, Ting-Ya Chang, Maxim Pyatkov, Shashi Gandhi, Trevor Siggers, David J. Waxman

## Abstract

The liver exhibits striking sexual dimorphism in gene expression that impacts drug and lipid metabolism and disease susceptibility, with males showing substantially higher predisposition to metabolic dysfunction-associated steatotic liver disease (MASLD) and its complications including hepatocellular carcinoma. These sex differences are primarily controlled by sexually dimorphic pituitary growth hormone (GH) secretion patterns; however, the underlying transcriptional and epigenetic regulatory networks remain only partially understood. Here, we generated paired single-nucleus chromatin accessibility (snATAC-seq) and gene expression (snRNA-seq) profiles from 46,188 liver nuclei isolated from male, female and continuous GH-infused male mice to comprehensively map the epigenetic basis of hepatic sexual dimorphism. We identified 127,957 accessible chromatin regions genome-wide, including thousands of novel regions enriched specifically in non-parenchymal cells. Sex-biased differentially accessible chromatin regions (DARs) were almost exclusively hepatocyte-localized, and continuous GH infusion feminized their accessibility, demonstrating that plasma GH patterns alone are sufficient to reprogram sex-biased hepatocyte chromatin landscapes. Correlation-based peak-to-gene linkage analysis mapped these DARs to sex-biased gene targets and revealed that regulatory interactions are constrained by topologically associated domain boundaries. Motif enrichment analysis identified both established regulators (STAT5, CUX2, BCL6) and novel transcription factors (TFs) at sex-biased DARs. ATAC-seq footprinting revealed novel TF motifs predicted to be occupied at DARs linked to sex-biased genes implicated in MASLD, providing mechanistic insights into the male bias in fatty liver disease. Further, motif co-occurrence analysis revealed TF clusters likely cooperating to regulate sex-dependent gene expression programs. We also identified stringently cell type-specific regulatory regions with cell type-specific TF motifs that define the regulatory architecture underlying hepatocyte and non-parenchymal cell identities. This comprehensive multiOmic atlas elucidates TF networks controlling sex-dependent liver gene expression and serves as a foundational resource for understanding molecular mechanisms underlying sex disparities in MASLD and other liver diseases.

## Introduction

The liver shows striking sex differences in the expression of hundreds of genes impacting liver physiology and disease, a majority of which are regulated by the sex-dependent temporal patterns of pituitary growth hormone (GH) secretion [1]. Many of these genes metabolize lipids and steroids to help meet sex-specific physiological requirements, including those related to metabolism of fluctuating steroid levels during pregnancy and the estrus cycle [2]. Liver sex differences also impact drug metabolism and pharmacokinetics [3] and liver disease [4, 5], with females more susceptible to alcoholic liver disease [6] and primary biliary cirrhosis [7], and males more susceptible to metabolic dysfunction-associated steatotic liver disease [8–10], hepatitis [11] and hepatocellular carcinoma [12].

GH is the primary proximal hormonal regulator of sex differences in hepatic gene expression. GH is secreted by somatotrophs within the anterior pituitary gland in a sex-dependent manner, giving rise to an intermittent (pulsatile) plasma GH pattern in males and a persistent (near continuous) plasma GH pattern in females, as seen in rats, mice and humans [1]. These sex-differential temporal patterns of pituitary GH release regulate liver transcription through the actions of the GH-responsive TF STAT5 [13] and its downstream target genes, most notably the TFs CUX2, BCL6 and ONECUT1/HNF6. These TFs work in concert with STAT5 to control sexually dimorphic hepatic gene expression patterns, as seen in mouse and rat models. In male liver, STAT5 is activated in a pulsatile manner by plasma GH pulse-induced STAT5 tyrosine phosphorylation, STAT5 dimerization and nuclear translocation followed by localized chromatin opening associated with STAT5 binding to liver chromatin and transcriptional activation of STAT5 target genes. These cellular responses to plasma GH pulse stimulation are overridden when male mice are given a continuous infusion of GH (cGH treatment), which substantially feminizes gene expression in the liver within 7-12 days [14]. cGH treatment also reverses the sex-dependent patterns of chromatin accessibility in male liver by closing chromatin at genomic sites that are more open (more accessible) in male as compared to female liver (male-biased DNase-I hypersensitive sites, DHS) while inducing chromatin opening at many female-biased DHS in male liver [15]. These chromatin changes are closely associated with repression of male-biased gene transcription and induction of many female-biased genes to female-like levels of expression [15].

Single-cell sequencing has dissected the cellular heterogeneity and zonation across the liver lobule at high resolution [16], increasing our understanding of the genes and pathways that define hepatic cell identity. scRNA-seq atlases reported for human liver [17] and mouse liver [18] have shown how transcriptional regulation contributes to cell-type specificity. Furthermore, single nucleus (sn)RNA-seq has advantages over single cell-seq, including the ability to isolate and then sequence RNA from nuclei representing all cells in a frozen tissue sample while maintaining cell viability and with increased sensitivity for detecting lncRNAs.

snATAC-seq (single nucleus assay for transposase-accessible chromatin using sequencing) is an extension of bulk ATAC-seq [19] and uses the transposase Tn5 to profile the chromatin accessibility patterns for thousands of individual nuclei [20, 21]. Integration of snRNA-seq with snATAC-seq identify cell subpopulations more accurately and give novel insight into gene regulation [22]. Traditionally, such multimodal analyses was done separately on different sub-samples from the heterogeneous population, measuring gene expression and chromatin accessibility [23], which then required computational algorithms to jointly analyze the two modalities. However, single cell 10x Genomics multiOmic technology [24] allows joint profiling of gene expression and chromatin accessibility on the same individual sets of cells, which constitutes a more robust approach for linking function between *cis*-regulatory elements and target genes.

Here, we obtained paired single-nucleus chromatin accessibility (snATAC-seq) and gene expression (snRNA-seq) profiles from 46,188 liver nuclei isolated from adult male, adult female, and continuous GH-infused male mice to comprehensively map the regulatory architecture controlling sex differences in hepatic gene expression. We developed a novel intronic-monoexonic UMI counting strategy to identify and correct for polyribosome-associated secretory protein RNA contamination that can affect liver snRNA-seq datasets. Further, we identified differential accessible chromatin regions (DARs) specific to each major liver cell type, as well as DARs whose accessibility differs between sexes and is regulated in a GH-dependent manner. Correlation-based linkage analysis was used to map sex-biased DARs to their putative sex-biased gene targets, and ATAC-seq footprinting indicated novel TF occupancy at sex-biased DARs linked to MASLD-enabling and MASLD-protective sex-biased genes. Further, by leveraging hypophysectomy response classifications for sex-biased genes [25, 26], we identified distinct TF motif signatures associated with GH-activated (class I) versus GH-repressed (class II) regulatory programs. Furthermore, as TF motifs can co-bind, co-operate and co-regulate gene function [27], we explored motif co-binding events at sex-biased DARs linked to sex-biased genes. Finally, we mapped the global landscape of intercellular signaling between liver cell types and identified sex-biased differences in intrahepatic cell-cell communication using CellChat [28]. These integrated analyses provide a comprehensive resource for understanding the transcriptional and epigenetic mechanisms controlling sex-dependent liver function and disease predisposition.

## Results

### Single nucleus transcriptional and chromatin accessibility profiling of mouse liver

Nuclei were extracted from adult male and adult female mouse liver, and from livers of adult male mice continuously infused with GH for 12 d (cGH treatment), which overrides the endogenous adult male pattern of pulsatile plasma GH stimulation of hepatocytes and largely feminizes gene expression within 1 week [29]. snATAC-seq and snRNA-seq 10X Genomics multiOmic sequencing libraries [24] were prepared from nuclei extracted from n=3-4 mice per group to obtain paired chromatin accessibility and gene expression data for thousands of individual nuclei for each biological condition. snRNA-seq data were collected using a custom mouse mm10 GTF reference file representing 76,011 genes, including 48,261 liver-expressed lncRNA genes, a majority of them novel [30].

Nuclei with high exonic to intronic snRNA-seq read ratios, indicative of cytoplasmic RNA contamination, were filtered out. Further, gene expression profiling was implemented using a novel intronic + mono-exonic read counting method, which excludes sequence reads derived from spliced (mature) transcripts and minimizes contamination of liver non-parenchymal cell (NPC) clusters by highly expressed hepatocyte secretory protein RNAs, such as *Albumin*. ATAC-seq peaks were identified using MACS2 [31], applied separately to each multiOmic sample and each annotated liver cell type. Batch correction and sample integration were performed using Harmony [32] for both the snRNA-seq and snATAC-seq modalities. snRNA-seq UMI counts and scATAC-seq cut sites per cell were processed using Seurat [33] and Signac, respectively. Filtering across both data modalities resulted in a total of 46,188 nuclei across all three biological conditions that passed our quality control thresholds for both transcriptomic and chromatin accessibility profiles (Fig. S1A, Table S1B). A combined UMAP based on both modalities (snATAC-seq and snRNA-seq) was developed using weighted nearest neighbor (WNN) analysis, an unsupervised framework for integrating multimodal single-cell data [13] (Fig. 1A). Five major liver cell types were identified by their marker gene expression patterns (Fig. S1B): hepatocytes, endothelial cells, hepatic stellate cells, immune cells and cholangiocytes. Three major hepatocyte zonation clusters were identified by their liver zonation marker genes: periportal, mid-lobular and pericentral hepatocytes. Nuclei counts for each cell type are shown in Table S1B.

**Fig. 1.**
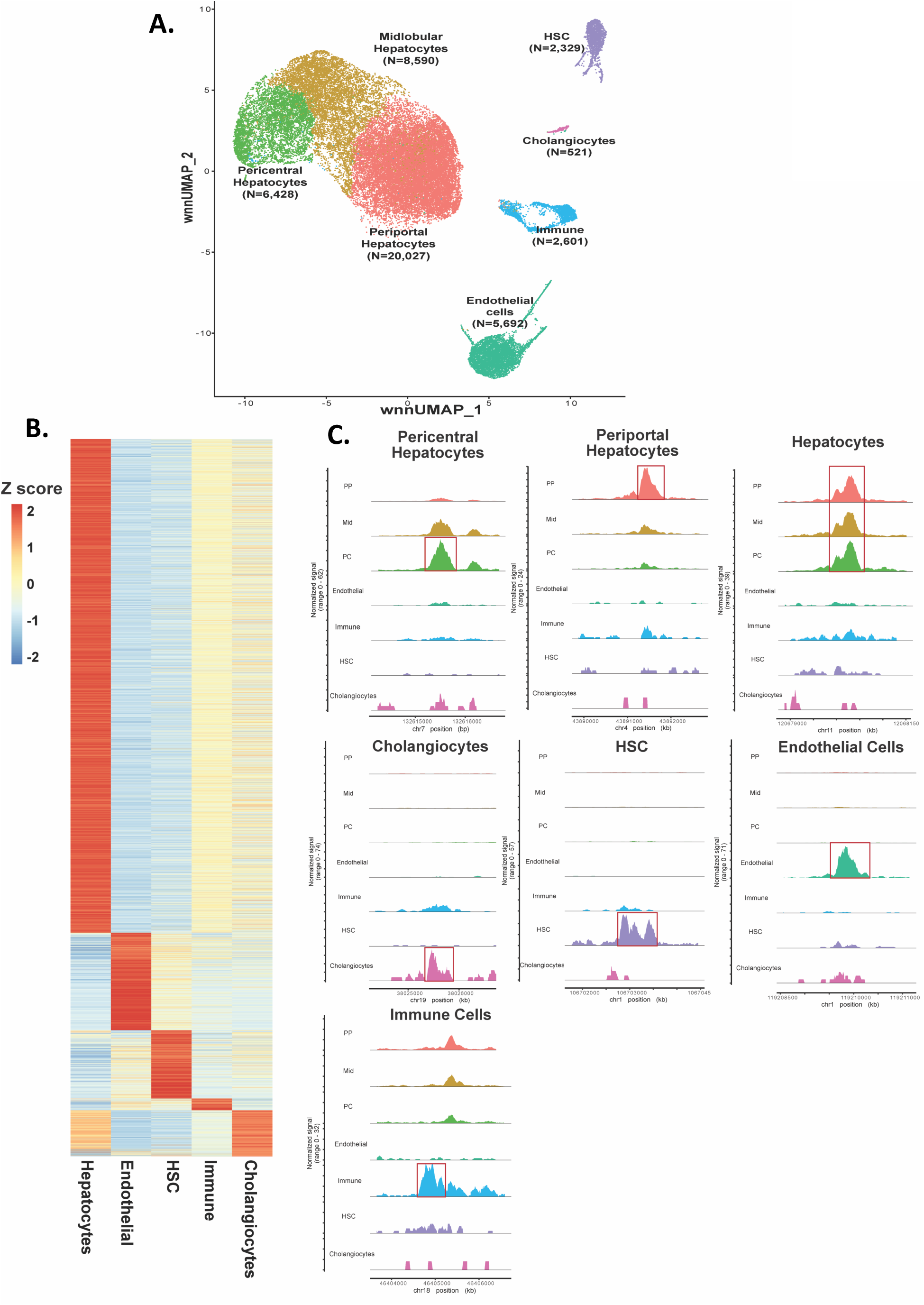
Single-cell transcriptional and chromatin accessibility profiling of mouse liver. **A.** UMAP of mouse liver cell types based on transcriptomic and chromatin accessibility profiles of 34,364 single nuclei obtained using 10X Genomics MultiOmic technology (snRNA-seq + ATAC-seq) integrated from four samples representing three biological conditions: male liver, female liver and cGH-treated male liver. Nuclei counts for each cell type are shown in parentheses. **B.** Scaled heatmap of 6,543 differentially accessible regions (DARs) identified across cell types (Table S2A). The average number of ATAC-seq cut sites within each DAR for each cell type is represented as a Z-score scaled by row. **C.** ATAC-seq genomic DNA fragment coverage surrounding several cell type-specific DAR peak regions. The track was generated using the coverageplot function in Signac.

### snATAC-seq peak characterization

ATAC peak regions were merged across all samples using Signac [34] to obtain 127,957 merged peaks (mean peak width: 549 ± 354 nt (SD); Table S1C, column BS). 74,860 (59%) of these ATAC regions overlapped accessible chromatin regions identified as DHS in nuclei extracted from bulk mouse liver tissue [35–37] and 53,097 (41%) were novel (Fig. S2A, Table S1D). Hepatocyte snATAC-seq activity (mean normalized ATAC signal intensity in hepatocyte cluster) was 4.6-5.3-fold higher at the 74,860 ATAC peaks that overlapped a liver DHS than at the no overlapping ATAC peaks (Table S1D); thus, weak hepatocyte chromatin accessibility may explain the absence of the latter ATAC sites in the published bulk liver tissue DHS peak sets, where hepatocytes are the dominant cell type. Consistent with this, snATAC peaks that did not overlap a bulk liver DHS showed a 4.33-fold higher maximal ATAC activity across the major NPC populations than in hepatocytes (Table S1D). Moreover, 83% of the non-bulk liver DHS overlapping snATAC peaks are maximally accessible in NPCs (Table S1E.1, column N). Thus, the snATAC-seq peak set includes many novel NPC chromatin accessible regions.

ChIP-seq signals for bulk liver indicate that 67% of the 54,923 snATAC peaks overlapping a DHS in [37] (Table S1C, Set III DHS) are enhancers (H3K4me1 marks), 18% have promoter marks (H3K4me3) and 15% are insulators (CTCF ChIP-seq signal; Fig. S2B). Further, based on chromatin state maps defined separately for adult male and female mouse liver [38], the largest subset of ATAC peaks are in an enhancer state (42-43%), followed by inactive states (21-22%), transcribed states (13-15%), promoter states (11%) and a bivalent state (8-10%) (Fig. S2C). Thus, the liver snATAC-seq dataset is robust and detects accessible chromatin with the expected histone marks and epigenetic patterns in mouse liver.

### Chromatin accessibility patterns define liver cell-type specificities and sex-biased chromatin responses

One goal of this work is to identify ATAC regions, and their associated regulatory sequences, enriched in each liver cell type. We identified a total of 6,543 such liver cell type-enriched ATAC regions (Fig. 1B, Table S2A), of which 683 regions passed a stringent threshold for liver cell type specificity (Fig. 1C, Table S1E.4). A second goal was to determine the liver cell type-specificity of regulatory regions that are sex-biased and GH-regulated.

Strikingly, sex differences in snATAC peaks were almost exclusively found in hepatocytes, with fewer than 2% of sex differences in ATAC regions found in liver NPCs (Table S2B). Further, by comparing ATAC peaks between male and female hepatocytes, and also between male hepatocytes and cGH-treated male hepatocytes, we identified 1,230 ATAC regions with greater chromatin opening in female compared to male liver (female-specific DARs, at FDR < 0.05) whose accessibility in male liver increased significantly following cGH infusion.

Similarly, we identified 3,518 ATAC regions significantly more open in male than female liver (male-specific DARs, at FDR < 0.05) whose accessibility was significantly decreased following cGH infusion (Fig. 2A, Fig. S3A (ATAC sets D and E; Table S2C). Importantly, 86-95% of these sex-biased DARs (1,062 of 1,230 female-specific DARs; 3,325 of 3,518 male-specific DARs) showed no significant difference in accessibility between female hepatocytes and cGH-treated male hepatocytes (Table S2C). Thus, cGH infusion in male liver is sufficient to feminize the sex-biased pattern hepatocyte chromatin accessibility at more than 4,300 genomic regions in the absence of other exogenous hormonal factors.

**Fig. 2.**
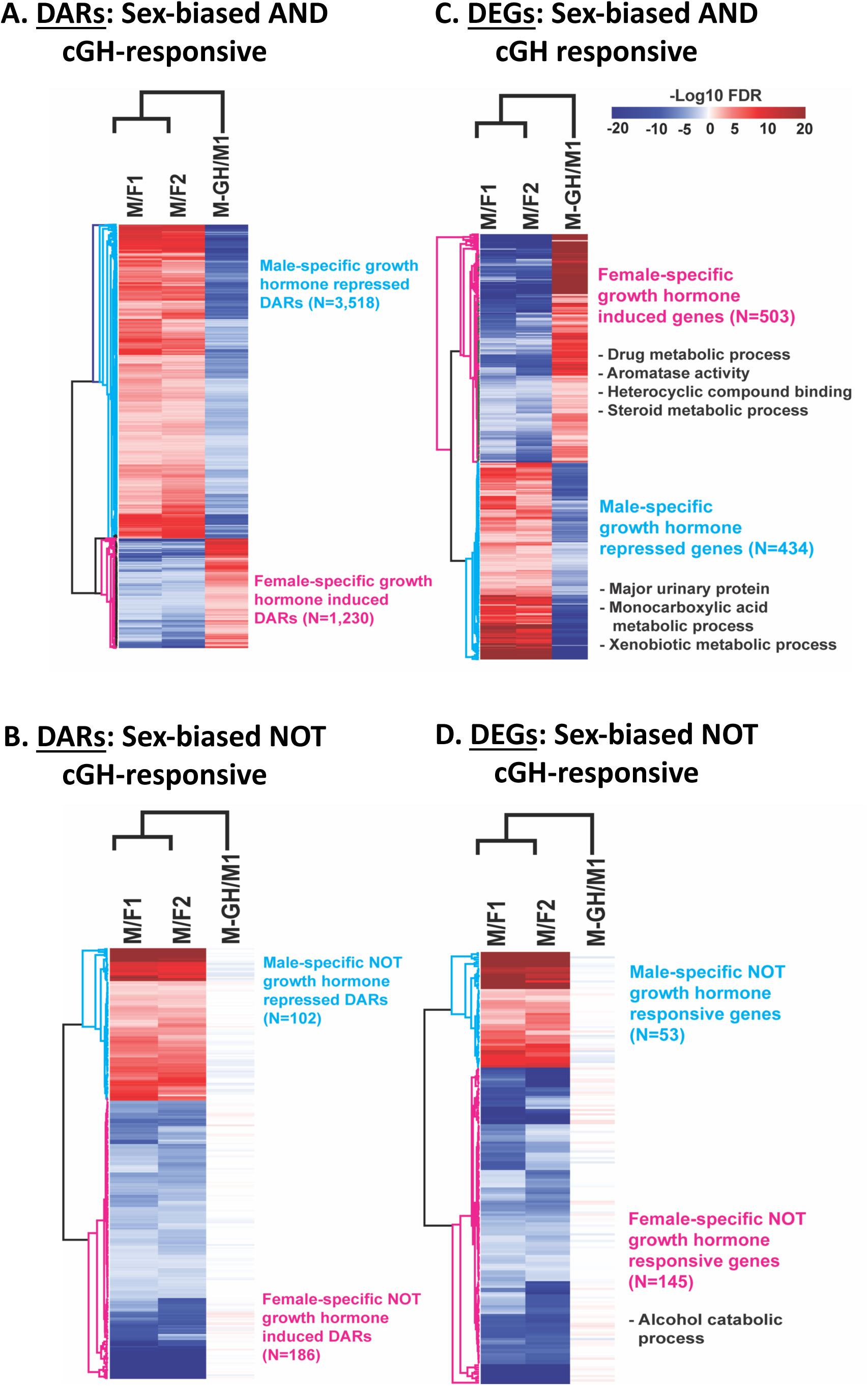
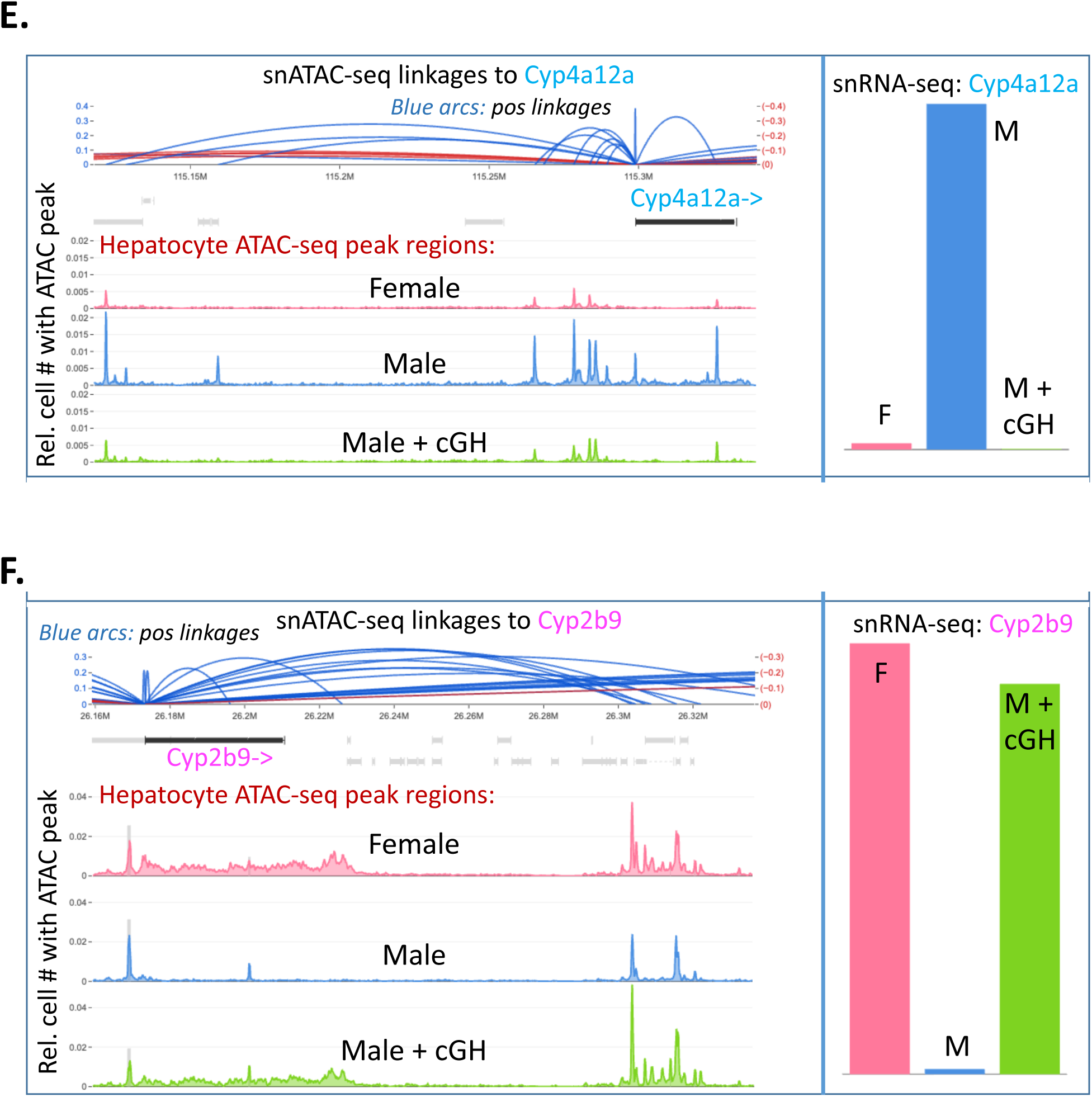
Sex-biased hepatocyte DARs and DEGs and their responsiveness to cGH treatment. **A.** Heatmap of hepatocyte DARs that are M-specific and cGH-repressed or are F-specific and cGH-induced. **C.** Heatmap of DEGs that are M-specific and cGH-repressed or are F-specific and cGH-induced in hepatocytes, with the top DAVID functional enrichment terms shown at the *right*. **B, D.** Corresponding heatmaps of M-specific DARs that are not cGH non-responsive and consistently show higher ATAC-seq activity in cGH-treated male than in female hepatocytes, or F-specific DARs that are not cGH non-responsive and consistently show lower ATAC-seq activity in cGH-treated male than in female hepatocytes. The color scale represents the significance of the differential chromatin accessibility (A, B) or of differential gene expression (C, D), with comparisons yielding a positive fold-change value shown in blue (male-biased) and those yielding a negative fold-change value shown in red (female-biased). Numbers of DARs and DEGs are as shown. **E, F.** Browser screen shots showing ATAC regions linked to the strongly male-biased Cyp4a12a (E), and to the strongly female-biased Cyp2b9 (F) in male, female, and cGH-infused male liver. Positive linkages are shown in blue and negative linkages in red, with the height of the arc from the ATAC site to the gene TSS indicating the extent of correlation (Y-axis). Gene expression levels are shown at the right.

We also identified a much smaller set comprised of 102 male-specific ATAC regions (Fig. S3, DAR set F) that showed no significant response to cGH infusion, and consequently, were more accessible in both untreated male and cGH-treated male hepatocytes than in female hepatocytes, i.e., chromatin accessibility was male-biased but was stringently non-responsive to cGH. 82 of these 102 male-specific but cGH-unresponsive DARs were autosomal and 20 were on ChrY. Similarly, we identified 186 female-specific DARs that were unresponsive to cGH infusion and consistently showed greater chromatin accessibility in female than untreated male or cGH-treated male hepatocytes, i.e., their accessibility was female-biased but was stringently unresponsive to cGH (Fig. 2B). 145 of these 186 DARs were autosomal and 41 were on ChrX.

We conclude that sex differences in liver chromatin accessibility are widespread, are almost exclusively localized to hepatocytes, and are highly responsive to GH treatment, with continuous GH infusion in male mice feminizing chromatin accessibility at thousands of ATAC regions.

### Sex differences in gene expression are primarily limited to hepatocytes

We used the snRNA-seq datasets to characterize sex differences in gene expression in each liver cell population. Consistent with our prior findings [39], sex differences in expression were largely limited to hepatocytes (Table S2B): 503 genes were female-specific and cGH-induced, and 434 genes were male-specific and cGH-repressed (Fig. 2C, Fig. S3B). This finding is in accord with our finding, above, that sex differences in chromatin accessibility are a characteristic of hepatocytes but not in other liver cell populations. The male-specific genes were enriched for functions related to monocarboxylic acid metabolic process and xenobiotic metabolism and the female-specific genes were enriched for drug metabolic process and aromatase activity (Table S2F). We also identified 145 female-specific genes and 53 male-specific genes that were stringently unresponsive to cGH infusion, based on the comparison between cGH-treated male hepatocytes and 2 independent female liver snRNA-seq samples (Fig. 2D, Fig. S3B, Table S2D). Thus, a subset of sex-biased genes is unresponsive to cGH in hepatocytes, indicating that the incomplete feminization described earlier for cGH-infused male liver using bulk liver RNA-seq [29] is not due to differences in sex-biased gene responses to cGH in hepatocytes versus NPCs.

### Peak-to-gene correlations link sex-biased DARs to their regulatory gene targets

We integrated the multiOmic data for chromatin accessibility and gene expression to associate snATAC peaks to specific genes through linkage analysis. This analysis is based on Pearson correlations between ATAC-seq peaks and genes across the population comprised of thousands of individual cells and gives a direct and interpretable measure of the linkage strength (values ranging from -1 to 1). Overall, we identified 226,841 ATAC-seq peak─gene linkages across the genome, including linkages to 12,891 liver-expressed lncRNAs whose expression patterns are captured by snRNA-seq [39] (Table S3B). Most (77%) of the peak-gene linkages showed a positive correlation, indicating positively acting enhancer activity. Individual ATAC peaks were linked to 3.9 genes (mean) ± 4.2 (SD) (Table S3A), and individual genes were linked to 9.4 ATAC-seq peaks (mean) ± 15.5 (SD) on average (Table S3B). ATAC-seq profiles, linkages to associated regulatory regions, and snRNA-seq-determined expression data in both male and female liver, and in livers of male mice given cGH infusion are shown for two highly sex-biased genes, *Cyp4a12a* (male-biased) and *Cyp2b9* (female-biased) (Fig. 2E, Fig. 2F). These data also illustrate how cGH feminizes ATAC-seq profiles in addition to gene expression, results in near-complete down regulation of *Cyp4a12a* and decreases in chromatin accessibility at each of the positively linked ATAC regions, with the opposite responses observed in the case of *Cyp2b9*.

We identified four major sex-biased DAR sets based on their linkage patterns (Fig. S5A.1): **Set D1**, male-specific, cGH-repressed DARs that were linked to male-specific, cGH-repressed DEGs (N=408); **DAR Set D2a**, male-specific, cGH-repressed DARs that were linked to non-sex-biased genes (N=650); **DAR Set E1**, female-specific, cGH-induced DARs that were linked to female-specific, cGH-induced DEGs (N=286); and **DAR Set E2a**, female-specific, cGH induced DARs that were linked non-sex-biased genes (N=278). We also analyzed the DAR-gene linkage data to identify sets of sex-biased DARs linked to a set of 205 sex-biased genes whose responses to hypox were established by bulk RNA-seq [26] (Table S2G). We considered the two major classes of sex-biased genes, designated hypox class I and hypox class II, which show a fundamental difference in how they respond to the loss of GH after surgical removal of the pituitary gland by hypophysectomy (hypox) [25, 26].

Class I sex-biased genes are activated by the plasma GH pattern of the sex where the gene is more highly expressed, whereas class II sex-biased genes are repressed by the plasma GH pattern of the sex where the gene shows lower expression [25, 40]. We identified n=71 male-biased and cGH-repressed hypox class I genes, which were linked to n=226 male-biased and cGH-repressed DARs (Set 1 genes), as well as n= 15 male-biased and cGH-repressed hypox class II genes linked to n=52 male-biased and cGH-repressed DARs (Set 2 genes).

Corresponding sets of class I and class II cGH-induced female biased genes linked to cGH-induced and female biased DARs (Set3 and Set 4 genes) were also identified (Fig. S4). These DAR sets were analyzed further as described below.

### Sex-biased DARs linked to sex-biased genes are preferentially located in the same TAD

We investigated the potential for sex-biased DARs to form *cis*-regulatory interactions, as indicated by co-localization of a DAR within the same topologically associated domain (TAD) [41] as its linked target gene(s). TADs are dynamic intrachromosomal chromatin loops most often anchored by oriented CTCF and cohesion complexes, which interact and thereby link distal chromatin regions, bringing linearly distant regulatory elements to their target genes in *cis*. TAD annotations were assigned to each ATAC-seq region and its linked genes using TAD definitions for mouse liver [37, 42]. Overall, 76-84% of sex-biased and cGH-responsive DARs in sets D1 and E1, which are linked to correspondingly sex-biased and cGH-responsive genes (Fig. S5A.1), mapped to the same TAD as their linked genes (Fig. S6A, Fig. S6D). In contrast, a majority of the sex-biased, cGH-responsive DARs linked to non-sex-biased genes (DAR sets D2a and E2a; Fig. S5A.1) or those involving other classes of sex-biased genes (DAR sets D4 and E4; Fig. S5) mapped to different TAD compartments than their linked genes (Fig. S6B-S6C, Fig. S6E-S6F). A majority (65-87%) of sex-biased, cGH-responsive DARs that were linked to sex-biased genes with a defined hypox response class (Fig. S4, Table S2G) were also within the same TAD compartment (Fig. S7). Across all DAR−gene sets, linkages to genes in other TADs crossed much longer distances (Fig. S6G) and were generally weaker (lower linkage correlation value) and therefore less reliable, than within-TAD linkages (Fig. S6H). The strongest linkages were seen for sex-biased and cGH-responsive DARs linked to correspondingly sex-biased, cGH-responsive genes (DAR set D1 and E1) localized within same TAD (Fig. S6H). Finally, a majority (60%) of non-sex-biased ATAC-seq regions linked to non-sex-biased genes mapped to different TADs and showed weaker mean interaction correlations over longer genomic distances than within-TAD linkages (Fig. S6I). We conclude that sex-biased DAR-gene linkages are at least in part constrained by the insulation imparted by dynamic TAD looped chromatin structures.

### Interactions between sex-biased DARs and sex-biased genes are enhancer-based

We used the signage of the ATAC-gene linkage correlations to classify the interacting ATAC regions as either stimulatory, enhancer-based (positive correlations) or repressive (negative correlations). Almost all interactions (>97%) involving sex-biased, cGH-responsive DARs and target genes of the same sex bias and cGH responsiveness were identified as enhancer based (Fig. S8A, S8C). In contrast, only 58% of the much larger set of 42,206 non-sex-biased DARs linked to non-sex-biased genes formed exclusively positively correlated linkages; 33% of these DARs were involved in both positively and negatively correlated linkages with their target genes and 9% made exclusively negatively correlated gene linkages (Fig. S8G). Many negatively correlated (repressor-like) interactions were found for sex-biased, GH-responsive DARs linked exclusively to non-sex-biased genes (sets D2a, E2a) and for sex-biased DARs linked to sex-biased genes of the opposite sex bias, in addition to non-sex-biased genes (sets D4.1/D4.2 = D4a, E4.1/E4.2 = E4a) (Fig. S8B, S8D-S8F; Table S3A, column BW). This indicates a role for these sex-biased DARs in repression of the linked target genes in the sex where the DAR adopts a more open chromatin structure.

### Chromatin state distributions of sex-biased DARs

Genomic regions characterized by sex differential chromatin states are closely linked to sex-biased gene expression, as shown using chromatin state maps developed for both male and female mouse liver [38]. These maps, which are defined for sequential 200 bp genomic segments across the mouse genome, were used to annotate the chromatin states of each sex-biased DAR set. Chromatin state distributions in male mouse liver were very similar for male-biased DAR sets D1 (ATAC sites mapping to male-biased, cGH-repressed genes) and D2a (ATAC sites not mapping to any sex-biased genes), with enhancer state E6 comprising the largest subgroup (47-49%) of each DAR set (Fig. S11C, Fig. S11D; Table 5 of *Percentages* Excel file). State E6 has a high probability for open chromatin (DHS) associated with two active enhancer marks (H3K4me3, H3K27ac) and is relatively deficient in active promoter marks (H3K4me3) (Fig. S2D). Thus, DAR sets D1 and D2a are primarily comprised of enhancers and not promoter regulatory elements. By contrast, the male-biased, cGH-repressed ATAC regions in DAR set D3, which were not linked to any genes, showed a much lower frequency of the DHS-containing enhancer state E6 (19%) and of promoter state E7 (0.6% in D3 vs 8.6% in D1). Set D3 DARs also showed increased frequencies of enhancer state E11 (H3K4me1 marks only; 18.5% in D3 vs 6.4% in D1), bivalent state E12 (H3K4me1 mark + repressive chromatin mark H3K27me3) (17.8% in D3 vs. 3.9% in D1) and inactive state E2 (10.8% in D3 vs. 3.4% in D1) (Fig. S12C).

The relative absence of the DHS-containing chromatin states in DAR set D3 indicates that liver chromatin is generally less open than the male-biased, gene-linked DAR sets D1 and D2a, which when combined with the lack of H3K27ac enhancer marks and/or the presence of H3K27me3 repressive marks, likely accounts for the absence of gene linkages for set D3 DARs. Indeed, ATAC-seq intensity was significantly lower for DAR set D3 than for DAR sets D1 and D2a (Fig. S5C; also see Table 2 of *Percentages* Excel sheet). Set D3 DARs also showed a significantly lower frequency of nearby expressed genes, greater distance to the nearest hepatocyte-expressed gene TSS (transcription start site), which likely is an additional factor in the absence of gene linkages compared to set D1 and set D2a DARs (Fig. S5D, Table S3D), given that the ATAC-seq peak-to-gene linkage algorithm considers linkages up to a distance of 1000 kb. In female livers, the genomic regions encompassing all three male-biased, cGH-repressed DAR sets (D1, D2a, D3) showed a decreased frequency of enhancer state E6 and promoter state E7, and an increased frequency of the inactive state E2 and the transcribed state E13 (Fig. S11C, Fig. S11D, Fig. S12C), as was expected.

We examined the chromatin states in female liver of female-biased, cGH-induced DARs: set E1 (linkages to correspondingly regulated female-biased genes), set E2a (linkages to non-sex-biased genes only), and set E3 (not linked to any genes). Similar to the trends seen with male-biased DARs, the frequency of enhancer state E6 and promoter state E7 decreased in going from female-biased DAR set E1 to E2a to E3 (state E6: 49%, 40%, 29%; state E7: 14%, 2.2%, 1.6%), while there were increases in the frequency of enhancer state E11 (H3K4me1 only) (E1: 10%, E2a: 23%, E3: 29%), bivalent state E12 (E1: 6%, E2a: 13%, E3: 14%) and inactive state E2 (E1: 1.1%, E2a: 1.8%, E3: 5.3%). The absence of gene linkages for female-biased DAR set E3 can therefore be understood in terms of chromatin states that are less frequently open and less active, as indicated by histone mark patterns, when compared to DAR set E1. Consistently, ATAC-seq intensity was significantly lower for DAR set E3 than set E1 (Fig. S5B, Fig. S5C); further, distance to the nearest hepatocyte-expressed gene was significantly greater for DAR set E3 (Fig. S5D), indicating a role for both factors in the absence of ATAC peak-gene linkages for set E3 DARs. Finally, the 3 female-biased DAR sets generally showed decreased frequency in male liver of active chromatin states (enhancer states E6 and E11, promoter state E7) and an increased frequency of inactive states (E1 and E2) and transcribed state E13, as expected (Fig. S11E, Fig. S11F, Fig. S12E). We conclude that the distinct gene linkage profiles of each sex-biased DAR set can in large part be explained by their chromatin state differences.

### Functional characterization of sex-biased DARs

Gene sets linked to each major sex-biased DAR set were input to Metascape [43] for functional enrichment analysis with cross comparisons (Fig. S13, Table *Metascape*). DAR set D1 target genes (male-biased) were enriched for functions related to metabolism of lipids, steroids and amino acids, amongst others, while DAR set E1 target gene (female-biased) enrichments included negative regulation of fatty acid oxidation, JAK-STAT signaling and biomineralization. Enrichments common between DAR set D1 and DAR set E1 targets included drug metabolism/cytochrome P450, chemical carcinogenesis and metabolism of lipids, consistent with the widespread sex differences in these liver metabolic pathways seen in both sexes. DAR set D2a (non-sex-specific target genes) was specifically enriched in diverse pathways and functions, including cell migration, proliferation, cell cycle and inflammatory response, while DAR set E2a shared many pathways with set D2a, but was also specifically enriched for other functions, including iron ion transport, choline catabolism and alcoholic liver disease. Importantly, the finding of many significant functional enrichments for the linked gene targets of DAR sets D2a and E2a gives strong supporting evidence that these linkages are specific and not random correlative associations.

### TF motifs associated with liver cell type-specific open chromatin regions

We used two complementary approaches to identify TF motifs associated with specific liver cell types and likely to be involved in cell type-specific regulation. First, we used chromVAR [44] to calculate chromatin accessibility activity values for each TF motif in each cell and then compared the overall accessibility of the motif in each liver cell type to its average accessibility across all liver cells. The results are expressed as per cell Z-scores (Fig. 3A, top, feature plot examples) or as average Z-scores (Fig. 3A, bottom) for each TF motif; further, log2 fold-change and associated FDR values quantify the extent of differential motif activity for each liver cell type-specific motif (Table S4B). Second, we performed motif enrichment analysis to identify TF motifs whose sequences are statistically overrepresented in each set of stringent cell type-specific DARs described above (683 DARs across 5 liver cell types; Table S1E.4) when compared to cell type-invariant background genomic regions of similar ATAC intensity (Table S4B.5).

**Fig. 3.**
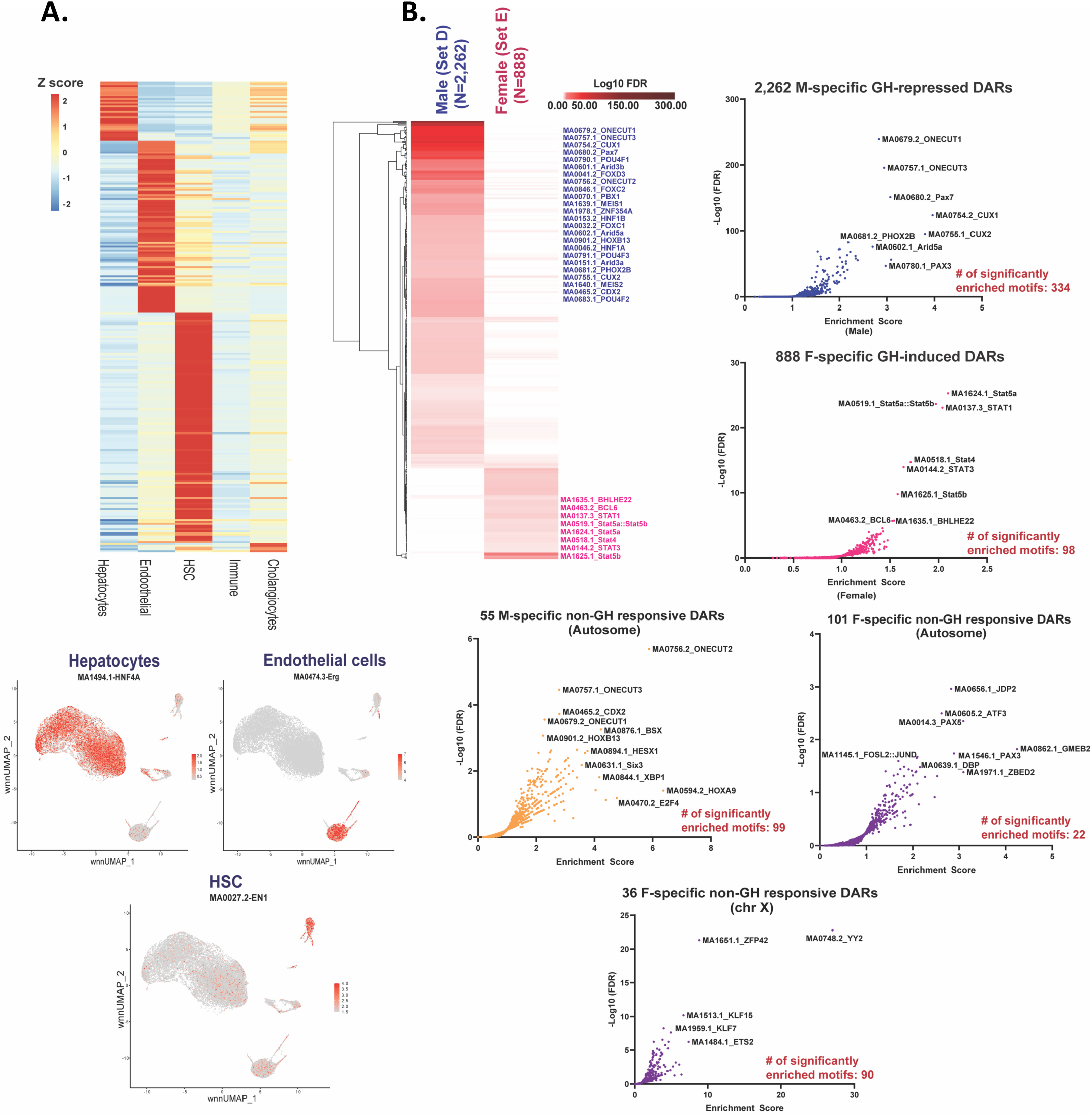
Liver cell type-specific and hepatocyte sex-biased motifs. **A. (Top)** Heatmap of average chromVAR motif activity (see Methods) for each cell type. The figure shows 255 cell type-specific motifs identified in Table 4A. The color scale represents a z-score scaled by row. **(Bottom)** Feature plot displaying motif activities in each nucleus for three cell type-specific motifs. **B. (Left)** 425 motifs enriched in sex-biased DARs (Table S4B, columns Z and AA). Background was set as the ATAC regions that were linked to non-sex-biased and non-GH-responsive genes (Set B1 in Fig. S5; see Table S3A summary). Listed at the right of the heatmap are top motifs (lowest - log10 FDR) for each sex. The color scale represents -log10 FDR. **(Right, Bottom)** Volcano plots of all 841 motifs from JASPAR database, with the number of significantly enriched motifs (enrichment score >1 and FDR <0.05) in each sex-biased DAR set indicated in each panel (see Table S4B). Male-specific and Female-specific DARs that did not response to cGH treatment (M:102 DARs, F: 186 DARs) are grouped separately, based on their location on sex chromosomes vs. autosomes.

ChromVAR analysis identified 255 motifs (40 motif clusters; see Methods) whose accessibility showed significant liver cell type-specificity. A majority of these motifs (222 motifs in 37 motif clusters) were specific for NPC clusters (Fig. 3A, Table S4B), likely due to the reduced sensitivity for discovery of motifs enriched in hepatocytes, which comprised 76% of cells in the overall cell population analyzed (Table S1B). We focused on 151 motifs (18 motif clusters) whose ChromVAR-identified liver cell type-specific motif activity was supported by a significant over representation of the motif in DARs specific to the corresponding liver cell cluster (Table S4B.4). Top hepatocyte-enriched motifs included HNF4A [45] and other TFs important for liver development and hepatocyte function, including HNF1B, ONECUT1 and ONECUT2 [46], various Nuclear Receptor family members with roles in hepatocyte metabolism, including PPARA [47, 48] and NR1H2/LXRβ [49], and TCF7L2, which is a key regulator of hepatocyte zonation [50]. Top motifs in liver endothelial cells included FOXO/FOXI/FOXJ family and various ETS family members (ELF, ELK, ETV, ERF, FLI, EHF, ERG subfamilies). FOXO1 is a regulator of vascular growth that couples metabolic and proliferative activities in endothelial cells [51] and ETS family TFs have important roles in vasculogenesis and angiogenesis [52], including ETV and ERG TFs, which regulate endothelial cell migration and angiogenesis [53, 54]. Top endothelial cell-enriched motifs included ETS−FOX heterodimeric motifs, such as ETV5:FOXI1 and ETV5:FOXO1. ETV:FOX, which are strongly associated with endothelial cell-expressed genes [55]. Top motifs in hepatic stellate cells, a mesenchymal cell population, included: FOXF1, which inhibits hepatic stellate cell pro-fibrotic gene activation during liver injury [56]; FOXK2, which is required for epithelial-mesenchymal transition in hepatocellular carcinoma [57]; PRRX1 and PRRX2, which regulate mesenchymal cell fate [58]; and EN1, which can promote epithelial-mesenchymal transition [59] (Table S4B.4).

### Motifs associated with sex-biased gene expression in hepatocytes

We sought to identify motifs for novel TF regulators of sex-biased gene expression based on their overrepresentation within genomic regions showing sex-biased and cGH-responsive ATAC activity in hepatocytes. We investigated 6 such DAR sets (Fig. S3A): male-specific and cGH-repressed (Set D); female-specific and cGH-induced (Set E); male-specific and cGH-unresponsive and located on ChrY (Set F_chrY) or located on autosomes (Set F_Auto); and female-specific and cGH-unresponsive and located on ChrX (Set G_chrX) or located on autosomes (Set G_Auto). We first filtered out ATAC peaks showing low activity in hepatocytes (bottom 30% of ATAC-seq regions ordered by normalized hepatocyte ATAC-seq signal intensity; Table S1C, column R), which removed only 4% of sex-biased DARs. We also removed ATAC sites without enhancer state designations. ATAC-seq regions that were in an enhancer state and whose ATAC activity did not show sex bias or respond to cGH treatment and were linked exclusively to non-sex-biased genes were used as a background set for the motif enrichment analysis (subset of ATAC-seq set B1 after removing regions with low hepatocyte ATAC activity; Fig. S5B, Table S1C, column O).

We identified 334 motifs (comprising 57 motif clusters) that were enriched at FDR < 0.05 at male-biased, cGH-repressed enhancer state DARs (DAR set D) and 98 motifs (36 clusters) enriched at female-biased, cGH-induced enhancer state DARs (DAR set E) (Fig. 3B, Table S4C). The top male-specific, cGH-repressed DAR motifs included ONECUT1 (HNF6A) and ONECUT3, PAX7, CUX1 and CUX2, PHOX2B and ARID5A. Few differences in the sets of top enriched motifs were found when we examined the DAR set D subsets comprised of DARs linked to male-biased, cGH-repressed genes (DAR set D1), as well as those exclusively linked to non-sex-biased genes (set D2a) and also for those not linked to any genes at all (DAR set D3), albeit with some differences in relative rankings of the enrichments (e.g., lower ranking of CUX2 and PHOX2B motifs in DAR set D1; Table S4C). In contrast, the female-biased, cGH-induced DAR set E and subset E1 (linked to female-biased, cGH-induced genes) showed top enrichments for STAT5 and other STAT motifs, as well as motifs for BCL6 and HAND2. The female-biased, cGH-induced DAR subsets linked to non-sex-biased genes (set E2a) or not linked to any genes (set E3) also showed the greatest enrichment for STAT5, but exhibited unique enrichments to other motifs, notably TFAP4, SNAI2, ZNF692 and MSC for DAR set E2a; and NEUROD2, OLIG2, NHLH1 and THAP1 for DAR set E3 (Table S4D).

Male-biased autosomal DARs that were unresponsive to cGH (set F_Auto; N= 55 DARs in enhancer state, out of 82) were enriched for many unique motifs (Table S4C), including 4 motifs from motif cluster 3 (BSX, RAX2, HESX1, SHOX). The cGH-unresponsive female-biased autosomal DARs (set G_Auto) were also enriched for many novel motifs (e.g., JDP2, ATF3, PAX5, GMEB2), as were the X-linked cGH-unresponsive female-biased DARs (set G_chrX), whose top enriched motifs include YY2, ZFP42 (REX1), KLF15 and ETS2 (Fig. 3B, Table S4D). YY2 shares high homology with two TFs involved in X-chromosome inactivation, YY1 and REX1/ZFP42 [60, 61]; KLF15 regulates hepatic gluconeogenesis and metformin action [62]; and ETS2 is a regulator of the mouse and human hepatic stellate cell lineage [63]. Thus, our multimodal analysis demonstrates sex-biased differences at the level of chromatin accessibility and identifies novel TFs that likely contribute to these differences.

### Motifs enriched in sex-biased DARs linked to hypox class-defined sex-biased genes

We searched for TF motifs enriched in DARs linked to sex-biased genes whose hypox response class has been defined (DAR sets 1-4; Fig. S4). We identified 364 motifs grouped into 73 clusters that were significantly enriched (FDR < 0.05) in at least one of the four DAR sets (Table S4E); a Z-score heatmap of their -log10 FDR values is shown in Fig. S14B (Table S4F). 168 of these motifs were uniquely enriched in Set 1 hypox DARs, with the top enriched motifs being NR5A2, PBX2, HLF and VAX1. NR5A2 is a critical regulator of hepatic glucose metabolism [64], PBX2 represses activation of the *Ugt2b17* promoter in liver cells [65], and HLF is an activator of hepatic stellate cells during liver fibrosis [66]. The top Set 2-specific enriched motifs (n= 19) were LHX4, ZNF8, ALX4 and SOX6. LHX4 may act as a tumor suppressor in hepatocarcinogenesis [67], and ALX4 inhibits proliferation, invasion and epithelial–mesenchymal transition in hepatocellular carcinoma [68]. Hypox-responsive DAR set 3 was specifically enriched in 28 motifs, including CEBPG, MEIS1, TEAD1, ESR1, TBX4 and STAT4. CEBPG is a pivotal regulator of liver nutrient metabolism and its control by hormones, acute-phase response and liver regeneration [69]. TEAD TFs regulate the Hippo signaling pathway in liver [70], ESR1 mediates liver responses to estrogens [71] and TBX4 regulates myofibroblast accumulation [72]. Finally, activation of STAT4 is associated with several mouse models of liver injury and with chronic liver diseases in humans [73]. 13 motifs were specifically enriched in hypox-responsive set 4 DARs, including NR3C1, NR3C2, SREBF2, HOXA4 and HES6. NR3C1 (glucocorticoid receptor) regulates gluconeogenesis in the liver [74, 75], SREBF2 is a master regulator of sterol and fatty acid synthesis [76], and HES6 represses HNF4α-activated PPARγ2 transcription, which decreases hepatic fat accumulation in obese mice [77]. Thus, we discovered several top enriched motifs with well-defined functions in the liver but whose roles in liver sex bias is still unknown and will require further experimental investigation.

### Sex-biased MASLD-linked genes regulated by plasma GH patterns

The sex bias of MASLD and its downstream pathologies, including inflammation, fibrosis and hepatocellular carcinoma, is driven by sex differences in hepatic GH signaling, which regulates the sex-dependent transcription of many MASLD susceptibility genes[10]. We performed an in-depth literature analysis to identify **s**ex-biased genes whose function impacts MASLD and/or associated liver pathologies, either by contributing to, protecting from disease development or severity, including genes whose impact on liver disease is more complex and context-dependent (Fig .4A). Key MASLD-enabling male-biased genes include *Bcl6*, which enhances MASLD progression by repressing PPARA-dependent hepatoprotective fatty acid oxidation genes [78, 79] and promotes hepatocellular by an immunosuppressive mechanism [80]) and *Nox4*, which prevents oxidative damage in early stages of disease [81] but become maladaptive and mediates oxidative stress, inflammation, and hepatic fibrosis as disease advances [82]. Female-biased genes that are MASLD-protective include: Trim24, which is an epigenetic co-regulator of transcription that represses steatosis, inflammation and fibrosis in mice [83, 84], but whose overexpression is linked to poor outcomes in human hepatocellular carcinoma [85]; Fmo2, which inhibits SREBP1 translocation, blocking steatosis [86]); and Hao2, a metabolism-biased tumor suppressor in hepatocellular carcinoma [87, 88].

The effects of cGH infusion in male mouse liver [29] and hepatocyte-specific STAT5 loss [89] reveal that these MASLD-relevant genes are primarily regulated by GH, in many cases via STAT5 signaling (Fig. 4A). In contrast, the estrogen surge in diestrus has no effect on expression of most of these genes (GEO dataset GSE131172). Further, the absence of a direct effect of androgen is indicated by the lack of response to liver androgen receptor (AR) knockout, which contrasts dramatically with the major effects in male liver of global AR knockout, and of castration and testosterone replacement (GEO dataset GSE147609). The latter responses are consistent with the role of androgen in setting hypothalamic regulation of pituitary GH secretory patterns.

**Fig. 4.**
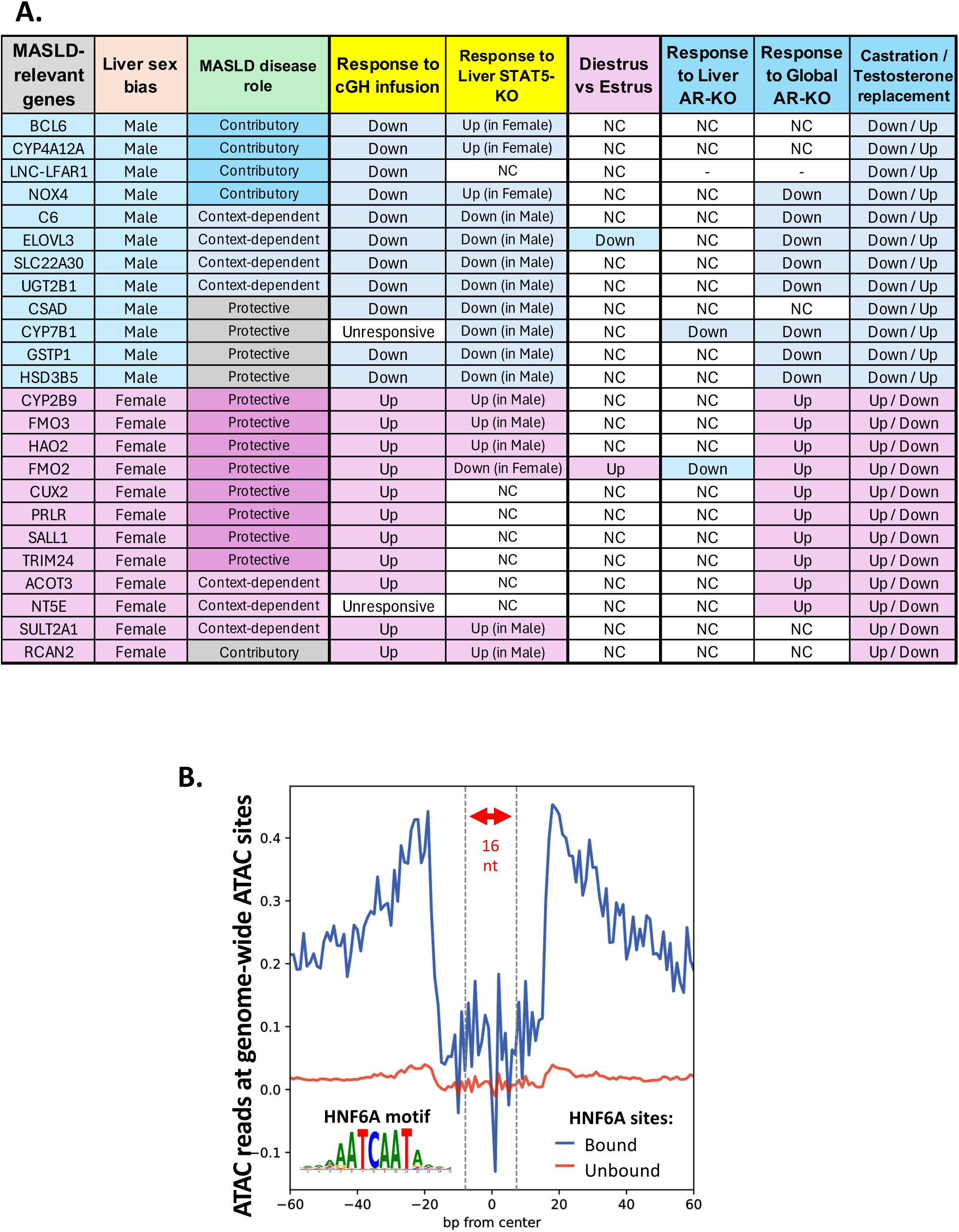
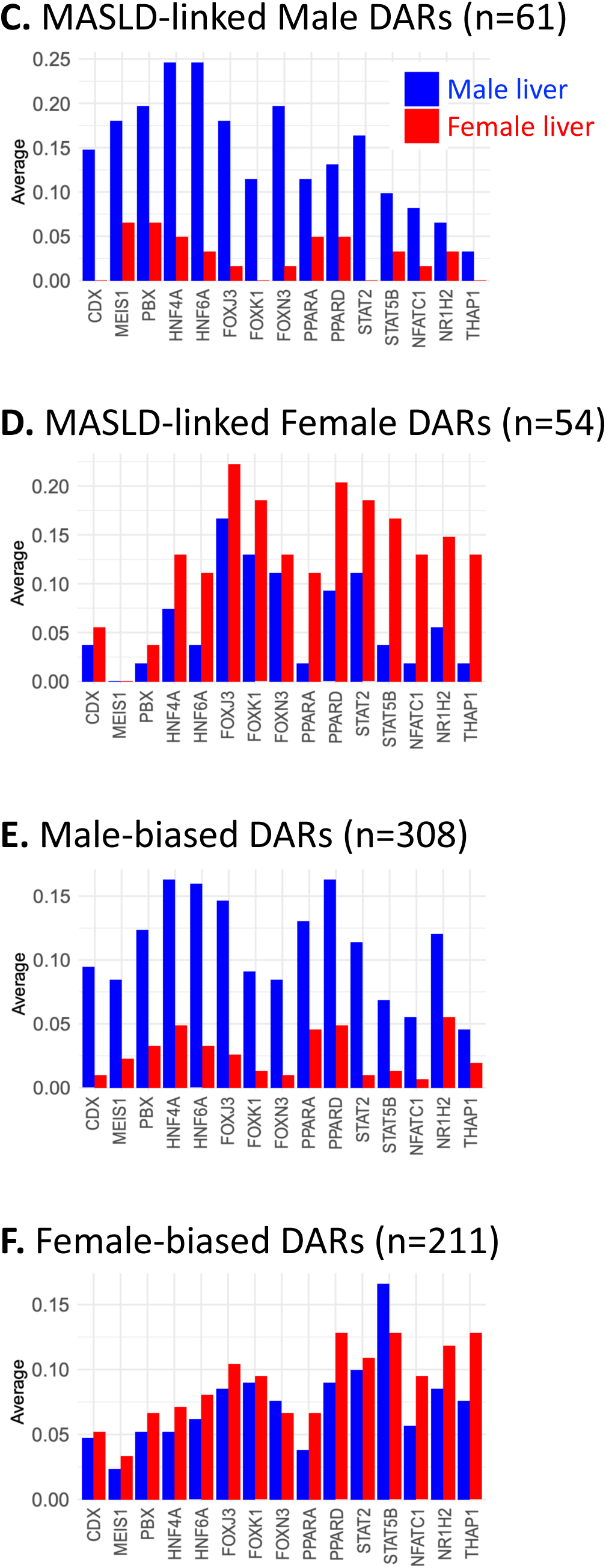
MASLD-relevant genes (A), and ATAC-seq footprinting analysis of motif occupancy at the full sets of sex-biased enhancer DARs, or at the DAR subsets linked to MASLD-relevant genes (B, C). **A.** Liver sex-biased genes whose function impacts MASLD and/or associated liver pathologies, by either contributing to, protecting from, or exerting effects that are more complex and context-dependent, as indicated. Also shown is a summary of published data indicating that plasma GH secretory patterns, and not androgen or estrogen is the major hormonal regulator of their liver sex-bias (see text). **B.** Example of ATAC-seq footprint identified by TOBIAS for Hnf6a at motifs predicted to be Hnf6a bound vs unbound in 8 wk male mouse liver bulk liver ATAC-seq data, aggregated across all 94,000 liver ATAC regions identified in that ATAC-seq sample and centered at the Hnf6a motif. C-F. Motifs identified as *bound* in footprint analysis by TOBIAS in bulk liver ATAC-seq data from 8 wk old male and female mice, analyzed separately for each sex. Motifs were included in the output if they were annotated as bound at a DAR in at least 10% of any of the 4 sets of sex-biased DARs that were analyzed: MASLD gene-linked male-biased and female-biased enhancer DARs, and the full sets of male-biased and female-biased enhancer DARs D1 or E1 in the bulk ATAC-seq data from at least one sex. Y-axis: average motif frequency, calculated across the each of the above four DAR sets.

### Motif occupancy determined by ATAC-seq footprint analysis

We identified 61 male-biased, cGH-repressed set D1 enhancer DARs and 54 female-biased, cGH-induced set E1 enhancer DARs linked to the above sets of male-biased and female-biased MASLD-relevant sex-biased genes using the peak-to-gene linkage analysis data in (Table S3B). Next, we used TOBIAS [90] to predict transcription factor binding to these sets of MASLD gene-linked DARs through footprinting analysis, and to compare results to the full sets of sex-biased enhancer DARs (set D1, set E1). TOBIAS scans for motifs for each TF and calculates a footprinting score that integrates both ATAC-seq read depletion and accessibility signals and is used to call individual motifs as ‘bound’ or ‘unbound’ by the TF of interest, after correcting for the intrinsic Tn5 transposase cut site bias based on sequence composition using a dinucleotide weight matrix [90]. Fig. 4C and 4D show the relative frequencies with which motifs for individual liver-expressed TFs of interest are predicted to be occupied in mouse liver chromatin from male (blue bars) and female mouse liver (red bars). Many motifs showed a higher frequency of TF occupancy at the set of 61 male-biased MASLD gene-linked enhancer DARs in male than in female liver, as expected due to the male bias in chromatin accessibility at these DARs (Fig. 4C). Similarly, a female-biased pattern of TF occupancy was seen at the set of 54 female-biased MASLD gene-linked enhancer DARs (Fig. 4D). However, several TFs showed distinctly different occupancy patterns between the two sets of MASLD gene-linked sex-biased DARs. Notably, motifs occupancy for Cdx, Meis1 and Pbx factors showed several-fold greater frequency at the set of male-biased MASLD gene-linked DARs, while Nfatc1, Nr1h2 (LXR-β) and Thap1 showed several-fold higher frequency at the set of female-biased MASLD gene-linked DARs, suggesting these factors play important roles in the expression of MASLD-relevant genes in each sex. Motif occupancy profiles for the full sets of sex biased gene-linked male-biased and female-biased enhancer DARs are shown in Figs. 4E and 4F for comparison. The liver-expressed CDX family factor Cdx4 shows female-biased expression [91], suggesting it may act to repress male-biased genes in female liver. Meis and Pbx factors form stabilizing heterodimers [92–94], and both factors contribute to male-biased liver disease: Meis factors promote tumor progression in human hepatocellular carcinoma [95] and Pbx1 contributes to liver fibrosis in high fat diet mouse model [96]. Regarding the TF family members showing greater occupancy at MASLD gene-linked female-biased DARs: Nfatc1 plays a role in hepatocellular carcinoma progression [97]; Nr1h2 (LXR-β) is a master lipogenic TF that promotes lipogenesis [98], and the related Nr1h1 (LXR-α) shows sexually dimorphic functions in liver metabolism [99]; and Thap3 is positively associated with oxidative phosphorylation gene expression in liver cancer, providing a metabolic advantage to cancer cells [100].

### Motif co-occurrence analysis

Motifs and their bound TFs can function in isolation but in many instances, TFs co-bind, co-operate and co-regulate gene function, which enables a TF bound at one motif to impact the activity of a TF bound at a nearby motif [28]. We analyzed motif co-occurrence events in the sets of sex-biased enhancer state DARs linked to sex-biased genes (DAR sets D1 and E1) and in the hypox response class-defined sex-biased enhancer state DARs (DAR sets 1-4) by hierarchical clustering, where the presence of a motif in each DAR peak is indicated in Fig. 5 (rows represent DARs, columns display motifs at each DAR; full details in Table S5A). Co-occurring motifs most often belonged to the same motif family, e.g., as seen for FOX family motifs, which co-occurred in both male-biased and female-biased DARs (Fig. 5A, orange rectangle; RSAT motif family cluster 10). However, these FOX family motifs co-occurred within a larger cluster of motifs (cluster 4) comprised of motifs mapping to other, distinct RSAT motif family clusters. We also identified motif clusters whose co-occurrence was sex-dependent, for example, clusters 3 motifs displayed stronger patterns of co-occurrence at male-biased DARs than at female-biased DARs (blue rectangle shown on heatmap; Fig. 5A). This difference is also indicated by the greater total number of motifs from cluster 3 in DAR Set D1 (n= 2,272) than in DAR Set E1 (n = 748) (Table S5A). These cluster are comprised of homeodomain factors (blue highlight in Fig. 5A). Cluster 12, consisting of STAT family TFs and the TF Meis1, showed more extensive associations with the set of female-biased DARs.

**Fig. 5.**
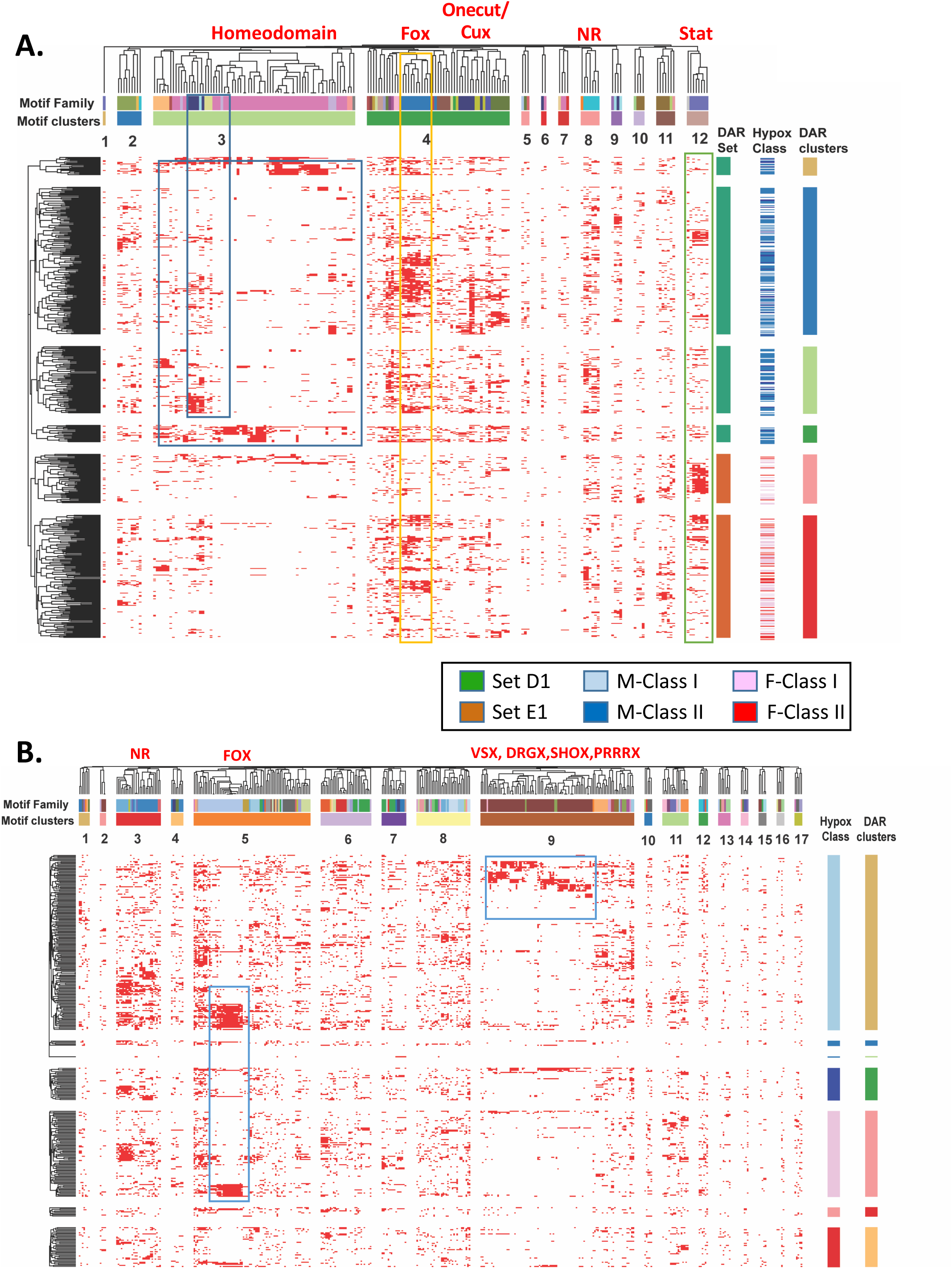
Motif co-occurrence at sex-biased DARs (A) and at hypox class-defined sex-biased DARs (B). **A.** Heatmap showing motif co-occurrence at sex-biased enhancer state DARs linked to sex-biased genes and that contain at least one motif, after removing low activity DARs (DAR sets D1 (N=305) and E1 (N=210; see Fig. S5). Columns represent motifs enriched in DAR sets D1 and E1 (see Table S4D); rows represent DARs. Hierarchical clustering was performed on the DAR─motif matrix where each entry is a binary value indicating the presence/absence of the motif in that DAR peak. The motif clusters and the DAR clusters shown were obtained by branch cutting of the dendrogram. The column annotation indicates motif families identified by the RSAT clustering tool. Row clustering was supervised under conditions that allowed the DARs to cluster within their respective DAR sets. Row annotations (at *right*) identify DAR sets by sex bias (D1, E1) and by hypox class, and DAR clusters defined by the dendrogram. **B.** Heatmap showing motif co-occurrence at DARs linked to sex-biased genes with defined hypox class (DAR sets 1, 2, 3 and 4; Fig. S4). Columns represent enriched motifs in sets 1-4 from Table S4E. Rows represent enhancer DARs from sets 1-4 that contain at least one motif, after removing low activity DARs. Hierarchical clustering and data presentation was as described in Fig. 5. The column annotation also shows motif clusters identified by the RSAT clustering tool. The row annotations show DAR clusters and hypox class sex-biased DAR sets.

We created a non-redundant motif co-occurrence heatmap by displaying a single representative motif for each motif family. Motif clusters 1, 2 and 3 showed stronger patterns of co-occurrence at male-biased DARs (Fig. S15A, blue highlight), while motif clusters 8 and 9 showed stronger patterns of co-occurrence at female-biased DARs (green highlight).

We also clustered the sets of sex-biased DARs linked to hypox class-defined sex-biased genes and their enriched motifs (Fig. 5B, Table S5E). Of note, cluster 9 motifs co-occurred specifically in the set of sex-biased DARs linked to male class I sex-biased genes. These motifs included the TFs VSX, DRGX, SHOX and PRRRX. TFs represented by motif cluster 5, which included many FOX family factors, showed strong patterns of co-occurrence for sex-biased DARs linked to class I male-biased genes and also class I female-biased genes. The corresponding non-redundant motif family co-occurrence heatmap (Fig. S15B, Table S5E) identified a motif cluster, cluster 6, whose motifs showed strong patterns of co-occurrence at male class I DARs, while cluster 9 motifs showed strong patterns of co-occurrence at female class I DARs. In an alternative analysis, motifs strongly correlated with each other (Spearman correlation ≥0.5) were retained and used to prepare DAR-motif heatmaps (Fig. S16-S17; data in Table S6A-S6H). Examples of motifs pairs showing co-occurrence included PAX3 and PAX7, which were strongly correlated with each other in set D1 DARs (Table S6B). PAX3 and PAX7 jointly activate a large panel of genes involved in muscle stem cell function [101]. Another co-occurrence pair, HNF1A and HNF1B, was found in DAR sets D1 and E1 (Table S6B and S6F). HNF1A and HNF1B form both homo and heterodimers that regulate genes controlling homeostasis of lipid metabolism [102].

### Sex differences in intrahepatic cell-cell communication

We used CellChat [103] to elucidate GH-regulated sex differences in cell–cell communication patterns by comparing the potential for ligand-receptor interactions within and between each liver cell type for female livers (F1,F2) to those for control male (M) and cGH-infused male liver (M-cGH) (Fig. 6, Table S7). Similar numbers of ligand-receptor interactions (n=90 to 112) were found in female compared to male and cGH-infused male liver (Fig. S18). The interactions were grouped into 23 signaling pathways (Fig. S18). Three pathways were active in female and/or in cGH-treated male liver but not in control male liver (Fig. 6). These pathways involved vascular growth factors (ANGPT, VEGF) and cell adhesion molecules (JAM, seen in M-cGH only). Previous studies have found higher circulating levels of VEGF protein in female as compared to male mouse(?) liver [104]. Further, VEGF is a major angiogenic factor in female reproductive organs [104]. Next, we compared cell-cell communication differences between male liver and the other three samples (F1, F2, male-cGH) and found robust differences in communication patterns for four signaling pathways (SEMA3, SEMA6, HGF, CDH). Semaphorins are extracellular signaling proteins essential for axonal guidance, vascular development and tumorigenesis [105]. We observed decreased interactions involving Sema3 and Sema6 signaling in both female and cGH-feminized male liver as compared to control male liver. We also observed decreased intracellular communication involving HGF (hepatocyte growth factor) signaling between hepatic stellate cells and cholangiocytes and immune cells in female and cGH-treated male liver compared to male liver. HGF is essential for regeneration during liver injury [106]. Finally, we observed increased cadherin (CDH) signaling between hepatocytes and hepatic stellate cells in female liver and cGH-treated male liver, as compared to male liver. Cadherins mediate calcium dependent cell-cell adhesion and cell-extracellular-matrix adhesions are essential for tissue integrity [107, 108].

**Fig. 6.**
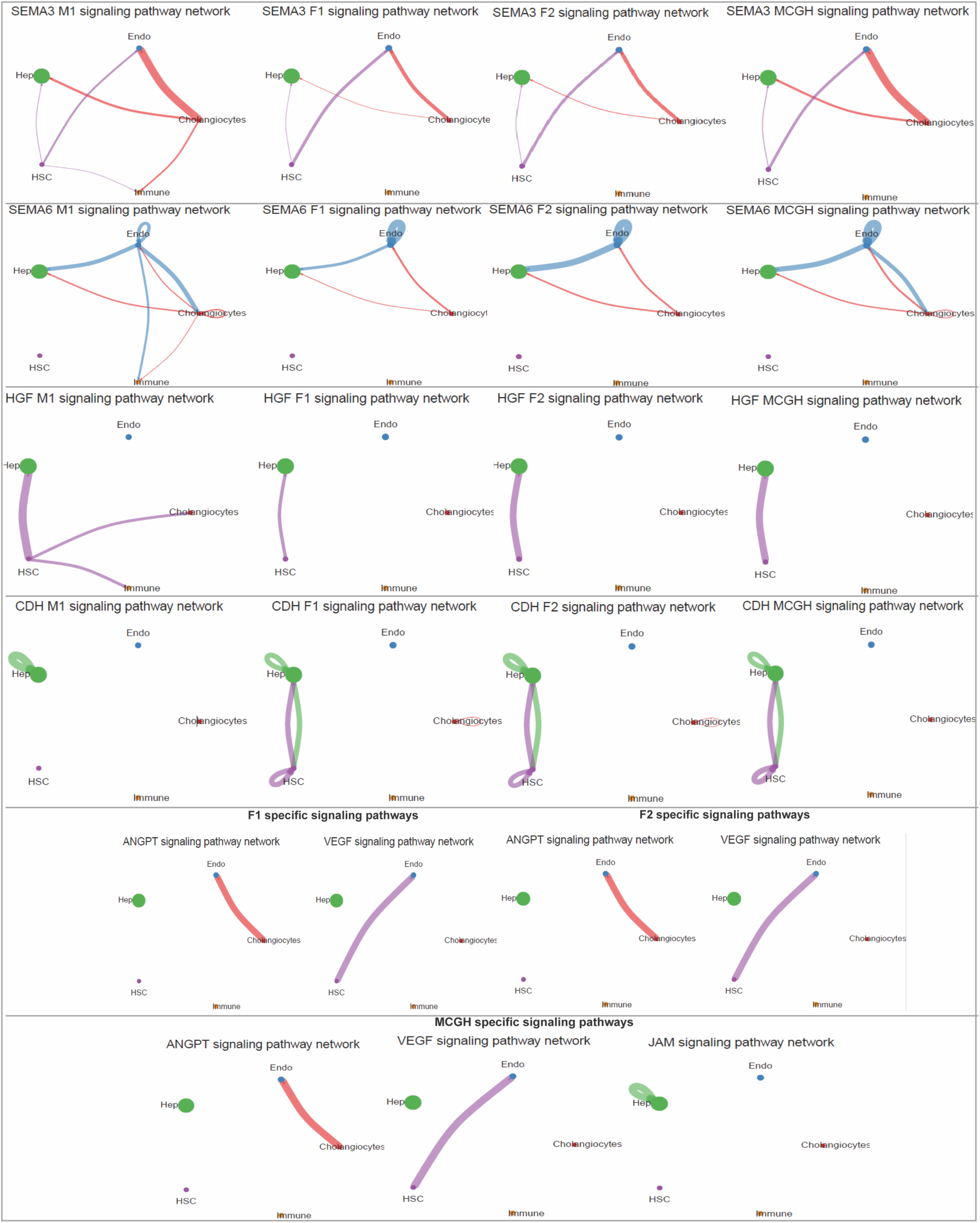
Sex-biased changes in cell-cell communication. Circle plots showing examples of sex-biased intercellular communication patterns in mouse liver across four samples: control male (M), control female replicate-1 (F1), control female replicate-2 (F2), and male treated with cGH (M-cGH). Edge weights are proportional to the interaction strength. A thicker edge line indicates a stronger signal, and circle sizes are proportional to the number of cells in each cell type.

## Discussion

We employed single cell transcriptomics and epigenomics to profile single nuclei extracted from frozen mouse liver to identify liver cell type-specific regulatory elements and their associated TF motifs and to elucidate transcriptional and epigenetic regulatory processes that govern sex differences in liver gene expression. Genome-wide maps of open chromatin regions were obtained for each of five major liver cell types to facilitate functional analysis of *cis*-and *trans*-acting regulatory elements and their connections with downstream gene targets. Traditionally, scRNA-seq expression profiling and scATAC-seq accessibility profiling has been performed using different cell populations and then jointly associated and integrated using computational algorithms to deduce linkages between regulatory elements and target genes [33, 109]. However, this approach faces the intrinsic limitation of using distinct cell populations for each analysis. Here, we used 10x Genomics multiOmic technology to simultaneously measure gene expression (RNA levels) and chromatin accessibility (ATAC) in the same individual cells and thereby obtain paired transcriptomic and epigenomic atlases for nuclei from livers of male, female and cGH-infused male mice. The rich datasets obtained were used to identify liver cell type-specific genomic regulatory elements generally, and more specifically, to discover linkages between sex-dependent regulatory elements and their gene targets, which underly sex differences in liver physiology and disease susceptibility in mouse liver. We characterized thousands of sex-biased DAR−gene linkage interactions, both promoter-proximal and gene distal, a majority of which involved positive, stimulatory (enhancer-like) interactions and a minority of which involved negative regulatory interactions between sex-biased DARs and gene targets with the opposite sex bias. Overall, we identified 127,957 accessible chromatin regions (ATAC sites) across a population of 46,188 nuclei representing all three biological conditions, 59% of which overlap accessible chromatin regions previously identified as DNase-I hypersensitive sites (DHS) in bulk liver tissue (Table S1C). 6,543 ATAC regions were found to be differentially accessible between liver cell types (Fig. 1B, Table S2A), of which 683 were stringently specific for a single liver cell type. Further, we identified 4,748 DARs that were robustly sex-biased and responded to cGH treatment in hepatocytes (Table S2C).

Analysis of snRNA-seq data identified 937 genes that showed sex-biased, cGH-regulated expression in hepatocytes (Table S2D) with no major sex differences found in the non-hepatocyte populations, consistent with our findings using an independent set of liver samples [42]. Some 200 of these sex-biased genes could be characterized into Hypox response class I or class II [25, 26] based on available data from bulk RNA-seq datasets (Table S2F). Linkage analysis identified interactions between sex-biased ATAC-seq regions and the set of 937 sex-biased genes, with a subset linked to hypox response class defined genes (Table S3A). We used chromVAR and motif enrichment analysis to identify many cell type-specific motifs (Fig. 3A, Fig. S13B, Table S4B) and to identify 425 sex-biased TF motifs representing 80 motif clusters (Fig. 3B). Further, motif co-occurrence analysis (Fig. 5) identified motif clusters that likely co-regulate chromatin accessibility and downstream transcriptional responses. Finally, we mapped the global landscape of intercellular signaling between liver cell types and explored sex-biased changes in intrahepatic cell-cell communication (Fig. 6). Together, these findings highlight the utility of multiOmic analysis for mapping sex-dependent regulatory interactions on a global scale, generating new hypotheses for TFs that contribute to sex differences in hepatocytes.

During the course of this work, we observed that some of the liver nuclei – both hepatocytes and NPCs – were apparently contained varying levels of mature, spliced forms of several abundant, hepatocyte-specific secretory protein RNAs (e.g., Alb, Serpina1c, Ttr, Apoa1), which we surmise originate from low level contamination of the extracted nuclei by outer nuclear membrane-bound polyribosomes and their attached hepatocyte secretory protein RNAs [42]. We devised a novel computational strategy to identify and then exclude from downstream analysis such liver nuclei, which appeared as a distinct population when plotting total Intronic vs Exonic UMI counts for each nucleus and could be filtered out by applying a supervised machine learning support vector machine algorithm (Figs. S20-S23). The impact of this apparent cross contamination of NPC-derived liver nuclei with spliced hepatocyte secretory protein RNAs was most readily apparent when using the widely employed full gene body counting method for snRNA-seq (Fig. S24A), but we found could be mitigated by using a modified UMI counting method that we termed Intronic plus Mono-exonic UMI counting, and which yielded cleaner transcriptomic profiles (Fig. S24). Thus, intronic sequence reads, which are largely derived from unspliced RNAs, were counted and used to represent the expression of multi-exonic genes, whereas exonic read counting was employed for mono-exonic genes. Given that a majority of snRNA-seq sequencing reads typically map to intronic regions of the genome [110], our exclusion of exonic reads from multi-exonic genes did not result in a large decrease in sensitivity for detection of gene expression. A further advantage of Intronic plus Mono-exonic UMI counting in the context of the present mutiOmic analysis, is that unspliced intronic read counts are better correlated with transcription rates that mature RNA sequence reads [111], and therefore better suited for correlative linkage analysis with sex-biased ATAC regions that undergo highly dynamic changes in accessibility associated with changes in gene transcription rates.

Chromatin accessibility analysis depends on accurate peak calling, and this can be a challenge in tissues such as liver, where hepatocytes constitute the major cell population and peaks specific to non-hepatocytes may be missed when accessibility is determined by bulk tissue analysis. For each biological condition, we used MACS2 [30] to identify individual snATAC-seq peaks across clusters representing all major liver cell types, then combined the peak calls into a unified set of 127,957 ATAC peaks using Signac. MACS2 peak calling is reported to be better at identifying robust and distinct peaks than 10X Genomics Cell Ranger ARC, which tends to merge multiple distinct peaks into a single region [34]. A majority (59%) of the snATAC regions we identified overlap with our published bulk mouse liver DHS datasets from the same mouse strain. The remaining 41% comprise novel ATAC peaks whose mean normalized ATAC activity was ∼5-fold weaker than that of the bulk liver DHS-overlapping ATAC peaks. Moreover, the novel ATAC peaks showed 4.3-fold higher mean maximal ATAC activity across the major NPC populations than in hepatocytes. Thus, many of the novel ATAC peaks reported here are preferentially accessible in one or more NPC populations, which helps explain their absence in bulk liver DHS sets, where open chromatin signals from hepatocytes, representing ∼60-70% of all liver cells, are expected to dominate the DHS signal. Finally, the 41% ATAC peak set comprised only 9-13% of the sex-biased ATAC regions from DAR sets D1 and E1, consistent with the deficiency of sex-biased DARs in NPCs as compared to hepatocytes.

The combined snRNA-seq + snATAC-seq dataset was used to compute statistically robust DAR–gene linkages, which allowed us to address the fundamental question of how constraining TAD boundaries are to gene regulation. We investigated this question for sex-biased DAR–gene interactions (linkages), a majority of which were localized within the same TAD; thus, sex-biased gene-regulatory region interactions are in part constrained by TAD boundaries. Furthermore, the vast majority (>97%) of linkages between sex-biased, cGH-responsive DARs and correspondingly regulated sex-biased genes were positively correlated, consistent with these sex-biased DARs serving as enhancers, a proposal supported by our chromatin state analysis of DAR regions. Interestingly, we identified a much smaller number of negatively correlated linkages between sex-biased DARs and genes of the opposite sex specificity (e.g., male-biased DAR linked to female-biased gene), indicating a novel repressive role for these DARs in regulating sex bias in hepatocytes.

More than half of sex-biased DARs that were exclusively linked to non-sex-biased genes mapped to different TADs than their linked genes; those DARs tended to show lower linkage correlations and were found at much greater distances than in the case of sex-biased DARs linked to sex-biased genes. Conceivably, some (perhaps many) of the interactions between sex-biased DARs and non-sex-biased genes might be false positives driven by weak correlative associations. However, our finding that the linked non-sex-biased gene targets of sex-biased DARs show a broad range of strong functional enrichments (Fig. S13) suggests that many of these linkages are, in fact, specific and not random correlative associations. It is unclear, however, why the linked genes do not show sex bias, given their linkage to DARs with sex-biased chromatin accessibility. One possibility is that the non-sex-biased genes linked to a sex-biased DAR are also linked to one or more non-sex-biased DARs that normally dominate their regulation. Conceivably, the sex-biased DARs associated with these non-sex-biased genes could serve as latent sex-biased enhancers, whose sex bias only becomes manifested under conditions of stress. Many hepatic stressors act in a sex-biased manner, by either activating or repressing genes that have a latent tendency to exhibit sex bias in their expression under conditions of stress [2, 112, 113].

Chromatin accessibility and gene expression studies in mouse liver using bulk RNA-seq have identified four well studied liver TFs, STAT5B, CUX2, HNF6 and BCL6, that mediate dimorphic gene expression patterns in liver [29]. ChromVAR identified many other, novel liver cell type-specific motifs not found in earlier bulk DNase hypersensitivity assays (Fig. 3A). We also employed motif enrichment analysis to discover robust sets of cell type-specific motifs (Table S4B.4). TF motif enrichment also identified unique TF motifs associated with various sets of sex-biased DARs, including those linked to sex-biased genes, those linked to non-sex-biased genes, and also those linked to sex-biased genes showing distinct responses to hypox (Table S4C). Many of the motifs that were differentially enriched between male and female liver DAR sets were consistent with previous findings, but we also discovered enriched motifs for TFs that were not previously associated with liver sex-bias. Finally, motif co-occurrence analysis helped identify motif clusters that may co-regulate subsets of sex-biased DARs (Fig. 5). To address the concern that this analysis generated results for large numbers of redundant motifs, we used RSAT which is a motif clustering tool, which resulted in grouping of 841 motifs into 137 clusters (Table S5A).

Finally, our discovery of TF motifs whose occupancy at regulatory regions controlling MASLD-relevant genes, as indicated by ATAC-seq footprinting, provides molecular insights into the long-standing paradox of male predominance in fatty liver disease despite females’ greater relative adiposity. The identification of distinct occupancy profiles for TFs at male-biased versus female-biased MASLD-linked DARs leads us to propose that a specialized subset of the sex-biased transcriptional regulatory landscape is wired to control sex differences in metabolic disease susceptibility. The preferential occupancy of Cdx, Meis1, and Pbx factor motifs at male-biased MASLD gene-linked DARs is particularly noteworthy given the known roles of these homeodomain families in hepatic lipid metabolism and fibrosis. Meis factors, which function as obligate partners with Pbx proteins in stabilizing heterodimeric complexes [92–94], are emerging as critical regulators of hepatocellular carcinoma progression [95], suggesting that their enhanced occupancy in male liver may contribute not only to early MASLD but to the trajectory toward advanced liver disease and malignant transformation characteristic of male-predominant HCC. Conversely, the selective enrichment of occupied Nr1h2 (LXR-β) and Nfatc1 motifs at female-biased MASLD gene-linked DARs illustrates how the female GH secretion pattern activates a distinct regulatory program: while Nr1h2 promotes lipogenesis (potentially predisposing females to steatosis), concurrent female-biased activation of MASLD-protective genes such as Trim24 and Fmo2 may protect against the progression from simple steatosis to fibroinflammatory disease. The functional significance of these sex-selective TF regulatory networks is underscored by their positioning at chromatin regions constrained within topologically associated domains—consistent with our TAD analysis—suggesting that these are not incidental associations but rather reflect evolutionarily selected *cis*-regulatory architectures that implement sex-appropriate metabolic strategies. Future investigation of whether translationally relevant interventions targeting these sex-selective TF networks (e.g., selective Nr1h2 inhibition in females, or Meis/Pbx interference in males) might achieve improved sex-tailored therapeutic outcomes for MASLD is warranted.

## Conclusion

This study provides the first comprehensive single-nucleus multiOmic atlas mapping the regulatory landscape of GH-dependent sexual dimorphism in mouse liver. By simultaneously profiling chromatin accessibility and gene expression in thousands of individual liver nuclei, we discovered that sex-biased regulatory elements are almost exclusively hepatocyte-specific, preferentially regulate genes within the same topologically associated domain, and are highly responsive to changes in temporal plasma GH patterns. Our identification of sex-biased TF motifs predicted to be occupied—including both established GH-responsive regulators and numerous novel factors—significantly expands the repertoire of TFs implicated in hepatic sexual dimorphism. Critically, we established molecular links between specific *cis*-regulatory elements and their associated TF motifs and sex-biased genes involved in MASLD, providing mechanistic insights into the male bias in fatty liver disease susceptibility. The cooperative TF networks revealed by motif co-occurrence analysis suggest complex regulatory logic underlies sex-dependent gene expression programs. The multiOmic resource described here, comprising cell type-resolved chromatin accessibility maps, robust DAR-gene linkages, cell type-specific and sex-biased TF motif enrichments, and intercellular communication networks, establishes a foundational framework for understanding how transcriptional regulation controls sex differences in liver physiology and disease predisposition, with potential translational implications for sex-specific therapeutic strategies targeting liver metabolic diseases.

## Methods

### Animal work and tissue procurement

Mouse work was carried out in compliance with procedures approved by the Boston University Institutional Animal Care and Use Committee (protocol # PROTO201800698) and consistent with ARRIVE 2.0 Essential 10 guidelines [114], including study design, sample size, randomization, experimental animals and procedures, and statistical methods. Male and female CD-1 mice (ICR strain) were purchased from Charles River Laboratories (Wilmington, MA), fed standard rodent chow, and housed in a temperature and humidity-controlled facility on a 12 h light cycle, beginning at 7:30 AM. Where indicated, male mice (8 wk of age) were anesthetized with isoflurane and implanted with an Alzet osmotic minipump (cat. # 2002; Durect Corp., Cupertino, CA) subcutaneously, to deliver recombinant rat GH as a continuous infusion (cGH) at 25 ng/g BW/h then euthanized 12 days later. GH (purchased from Dr. Arieh Gertler, Protein Laboratories Rehovot, Ltd., Rehovot, Israel) was dissolved in 30 mM NaHCO_3_ and 0.15 M NaCl containing 0.1 mg/ml rat albumin (Sigma-Aldrich, catalog no. A6414), as described in [15]. Mice were euthanized under CO_2_ anesthesia by cervical dislocation at 8-10 wk of age, and livers were collected at a consistent time of day (between 10:30 AM and 11:30 AM) to minimize the impact of the circadian effects on liver gene expression [115]. Livers were excised and flash frozen in liquid N_2_ then stored at -80°C.

### Isolation of single nuclei from frozen mouse liver

Frozen mouse liver tissue (a total of ∼150 mg) was combined from each of n=3-4 biological replicates per group (biological condition) and processed together as a single sample to obtain liver nuclei representing the 3-4 livers included in each group. We adapted the Frankenstein protocol developed to extract nuclei from frozen mouse brain tissue [110, 116, 117] and optimized it to obtain high-quality nuclei from frozen mouse liver tissue, as follows. Frozen mouse liver samples were minced in the cold room, then added to 1 ml of ice-cold Nuclei EZ lysis buffer (cat. # NUC101, Sigma-Aldrich) in a 2-ml glass Dounce homogenizer, placed on ice for 1 min, and then homogenized on ice using 10 strokes of Dounce pestle A, followed by another 10 strokes with Dounce pestle B. Samples were kept on ice during homogenization to minimize RNA degradation. To reduce nuclei concentration and minimize aggregation of the nuclei during the later washing steps, another 1 ml of ice-cold Nuclei EZ lysis buffer was added to the lysate in the Dounce homogenizer tube and mixed gently, followed by a 5 min incubation. The lysate was filtered through a 40-mm cell strainer mesh (cat. #431750, Corning Life Sciences) and collected in a 50-ml conical tube. The lysate from each sample was then split into two Eppendorf tubes, which were not recombined until the final nuclei permeabilization step, below. The lysed nuclei in each tube were pelleted by centrifugation in a swinging bucket rotor at 500 x g for 5 min at 4°C, after which the pellet was gently resuspended by adding 1 ml of fresh EZ lysis buffer to each Eppendorf tube. The 5-min incubation and 5 min centrifugation steps were then repeated a second time. After removing the supernatant, an 0.5 ml aliquot of Nuclei Wash and Resuspension Buffer (2% FBS in phosphate-buffered saline, with 1 U/mL of Protector RNase Inhibitor (cat. #3335399001, Millipore-Sigma) added immediately before use) was added to each tube without disturbing the pellet, followed by a 5 min incubation on ice to allow for buffer exchange. The pellets were then resuspended after adding an additional 0.5 ml of Nuclei Wash and Resuspension Buffer to give a total volume of 1 ml in each tube, and then centrifuged at 500 x g for 5 min at 4°C. The supernatant was removed, and the pelleted nuclei were resuspended in 1 ml of Nuclei Wash and Resuspension Buffer. A 10 ml aliquot was removed to visualize and quantify the nuclei on a Cellometer Vision instrument (Nexcelom Bioscience, Lawrence, MA) using the AOPI Staining Kit (cat. #CS20106, Nexcelom Bioscience, Lawrence, MA). Typically, the concentration of nuclei ranged from 5,000-6,000 nuclei/ml when starting with 150 mg liver tissue, with viability typically < 10%, as determined by AOPI staining. High quality nuclei showed an intact nuclear membrane without peripheral blebbing (Fig. S1A).

### Nuclei permeabilization and MultiOmic ATAC + Gene Expression library preparation

The resuspended nuclei were kept on ice and transferred within 1 h to the MIT BioMicro Center (Cambridge, MA), where the nuclei were permeabilized using a protocol described in the 10x Genomics protocol # CG000375, Rev B: Nuclei Isolation from Complex Tissues for Single Cell MultiOmic ATAC + Gene Expression Sequencing. Permeabilization of the nuclei was performed immediately prior to Gel Bead-In EMulsion (GEM) formation and 10X MultiOmic ATAC + Gene Expression library preparation, to help maintain RNA and DNA integrity. To permeabilize the nuclei, the nuclei were pelleted at 500 x g for 5 min at 4°C and then resuspended in 100 ml of freshly prepared ice cold 0.1x Lysis Buffer (10 mM Tris-HCI, pH 7.4, 10 mM NaCl, 3 mM MgCl_2_, 0.01% Tween 20, 0.01% NP40, 0.001% Digitonin, 1% BSA, 1 mM DTT and 1 U/ml Protector RNase Inhibitor (cat. #3335399001, Millipore-Sigma) by pipetting the sample up and down gently 5 times, followed by a 2 min incubation on ice. Freshly prepared ice-cold Wash Buffer (1 ml of 10 mM Tris-HCI, pH 7.4, 10 mM NaCl, 3 mM MgCl_2_, 1% BSA, 0.1% Tween 20, 1 mM DTT and 1 U/ml Protector RNase Inhibitor) was then added and the sample was mixed by pipetting up and down gently 5 times. After centrifugation at 500 x g for 3 min at 4°C, the supernatant was removed and replaced with the diluted Nuclei Buffer provided in the Chromium Next GEM Single Cell MultiOmic ATAC + Gene Expression Reagent Bundle kit (cat. # PN-1000285, 10X Genomics). Immediately prior to the GEM formation procedure, the permeabilized nuclei were filtered through a 40 mm Flowmi® Cell Strainer (cat. # BAH136800040, Millipore-Sigma) to removed aggregated nuclei. The filtered nuclei were then quantified using trypan blue staining and diluted to an estimated concentration of 2,500 nuclei/ml. To prepare GEMs, nuclei were loaded onto a 10x Chromium instrument at the MIT BioMicro Center with a *targeted* recovery of 6,000-7,500 nuclei per sample (but see actual results, below), followed by MultiOmic ATAC + Gene Expression library preparation following the manufacturer’s protocol (Chromium Next GEM Single Cell MultiOmic ATAC + Gene Expression Reagent Bundle, cat. # PN-1000285).

### Illumina sequencing

MultiOmic ATAC + Gene Expression libraries [24] were prepared for 4 separate samples: adult male liver (M), adult female liver (2 independent pools, designated F1 and F2), and cGH-treated adult male liver (M-cGH), each comprised of nuclei extracted from n=3-4 individual mouse livers, as described above. The samples were prepared in two batches: one batch (‘batch G193’) included multiOmic samples M and F1 (fastq files G193M1/G193M5 and G193M3/G193M7, respectively) and the other batch (batch ‘G190’) included multiOmic samples F2 and M-cGH (fastq files G190M2/G190M6 and G190M3/G190M7, respectively) (Table S1A). For each sample, we performed pilot Illumina sequencing to obtain 40-50 million read pairs per sample prior to deeper sequencing to better estimate the number of nuclei per library and hence the total required sequencing depth per sample, based on the following parameters calculated for each library from the pilot sequencing run: number of nuclei, percentage of ATAC-seq high-quality fragments in cells, percentage of GEX Fraction of transcriptomic reads in cells. This pilot sequencing was performed based on our finding that, in practice, the actual nuclei number obtained in each library was based on 10X Genomics Cell Ranger ARC v2.0.1 analysis, varied somewhat between and was up to 2.5-3-fold higher than the targeted recovery values of 6,000-7,500 nuclei per sample (Table S1A). This discrepancy may be due to variability in nuclei aggregation and sampling effects. For the ATAC-seq libraries, we aimed for a total sequencing depth of 15,000 reads per cell (2 x 150 bp pair-ended reads, 28 bp for cell barcodes and UMI, and 8 bp for sample index), and for the gene expression (GEX) libraries, a total sequencing depth of 20,000 reads per cell (2 x 150 bp pair-ended reads and 2 x 10 bp dual index reads). Based on Cell Ranger ARC analysis, MultiOmic data was obtained for an estimated 16,676-20,000 cells per sample. The total sequencing depth for the ATAC-seq libraries was 254-318 million read pairs per sample, corresponding to 13,713-15,894 mean raw ATAC read pairs per cell. The fraction of high-quality ATAC fragments in cells was 76-89%, and a total of 91,762-112,923 ATAC peaks covering 2.9-3.4% of the mm10 mouse genome were discovered in each sample, with 37-48% of the high quality ATAC-seq fragments overlapped ATAC-seq peak regions (Table S1A). Further, the enrichment score for ATAC peaks at gene TSS was highly significant, ranging from 5.48-6.60. The GEX libraries were sequenced to a total depth of 347-616 million read pairs per sample, corresponding to 17,355-30,783 mean GEX reads per cell, with 89-96% of the transcriptomic reads in cells. The GEX libraries had a median of 2,674-6,113 UMI counts per cell and a median of 1,589-3,043 genes detected per cell, resulting in the detection of 51,152-62,034 genes per sample; these numbers include a significant fraction of the ∼ 48,000 liver-expressed lncRNAs included in the custom GTF file used for analysis of the snRNA-seq datasets (below). Full technical details of the sequencing output for each of the four ATAC and GEX libraries are provided in Table S1A.

### Gene expression analysis using Intronic + Mono-exonic Counting method

Sequencing data were processed and aggregated with Cell Ranger ARC v2.0.1. We used a custom mouse mm10 GTF file, multiOmic_genes_mm10.gtf (GTF file formatted per Cell Ranger ARC reference format using the mkref command). This GTF file includes all mouse RefSeq protein coding genes as well as ∼48,000 liver-expressed lncRNA genes [30] and was used to implement Intronic + Mono-exonic Counting, whereby gene expression is quantified by counting snRNA-seq UMIs for all intronic regions that map to multi-exonic genes, and for all exonic regions that map to mono-exonic genes, for reasons outlined below.

In earlier work we observed a variable extent of contamination of our 10X Genomics snRNA-seq profiles with mature secretory protein mRNAs derived from hepatocytes (Alb, Serpina1c, Ttr, Apoa1) and mitochondrial genes [39]. We therefore devised a method to filter out low quality nuclei using the following four steps. **Step 1:** For each individual snRNA-seq library, we plotted the total intronic UMI vs. total exonic UMI per nucleus; for some libraries, this graph revealed two distinct populations of nuclei (populations I and II), which were separated based on a supervised learning algorithm (support vector machine model [118]) that analyzes the data for classification (Fig. S20-S23). **Step 2:** The snRNA-seq nuclei from each individual sample were analyzed by graph-based clustering in Seurat (4.1.0) and the clusters were visualized in a UMAP. Dotplots presenting gene expression values for key marker genes were generated to assign cell type labels to each UMAP cluster. **Step 3:** The clustered nuclei were projected to the Intronic UMI vs Exonic UMI plot from step 1 to determine their distribution between the two populations of nuclei. Clusters of nuclei without unique, unambiguous liver cell type marker gene assignments, and clusters with high mitochondrial gene expression included many nuclei that mapped to population II; these clusters were eliminated with nuclei mapping to population II were filtered out prior to UMAP clustering downstream analysis (Fig. S20-S23, *left*). Nevertheless, gene expression profiles for liver non-parenchymal cell clusters (i.e., non-hepatocyte clusters) were apparently contaminated by RNAs for several abundant secretory protein that are well established hepatocyte markers (Alb, Serpina1c, Ttr, Apoa1) when the standard (default) full gene body counting method was used (Fig. S20-S23, *right*). **Step 4**: To address this issue, we modified the counting method to specifically count UMIs for snRNA-seq reads that mapped to intronic sequences (i.e., unspliced reads), in the case of multi-exonic genes, and to count UMIs that mapped to exonic sequences for mono-exonic genes (to ensure inclusion of these genes in downstream analysis). This hybrid counting method yielded liver non-parenchymal cell clusters that were largely free of hepatocyte-derived secretory protein contamination (Fig. S20-S23, *right*; also see Fig. S24).

### Samples aggregated for joint snATAC-seq and snATAC-seq (multiOmic) analysis

Fastq sequencing files from four 10X Genomics multiOmic sequencing libraries representing the 3 liver groups described above (control male, control females F1 and F2 (biological replicates), and cGH-treated male) were aggregated with data from three other 10X Genomics multiOmic sequencing libraries. The latter 3 multiOmic datasets were from the same batches of multiOmic sequencing libraries and were prepared from liver nuclei extracted from male and female mice exposed to the CAR (*Nr1i3*) agonist ligand TCPOBOP for 14 days [119] (details to be reported elsewhere). Data from all 7 multiOmic datasets, and representing adult mouse livers under 5 distinct biological conditions (control male, control females F1 and F2, cGH-treated male, TCPOBOP-treated males M-TCPO1 and M-TCPO2, and TCPOBOP-treated female), was aggregated to obtain a single harmonized UMAP, and for discovery of a single, integrated set of 127,957 ATAC-seq peaks using the peak caller MACS2 (Table S1C, and see below). The full set of 7 multiOmic datasets was also used as an input for peak-gene linkage analysis in Cell Ranger ARC, capitalizing on the added diversity of including all 7 independent samples representing 5 biological conditions to increase the statistical power of the linkage analysis. All other downstream analysis, including identification of sex-biased DARs, motif enrichment analysis, motif co-binding analysis and CellChat analysis was performed using multiOmic datasets from the control male, 2 control female and male + cGH treatment multiOmic samples only, each of which is comprised of nuclei from n=3-4 biological replicate livers per group, as noted above.

### MultiOmic data processing

The multiOmic (RNA + ATAC) data was processed using Signac (v1.1.0) [34] and Seurat (4.1.0) [33]. ATAC-seq peaks were identified separately for each individual sample and for each of 7 liver cell types, including 3 hepatocyte subclusters (periportal (PP), midlobular (Mid) and pericentral (PC), using MACS2 from within the Signac package [31]. This analysis was implemented using the function CallPeaks in Signac 1.1.0 with the arguments ‘group.by’ = ‘celltype_and_condition’ followed by keepStandardChromosomes() with “coarse” pruning, and subsetByOverlaps() using the Encode blacklist sites (blacklist_mm10.bed). Next, we used the CreateChromatinAssay function in Signac, a combine peaks option that aggregates genomic ranges of all peaks from all of the individual samples to create a “peaks” assay.

Data were filtered to retain nuclei with low mitochondrial contamination (<10%), high RNA content (nGene per cell > 400) and high ATAC counts (nATAC count per cell > 500, nucleosome signal score < 2, and TSS enrichment score > 1). Doublets were removed by combining the results from scDBlFinder [120] for snRNA-seq and the Amulet method [121] for snATAC-seq, which assumes that, in a diploid cell, any given genomic region should be captured at most twice. Ultimately, we retained for downstream analysis nuclei that passed quality control filters from *both* the RNA-seq and ATAC-seq modalities (overall, 74% of all nuclei; Table S1B). The filtered snRNA-seq samples were processed using Seurat (v4.1.0). Highly variable genes were identified using the FindVariableGenes function of Seurat with default parameters. snATAC-seq counts were processed using Signac (v1.1.0) package. Fragment counts for each ATAC-seq peak were quantified per cell using the FeatureMatrix function in Signac. We apply SCTransform to normalize RNA-seq counts and TFIDF to normalize ATAC-seq peaks. Dimensionality reduction for snRNA-based gene expression was performed by principal component analysis (PCA) and latent semantic indexing (at n=10), a topic modeling approach to reduce dimensionality of the ATAC-seq data. Batch correction for both the RNA-seq and ATAC-seq modalities was performed using Harmony (v1.0) [32] with default parameters. Then, we combined latent semantic indexing reduction and PCA reduction to construct a weighted nearest neighbor (WNN) graph using the WNN method [13] in Seurat, which reflected both modalities. This WNN graph was used to find clusters using the FindClusters function in Seurat (v4.1.0). The clusters obtained were visualized using Uniform Manifold Approximation and Projection (UMAP) at resolution 0.25 and PC=10.

### DAR to gene linkages

We used a bed file listing the 127,957 ATAC-seq peaks, discovered using MACS2 (above), as input for feature linkage analysis using Cell Ranger ARC and implemented using the ‘reanalyze’ command. Feature linkages mapping snATAC-seq regions (putative regulatory elements) to their gene targets were calculated by Cell Ranger ARC based on correlations between DNA accessibility and the expression of genes within a distance of 1 million bp and exported to Table S3A. The correlation values obtained provide an indication of the strength of the interaction, with positive correlations identifying ATAC regions that likely act as enhancers and negative correlations inferring a repressor-gene target relationship. ATAC-seq−gene pairs with strong linkage correlations are considered to be “co-expressed” and enrich for a shared regulatory mechanism (https://support.10xgenomics.com/feature-linkage). Correlation with lower significance (-log 10 FDR < 5) were removed from the final output.

### TAD localization

TAD boundaries previously defined by our lab [37] and refined as described [42] were used to assign TAD annotations to each ATAC region, and to its linked genes, using the Bedtools overlap command. The gene TSS and the midpoint of the ATAC peak region were used for TAD assignment. Next, we compared TAD assignments of ATAC peak regions and of their linked to gene targets using the Identical function in R. ATAC regions whose linked genes had the same TAD assignment were labeled “same TAD”; otherwise, they were labeled “different TAD”. ATAC peaks linked to multiple genes were labeled “mixedTAD” and excluded from downstream analysis if their linked genes mapped to two or more TADs. For the sets of DARs analyzed in Fig. S6, we used correlation values and distances between DARs and their linked genes, analyzed separately for same TAD and different TAD ATAC peaks. We compared the distributions of correlation and distance in multiple unmatched sets of DARs in a nonparametric manner, using the Kruskal-Wallis test to perform pairwise comparisons between each set of DARs, followed by Bonferroni FDR multiple comparison tests to obtain adjusted p-values; these are marked with asterisks defined as follows: adj-p > 0.05 (ns), adj-p ≤ 0.05 (*), adj-p ≤ 0.05 (**), adj-p ≤ 0.01 (***) and adj-p ≤ 0.001 (****).

### Chromatin state analysis of DARs

The chromatin state at each of the 127,957 ATAC-seq regions was characterized in both male and female liver using chromatin state maps that we previously developed for both male and female mouse liver [38] using ChromHMM [122]. Specifically, we used a 14-state model (states E1-E14; emission parameters shown in Fig. S11E) that was defined using a series of 6 informative histone marks, and DNase-I hypersensitivity data collected for adult male liver, and separately, for adult female liver; the genome-wide chromatin state models obtained encompass sex differences in chromatin state and chromatin structure and their relationships to sex-biased gene expression [38]. Bedtools overlap was used to assign the chromatin state in each sex to each ATAC-seq peak (Table S1C, columns BJ-BM). ATAC-seq peaks that overlapped genomic regions in two or more different chromatin states were assigned the state with the longest overlap in bp. The mapped chromatin states were used to create bar plots showing the chromatin state distributions, computed separately for male and female liver, for each DAR set (Fig. S11, Fig. S12).

### Comparison to published datasets of DNase-I hypersensitive sites

We downloaded three sets of DHS sites, as follows. Set I: 60,739 DHS (60,732 after conversion to mm10) DHS sites from [35]; Set II: 70,767 DHS (70,754 after conversion to mm10) from Table S5B of [36]; and Set III: 70,285 DHS (70,281 after conversion to mm10) from [37]. The genomic coordinates in mm9 were converted to mm10 using the UCSC genome browser LiftOver tool [123]. Next, we used Bedtools (v2.17.0) [124] to calculate the overlap between these three DHS sets and the set of 127,957 snATAC-seq peaks detected by MACS2. Peaks annotations reported earlier [36, 37] were used to characterize the 127,957 ATAC-seq peaks. See Table S1C for full details, including annotations of the 3 prior DHS datasets.

### Differentially accessible regions (DARs) and differentially expressed genes (DEGs)

Differential chromatin accessibility, and differential gene expression, between liver cell types and between biological conditions (male, female and cGH-treated male) was computed using the Loupe Browser (10x Genomics, Loupe Browser 6.0 (https://www.10xgenomics.com/), which uses the exact negative binomial test to test for differences in mean ATAC-seq sequence read density (cut sites per cell) in a given ATAC region, or differences in gene expression between groups of cells. Liver cell type-specific DARs were identified as follows. First, ATAC peaks that showed significant differential accessibility between cell types were identified using the Loupe Browser’s integrated Global Distinguishing function with a differential Log2 Fold-change >1 at FDR (Benjamini-Hochberg) < 0.05 (Table S2A). The resulting list of candidate cell type-specific DARs was filtered to remove ATAC peaks found in < 3% of nuclei (for non-hepatocyte clusters) or in < 5% of nuclei (for hepatocytes). The list was further filtered to removing any ATAC region that met the above criteria for DAR cell type specificity for 2 or more liver cell clusters. Further, DARs with a cell type specificity ratio ≤ 2 (hepatocytes, immune cells) or ≤ 4 (hepatocytes, hepatic stellate cells) were designated ’low specificity’ cell type-specific DARs and excluded from further analysis, resulting in a final list to 683 stringent cell-type-specific DARs (Table S1E). The pheatmap function in R was used to plot as a heatmap (Fig. 1B) the average number of Tn5 tagmentation sites within each DAR for each cell type, presented as a Z-score of the number of cut sites within each DAR scaled by row. We used the Loupe Browser’s integrated Locally Distinguishing function to calculate condition-specific DARs across five different pairwise comparisons: Male vs. Female F1, Male vs. Female F2, Male vs. cGH-treated male (M-cGH), Female F1 vs. M-cGH, and Female F2 vs. M-cGH, with a significance threshold of adjusted P-value (Bonferroni-corrected FDR) <0.05, except as noted. We used the same approach, based on the Loupe Browser’s Locally Distinguishing function, as described previously [39], to DEGs in each liver cell type for the same five comparisons, with results summarized in Table S2B.

### Hypox class sex-biased genes

Bulk liver RNA-seq datasets described earlier [26] were reanalyzed using an in-house RNA-seq pipeline to identify sex-biased genes, including sex-biased liver-expressed lncRNA genes (REF) whose expression was responsive to hypox at FDR <0.05 (Table S2G). The list of hypox class-defined genes were further filtered to remove any genes that did not show robust sex bias and GH responses in our snRNA-seq samples (see Table S2G for further details).

### Enrichment heatmap

Genes that were linked to DARs from DAR sets D1, E1, D2a and E2a (Fig. S5A.1) were used as input to Metascape [43] to perform multi-list comparative analysis and identify pathways that are shared between, or are specific to, these four gene sets. The set of all mouse RefSeq genes was used as a background and KEGG, Reactome, GO:BP, MF, and CC, GSEA hallmark gene sets were used as test sets for the enrichment analysis. The resulting clustered heatmap (Fig. S13A) depicts -log10 P-value of top enriched pathways across the four gene lists. See Supplementary Metascape Excel files for further details.

### Motif redundancy and motif clustering

To address the issue of motif redundancy, we used the RSAT matrix clustering tool [125] (https://rsat.france-bioinformatique.fr/teaching/matrix-clustering_form.cgi) to cluster a set of 841 motifs from JASPAR (2022) (https://jaspar.genereg.net/) [126] based on sequence similarity. The input was a set of positional weight matrices in JASPAR format. We used the average linkage method for clustering with default parameters to group the 841 motifs into 137 cluster families based on sequence similarity (Table S4A). This cluster information was used to group redundant motifs identified by motif enrichment analysis and was included in all Supplemental tables presenting motif enrichment and motif co-occurrence datasets. We created non-redundant co-occurrence heatmaps by selecting a single representative motif from each motif cluster defined by the RSAT clustering tool. The representative motif was defined as the motif in each cluster that had the lowest FDR value for motif enrichment across DAR sets D1 and set E1 (Table S4D).

### Motif activity analysis using chromVAR

Motif positional weight matrices from the JASPAR 2022 database [126] were used within ChromVAR (v1.6.023) [44] to calculate TF motif activity scores, which indicate the relative accessibility of each TF motif in a given cell cluster relative to the global average accessibility of the motif across all cells in the snATAC-seq dataset. Cell-type-specific chromVAR TF motif activities were calculated using the RunChromVAR wrapper in Signac (v1.1.0) [34] after inputting a matrix of the number of ATAC-seq fragments mapping to each of the 127,957 ATAC-seq peak regions (rows) for each liver cell in the snATAC-seq dataset (columns). A raw accessibility deviation score (equivalent to an intensity score) was calculated for each TF motif on a per cell basis, as follows: Score = (Observed – Expected)/Expected, where Observed = number of ATAC fragments summed across the set of ATAC peaks containing that motif, and Expected = number of fragments expected at each peak based on the global average of all cells and all peaks in the population. By aggregating ATAC-seq reads per cell across ATAC peaks containing the motif, the motif signal is much less sparse than the signal within an individual ATAC peak, which is intrinsically limited by the DNA copy number of 2 per diploid genome. The raw accessibility deviation score was then bias-corrected by ChromVAR using an equal number of background peaks matched for GC content and average chromatin accessibility and the output then scaled to give a Z-score for each motif and each cell. This bias-corrected Z-score, referred to as the *motif activity* score, indicates how significantly the motif’s accessibility in that cell deviates from the global average across all cells in the snATAC-seq sample, and can be displayed in a feature plot (Fig. 3A, bottom).

Cluster-level summaries were then generated by aggregating the per-cell motif activity scores across all cells in each cell type-specific cluster, followed by differential analysis of each cell type-specific cluster to obtain provisional lists of motifs that were differentially accessible in each cell cluster, at log2 fold-change >1 and FDR < 0.05, as compared to all other cells considered as a single group. A final list of cell type-specific TF motifs was obtained by removing any motifs that met the above thresholds for two or more liver cell type clusters.

Hepatocyte-specific motif activities were found to show lower differential activity scores compared to those associated with other, smaller cell clusters (e.g., endothelial cells and hepatic stellate cells) due to the strongly influence of the hepatocyte cluster on the global average used as the reference, as expected.

### TF motif enrichment at cell type-specific ATAC regions

In a complementary approach to discover cell-type-specific regulatory TFs, we searched for TF motifs that are statistically overrepresented in each set of cell type-specific DARs as compared to a defined background ATAC peak set. We considered separately two subsets of cell type-specific DARs: promoter-proximal DARs (peak center < 3 kb upstream or downstream of a protein-coding gene or a lncRNA TSS) and Distal DARs (≥ 3 kb from TSS) (Table S1E, column AV). Background peak sets (both promoter proximal and gene distal) were selected for each DAR set to match the ATAC-seq intensity (chromatin accessibility) of each cell type-specific DAR subset (Table S4B.5). We then used the FindMotif function built into chromVAR (motifmatchr R package (https://bioconductor.org/packages/motifmatchr)) to scan for motifs from the JASPAR 2022 database that were significantly enriched in the cell type-specific DAR set as compared to the background ATAC peak set. Significance was determined by a hypergeometric test, which gives the probability of observing the motif at the given frequency by chance in the foreground peak set as compared to the background peak set.

### TF motif enrichment at sex-biased ATAC regions

Sex-specific ATAC-seq regions were selected for motif analysis as follows. Initial analysis of the sets of sets of 3,518 male-specific, GH-repressed DARs and 1,230 female-specific, GH-induced DARs (DAR set D and DAR set E, Fig. S5A.1) revealed that 96% were within the top 70% of all 129,957 ATAC-seq peak regions when ranked by maximum hepatocyte ATAC-seq peak intensity across all samples, normalized to ATAC region width in nt (Table S3C, column B). We therefore restricted motif enrichment analysis, including background peak selection, to the top 70% (n= 89,600) of ATAC peaks (Table S3C, columns I, J). These 89,600 ATAC-seq peaks include 65,710 of 74,860 (88%) DHS defined by prior bulk liver DHS datasets (Table S3C, column L). Approximately 52% of the 89,600 ATAC peaks had enhancer state annotations (n = 46,473 enhancer state peaks in male liver; and n= 47,071 enhancer state peaks in female liver; Table S1C). The subset of the top 70% hepatocyte ATAC-seq activity enhancer state peaks within ATAC peak sets A1 and B1 (Fig. S5), which showed no sex bias or response to cGH infusion, were used as background peaks (Fig. S5B, Table S1C, column O). Motif enrichment was computed using the FindMotif function of chromVAR as described above for cell type-specific ATAC regions. Fig. S14 and Table S4E shows motif enrichment data as normalized Z-scores calculated using the scale function in R.

### ATAC-seq footprinting using TOBIAS

Footprinting analysis was performed using TOBIAS [90] to analyze TF protein occupancy in bulk liver ATAC-seq data obtained from 8 wk male and female CD-1 mouse liver, aggregated from n=3 individual STAT5 activity-positive [36] male livers and from n=4 individual female livers (all STAT5-activity low or negative). We analyzed TF binding at the following sets of DARs with enhancer marks (enhancer status based on chromatin state analysis for adult male and female mouse liver [38]): 308 male-biased, cGH-suppressed DARs (enhancer subset of DAR set D1) and 211 female-biased, cGH-induced DARs (enhancer subset of DAR set E1), defined in Table S1C, column Q. We also analyzed the subsets of these two enhancer DAR sets whose linked genes (determined by multiOmics peak-to-gene linkage analysis, above) are associated with a set of 15 male-biased genes and with a set of 13 female-biased genes whose gene functions have been shown to impact MASLD and associated liver diseases. Individual genes were designated MASLD-enabling or MASLD-protective, or ‘context-dependent’ in cases where the impact on disease susceptibility or progression varied with the context, based on a review of the relevant published literature (Fig. 5A).

Collectively, these sex-biased genes linked to a total of 61 male-biased DARs from set D1 and 54 female-biased DARs from set E1, respectively. We performed footprint analysis using TOBIAS to determine the predicted binding status of each TF motif instance in each DAR using the above-described bulk liver ATAC-seq data from 8 wk old male and female mice, analyzed separately for each sex. Motifs were included in the data output (Fig. 5) if they were annotated as bound at a DAR in at least 10% of either set of MASLD-linked sex-biased DARs or at the full sets of sex-biased DARs D1 or E1 in the bulk ATAC-seq data from at least one sex. For TF motifs from the same family, we retained the motif that gave the most binding events. Data are reported as average motif frequency values, calculated across the full DAR sets D1 and E1, or across the MASLD gene-linked DAR subsets described above.

### TF co-binding analysis

TF motifs enriched in the sets of sex-biased enhancer DARs linked to sex-biased genes (DAR sets D1 and E1), or in the sets of enhancer DARs linked to sex-biased genes whose hypox response class has been defined (DAR sets 1-4), were used as input for the CreateMotifMatrix function in Signac to select motifs with enrichment values >1 and FDR <0.05 and to generate a ATAC peak−TF motif matrix for each analysis. Each entry in this matrix indicates the presence/absence of the motif in a given feature. DARs with fewer than 5 motifs were filtered out. The matrix was analyzed by hierarchical clustering, where columns represent motifs and rows represent DARs. Row clustering was supervised, allowing the DARs to be clustered within their respective sets, and branch cutting was used to generate motif clusters and DAR clusters. In an alternative approach, we calculated Spearman correlation values between motifs based on their presence or absence in the DARs in each set. Motif pairs with a correlation value < 0.5 were filtered out. The remaining motif pairs were used to create a correlation heatmap, and hierarchical clustering was performed to identify motif clusters that were correlated. These motifs were then used to generate a DAR-motif matrix and heatmaps for each DAR set separately (Figs. S16-S17). Motif pairs and their correlation values and other information are presented in Table S6. We further defined motif co-occurrence events as motif pairs that belong to the same motif cluster group but did not belong to the same motif family, as determined by RSAT clustering.

### CellChat Analysis

We used the package CellChat [28], which inputs single cell-based gene expression data, assigns cell labels as input, and models cell-cell communication probabilities by integrating gene expression with prior knowledge of the interactions between signaling ligands, receptors, and their cofactors. snRNA-seq samples from livers of adult male and adult female, and cGH-treated adult male were provided as input along with cell labels extracted from the Seurat object. Next, we inferred cell-cell communication probability using the computeCommunProb function, which infers biologically significant interactions by assigning each interaction a probability value and performs a permutation test. We ran CellChat on each sample separately and then merged the different CellChat objects together using the mergeCellChat function. This resulted in a total of 152 interactions across biological conditions (M1, F1, F2, and M-cGH) (28 unique interactions, Fig. S18C), which were classified into 23 pathways, as defined by CellChatDB [28]. We identified conserved and condition-specific signaling pathways by comparing the information flow for each signaling pathway, which is defined by the sum of communication probability among all pairs of cell groups in the inferred network (i.e., the total weights in the network). Cell-Cell interaction networks were visualized using the circle plot layout in CellChat, with edge colors consistent with the cell sources, and edge weights being proportional to the interaction strength (thicker edge indicates stronger signal).

### Author contributions

KK and DJW were responsible for Project conception, and TyC for development and implementation of wet lab methods for nuclei isolation and nuclei qualification for 10X Genomics MultiOmics analysis. SG performed bulk mouse liver ATAC-seq wet lab analysis and MP performed the associated computational analysis. KK performed all of the other computational analysis, with some input from MP, except for motif occupancy analysis by ATAC-seq footprinting, which was carried out by TS. MP and DJW devised methods for generating cell type-specific DARs and matched background regions used for motif enrichment analysis, and MP performed the analyses in Figs. S20-S24. KK and DJW jointly prepared manuscript figures and Supplemental tables for publication. KK and DJW drafted the manuscript, and DJW revised and edited the manuscript for publication. DJW was responsible for supervision and overall project management. All authors approved of the final manuscript.

## Acknowledgements

The authors thank Hong Ma and Cameron Vergato for provision of frozen mouse liver tissue used in this study, and Samuel Mildrum and Stuart Levine (MIT BioMicro Center, Cambridge, MA) for preparation of MultiOmic ATAC + Gene Expression libraries and for consultation.

## Supplemental Figure Legends

**Fig. S1. Cell numbers and Cell type identification. A.** Venn diagram showing overlap of cells obtained from RNA modality using Dr. Maxim Pyatkov’s filtering approaches based on snRNA-seq data (’RNA only’) and ATAC-seq modality, where we applied filters to removed low quality ATAC-seq nuclei (‘ATAC only)’. The intersection shows nuclei that passed both RNA and ATAC filters and were used for downstream analysis. **B.** Dot plot showing average expression of marker genes (shown along X-axis) across the 5 liver cell clusters (Y-axis) shown in the UMAP in Fig. 1A based on liver nuclei isolated from control male and female liver and from male mice treated with cGH as an infusion over 12 d.

**Fig. S2. ATAC-seq peak annotation. A.** Venn diagram showing overlap of 127,957 (“127k”) ATAC-seq peaks with three mouse liver DHS sets from our laboratory. Overall, 74,860 of the 127,957 ATAC peaks (59%) overlap with a DHS identified in at least 1 of the 3 DHS peak sets. **B.** Distribution of ATAC-seq peaks annotated as promoter, enhancer, or insulator based on overlapping DHS sites annotated by Matthews et al. (2018). 54,923 of the 127,957 peaks overlapped with Set III in panel. Of these ATAC regions, 213 overlapped with two or more peaks from Set III that were a mix of enhancer, insulator, and promoter. As these 213 ATAC regions constitute a mixed group, they were removed from the pie chart. Percentages shown in the pie chart are this based on the 54,710 peaks common to the two sets (panel A). **C.** Distribution of the full set of 127,957 ATAC-seq peaks in each of the indicated chromatin states, based on our prior genomic annotations of chromatin state maps (14-state model) for male and female mouse liver [38]. **D.** Emission probabilities for six histone marks and DHS for each of the 14 chromatin states [38], and the grouping of the 14 states into the indicated superstates.

**Fig. S3. Flowchart of sex-biased, GH-responsive DARs and DEGs.** Flowchart shows the breakdown of ATAC-seq regions (N= 127,957) **(A)** and genes (N=58,275) **(B)** that are sex-biased and GH responsive or non-responsive, as determined by comparing: male vs. female replicate 1 (F1), male vs. female replicate 2 (F2), cGH-treated male vs. male (shown in Table S2C and Table S2D).

**Fig. S4. Flowchart of hypox response classified sex-biased genes.** Flowchart showing the breakdown of hypox responsive gene class (based on FDR<0.05 only) whose sex-bias expression was confirmed in our snRNA-seq dataset, and whose cGH responsiveness was also determined in our snRNA-seq dataset. We show number of sex-biased DARs that are linked to each hypox gene sets.

**Fig. S5. A.1 Flowchart of sex-biased DARs and their linkages to sex-biased genes.** Flowchart showing the breakdown of sex-biased DARs identified in Table S2C linked to sex-biased DEGs identified in Table S2D. See Table S3A, column BW for DAR listings. **Fig. S5 A.2.** Illustrative figure showing the six sets of DARs from Fig. S5A.1 that were used for motif analysis and other downstream analysis. **B. Average intensity.** Boxplots showing maximum ATAC-seq intensity values (*100) values (Table S1C, column BU) for the indicated DAR sets and background ATAC-seq peak sets used for motif analysis. Data are shown for enhancer state DARs after removing low activity DARs (i.e., lower 30% of ATAC-seq regions sorted by normalized maximum ATAC-seq intensity per nt, corresponding to only 1-2% of sex-biased DARs but a much higher percentage of background ATAC-seq regions; Table S3C). **C.** Plots showing maximum ATAC-seq intensity values (*100) (Table S1C, column BU) for the indicated DAR sets for all 127,957 ATAC-seq peaks (top) and for the top 89,600 ATAC-seq peaks (bottom) ranked based on normalized maximum hepatocyte ATAC peak intensities (Table S3C). **D.** Plots comparing the distribution of distances from each DAR to its nearest hepatocyte-expressed gene TSS. **C, D**. Significance based on Kruskal-Wallis test implemented in GraphPad Prism for pairwise comparisons between the indicated DAR sets, followed by Bonferroni FDR multiple comparison correction to obtain adjusted p-values, marked with asterisks, as follows: adj-p > 0.05 (ns), adj-p ≤ 0.05 (*), adj-p ≤ 0.05 (**), adj-p ≤ 0.01 (***), and adj-p ≤ 0.001 (****). See Table 2 of ‘Percentages’ Excel sheet for numbers of ATAC seq peaks in each of the DAR sets analyzed in panels B, C and D.

**Fig. S6. TAD distribution of sex-biased DARs linked to sex-biased genes. A-F,** TAD distributions shown as percentages and divided in two categories: Same TAD, i.e., the DAR and the DEG are in the same TAD region; and Different TAD, i.e., the DAR and the DEG are in different TADs. These distributions are shown for: **A,** Set D1: M-specific, cGH-repressed DARs linked to M-specific, cGH-repressed DEGs (N=408); **B,** Set D2a: M-specific, cGH repressed DARs linked non-sex-biased genes (N=650)**; C,** Set E1: F-specific, cGH induced DARs linked to F-specific, cGH induced DEGs (N=286); **D;** Set E2a: F-specific, cGH induced DARs linked non-sex-biased genes (N=278); **E,** Set D4: M-specific, cGH repressed DARs linked to other classes of sex-biased genes (N=87); and **F,** Set E4: F-specific, cGH induced DARs linked to other classes of sex-biased genes (N=48). **G.** Scatterplots comparing the distribution of distance from each DAR to its linked gene(s) (*left,* male-biased DARs; *right*, female-biased DARs) for DARs and their target genes in the same vs different TADs (y-axis). **H.** Scatterplots comparing the distribution of absolute correlation values between DARs and their linked genes, which provides an indication of the strength of the interaction. ATAC-seq−gene pairs with strong linkage correlations are considered to be “co-expressed” and enrich for a shared regulatory mechanism (https://support.10xgenomics.com/feature-linkage) for DARs and their target genes in the same vs different TADs (*left,* male sets, *right*, female sets). **I.** TAD distributions for non-sex-biased DARs linked to non-sex-biased genes (*left*), distribution of absolute correlation for their DAR-gene linkages (*middle*), and distribution of distances (*right*). Correlation analysis for panels G, H and I were performed using absolute correlation values for peak-gene interactions that were grouped into same TAD or different TAD based on their TAD assignments (see Methods). We used Kruskal-Wallis test to perform pairwise comparisons for each DAR set, followed by Bonferroni FDR multiple comparison tests to obtain adjusted p-values, summarized with asterix symbols, as follows: adj-p > 0.05 (ns), adj-p ≤ 0.05 (*), adj-p ≤ 0.05 (**), adj-p ≤ 0.01 (***), and adj-p ≤ 0.001 (****). Median values, horizontal lines on the correlation and distance plots.

**Fig. S7. Distribution of sex-biased DARs linked hypox class sex-biased genes.** TAD distributions for DARs linked to sex-biased genes in different hypox response classes, grouped into three categories: Same TAD (DAR and DEG in the same TAD), Different TAD (DAR and DEG are in different TADs), and Mixed (DARs linked to 2 or more DEGs, some in same TAD and other in a different TAD than the DAR). Distributions are shown for: **A,** Set D1 DARs linked to Set 1 (Class I M-specific cGH repressed genes) (N=227); **B,** Set D1 DARs linked to Set 2 (Class II M-specific cGH repressed genes) (N=52); **C;** Set E1 DARs linked to Set3 (Class I F-specific cGH induced genes) (N=128); and **D,** Set E1 DARs linked to Set4 (Class II F-specific cGH induced genes (N=62).

**Fig. S8. DAR-gene linkage correlations used to identify putative enhancer and repressor interactions.** Pie charts showing the number and percentage of DARs in each set that show positive correlations with their linked genes (marked as Enhancer), negative correlations with their linked genes (marked as Repressor), or show positive correlations with some linked genes and negative correlations with other linked genes (marked as Enhancer/Repressor).DAR sets analyzed: **A,** Set D1: M-specific, cGH repressed DARs linked to M-specific, cGH repressed DEGs (N=408); **B,**Set D2a: M-specific, cGH repressed DARs linked to non-sex-biased genes (N=650)**; C,** Set E1: F-specific, cGH induced DARs linked to F-specific, cGH induced DEGs (N=286); **D,** Set E2a: F-specific, cGH induced DARs linked to non-sex-biased genes (N=278); **E,** Set D4: M-specific, cGH repressed DARs linked to other classes of sex-biased genes (N=87); and **F,** Set E4: F-specific, cGH induced DARs linked to other classes of sex-biased genes (N=48).

**Fig. S9. A.** Bar plot showing enhancer and repressor distributions (as in Fig. S8) for sex-biased DARs linked to other classes of sex-biased genes (Set D4 and Set E4 from Fig. S5) grouped into subsets based on their linkages to genes with the same sex bias or the opposite sex bias, as detailed under the X-axis. **B, C,** Distributions of the indicated sets of sex-biased DARs based on whether they overlap a set II DHS peak (Fig. S2A) whose chromatin opening decreases (STAT5-Low), increases (STAT5-High), or is unchanged (STAT5-Static) in livers excised from individual male mice euthanized at the time when liver STAT5 activity is high (due to a recent plasma GH pulse), as compared to the activity in livers excised from mice euthanized at the time when liver STAT5 activity is high (time period between plasma GH pulses). In panel B, 2,616 out of 3,518 M-specific DARs and 698 out of 1,230 F-specific DARs overlapped with DHS peaks from set III shown in Fig. 2A. Seven out of 2,616 DARs and 2 out of 698 DARs overlapped with multiple peaks annotated as a mix of static or stat5_hi/lo. We labelled these DARs as ‘mixed group’ and removed them from the bar plot. In panel C, percentages are calculated based on DARs that overlapped with set III DHS and did not belong to mixed group (described above). Set D1 (N=408): 324 DARs, Set D2a (N=650): 529 DARs, Set D3 (N=2,373): 1682 DARs, Set D4 (N=87): 74 DARs, Set E1 (N=286): 183 DARs, Set E2a (N=278): 186 DARs, Set E3 (N=618): 287 DARs, Set E4 (N=48): 40 DARs.

**Fig. S10.** Chromatin state model, with states E1-E14 and emission parameters as shown. This model was defined using a series of 6 informative histone marks, and DNase-I hypersensitivity data collected for adult male liver, and separately, for adult female liver. The genome-wide chromatin state models obtained encompass sex differences in chromatin state and chromatin structure and their relationships to sex-biased gene expression [38].

**Fig. S11. Chromatin state analysis of DAR regions D1, E1, D2a, and E2a.**

**A-D** Chromatin state distributions of the indicated sets of DARs linked to sex-biased genes **(A, C)** or linked to non-sex-biased genes **(B, D)**, based on the 14-chromatin state model in Fig.S10.

**Fig. S12. Chromatin state analysis of DAR regions D3, E3, D4, and E4.**

**A-D** Chromatin state distributions of sex-biased DARs linked to other classes of sex-biased genes **(A, C)** or with no gene linkages **(B, D)**, based on the 14-chromatin state model.

**Fig. S13. A. Functional enrichment analysis of DAR gene targets.** Genes that were linked to DARs from DAR sets D1, E1, D2a and E2a were used as input to Metascape to perform multi-list comparative analysis and identify pathways that are shared between or are specific to each of these four DAR-linked gene sets. The scale represents -log10 P-value of top enriched pathways across the four gene lists, with darker purple coloration indicating more highly significant p-values. Enriched functional groups are shown at the *right*. See Table *Metascape* for details. **B.** Venn diagrams showing overlap of cell type-specific motifs discovered using four different approaches: Differential activity analysis using chromVAR based on the sets of 127,957 and 89,600 ATAC-seq peaks, and motif enrichment analysis using cell type-specific DARs for distal and promoter ATAC-seq peaks.

**Fig. S14. Motif z-score heatmaps. A.** Scaled heatmap showing -log10 FDR values for 432 TF motifs that are enriched in DARs that are linked to different set of genes (D1, D2a, D3, E1, E2a, E3, as defined in Fig. S5). **B.** Scaled heatmap data for 364 motifs that are enriched for DARs that are linked to hypox class sex-biased genes (sets 1-4, defined in Fig. S4). See Table S4F for full details.

**Fig. S15. Non-redundant motif co-occurrence at sex-biased DARs (A) and at hypox class-defined sex-biased DARs (B).** Data is related to Fig. 5A and Fig. 5B, except that the data shown here are non-redundant motif co-occurrence heatmaps based on *representative* motifs (lowest FDR value) from each motif cluster. In (**A**), a single representative motif was selected for each motif family as the motif with the lowest FDR value in Table S4D between set D1 and set E1 DARs. In (**B**), a single representative motif was selected for each motif family as the motif with the lowest FDR value in Table S4E across DAR sets 1, 2, 3 and 4.

**Fig. S16. Motif co-correlations with correlation heatmaps for Set D1 and Set E1 DARs (A, B) and for DARs linked to hypox class defined sex-biased genes (C, D). (Left)** Heatmap showing Spearman correlation values >0.5 computed for each motif pair for a set of 154 motifs found to be enriched in Set D1 **(A)** and calculated across the full set of DARs in Set D1. Off diagonal values indicate a significant correlation between 2 motifs. **(B)** the same analysis described in (A) was performed for 30 motifs enriched in set E1 DARs. Hierarchical clustering was performed to identify motif clusters that were correlated. Accordingly, motifs that are nearby along across the top are most highly correlated and tend to be redundant or near redundant motifs from the same gene families. **(Right)** The correlated motifs were used to generate DAR-motif for each set separately. Column clustering was supervised with motifs clustered within their cluster groups as identified from the correlation heatmap. Column annotation also show motif clusters identified by the RSAT clustering tool. Row annotations show DAR clusters and hypox response class. Row clustering in the heatmap was supervised to ensure that the DARs remained clustered within their hypox class. **(C, D)** Analysis is the same as described in (A) and (B), except that the heatmap (*left*) is based on 297 motifs in set 1 and 98 motifs in set 2 and the analysis shown is for DARs linked to hypox response class male-biased gene set 1 **(C)** and set 2 **(D)** (described in Fig. S4).

**Fig. S17. Motif co-binding with correlation heatmaps for DARs linked to hypox class female-biased genes.** Analysis is the same as described in Fig. S16, except that the heatmap (*left*) is based on 109 motifs in set 3 and 45 motifs in set 4 and the analysis shown for DARs linked to hypox response class female-biased gene set 3 **(A)** and set 4 **(B)** (described in Fig. S4).

**Fig. S18. Cell chat interactions**. **A.** Bar plot showing number of ligand-receptor interactions identified by CellChat in snRNA-seq cell clusters for each of the four snRNA-seq samples (male (M), female replicate 1 (F1), female replicate 2 (F2), and cGH-treated male. **B.** Ligand-receptor interactions identified in each dataset are grouped into different signaling pathways. X-axis shows the relative information flow of each signaling pathway (named on Y-axis), indicating signaling pathways that are either turned on (i.e., increased) or turned off (i.e., decreased) between different samples. **C.** Venn diagram showing overlap of ligand receptor interactions discovered from CellChat between liver samples M and F1, M and F2, and M and M-cGH.

**Fig. S19. Nuclei isolation.** Microscopic image of high-quality nuclei extracted from male and female mouse liver and used for preparation of the 10X Genomics multiOmic libraries. Nuclei quality is indicated by the presence of an intact nuclear membrane without peripheral blebbing.

**Fig. S20. Filtering contaminated nuclei in male liver snRNA-seq sample. A.** Intronic UMI vs. exonic UMI plot for snRNA-seq of male liver shows two distinct population of nuclei (populations I and II), separated from each other based on a support vector machine (svm) model. The lower Intronic to Exonic UMI ratio of the population II nuclei is consistent with their being contaminated by internal cell membranes (nucleus-bound endoplasmic reticulum) [39], which are enriched in the mature (spliced) RNAs for secretory RNAs that are particularly abundant in hepatocytes. **B. (Top)** UMAP of male snRNA-seq nuclei using graph-based clustering in Seurat and visualized using UMAP. **(Middle)** Clustered cells from UMAP above were projected on to the Intronic vs Exonic UMI plot to determine their distribution based on the two populations. **(Bottom)** Dotplot showing the cell type labeling of each UMAP cluster based on known marker genes across the X-axis. **C. (Top)** UMAP of the same male sample of snRNA-seq nuclei after removing population II nuclei, followed by graph-based clustering in Seurat and visualization in a UMAP. **(Middle)** Clustered cells (without population II) from the above projected on to the Intronic vs Exonic UMI plot. **(Bottom)** Dotplot for cell type labeling using known marker genes, as in **B**. The dotplot shows greater specificity of the hepatocyte marker genes for hepatocyte clusters and more effective removal of mitochondrial genes after removing population II nuclei.

**Fig. S21, Fig. S22, and Fig. S23. Filtering of contaminated nuclei from female (F1) liver, female (F2) liver and cGH-treated male liver, respectively.** Data display is the same as in Fig. S20.

**Fig. S24. A. (Top)** UMAP of aggregated cells from four samples (male, female F1, female F2 and cGH-treated male using whole gene body counting method. **(Bottom)** Dot plots of known marker genes for UMAP with gene body counting. **B. (Top)** UMAP of aggregated cells from the four samples using the modified intronic counting method (Intronic + Mono-exonic counting). **(Bottom)** Dot plots of known marker genes for UMAP with modified intronic counting. Results show that by using the Intronic + Mono-exonic counting method (*right*), which excludes exonic UMIs for all multi-exonic genes, including those of hepatocyte secretory protein RNAs, there is a substantial decrease in the apparent contamination of the non-hepatocyte clusters by hepatocyte marker gene RNAs (Alb, Ttr, Apoa1, Serpina1c) as compared to when using the whole gene body counting method (*left*).

## Abbreviations

lncRNA: long non-coding RNA
scRNA-seq: single cell RNA sequencing
UMAP: uniformed manifold approximation and projection
ATAC: assay for transposase-accessible chromatin
DAR: differential accessible chromatin region(s)
cGH: continuous growth hormone
Hypox: hypophysectomy
TF: transcription factor

