## Supplemental Figures for "Single Nucleus MultiOmics Links Novel Transcription Factor Motifs to Murine Hepatic Sex Differences in Chromatin Accessibility and Metabolic Dysfunction-Associated Steatotic Liver Disease"

### Fig. S1

A.

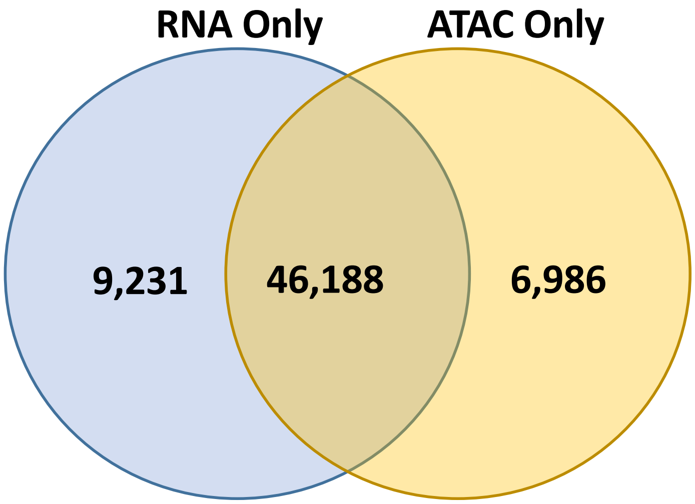
RNA Only
ATAC Only
9,231
46,188
6,986

B.

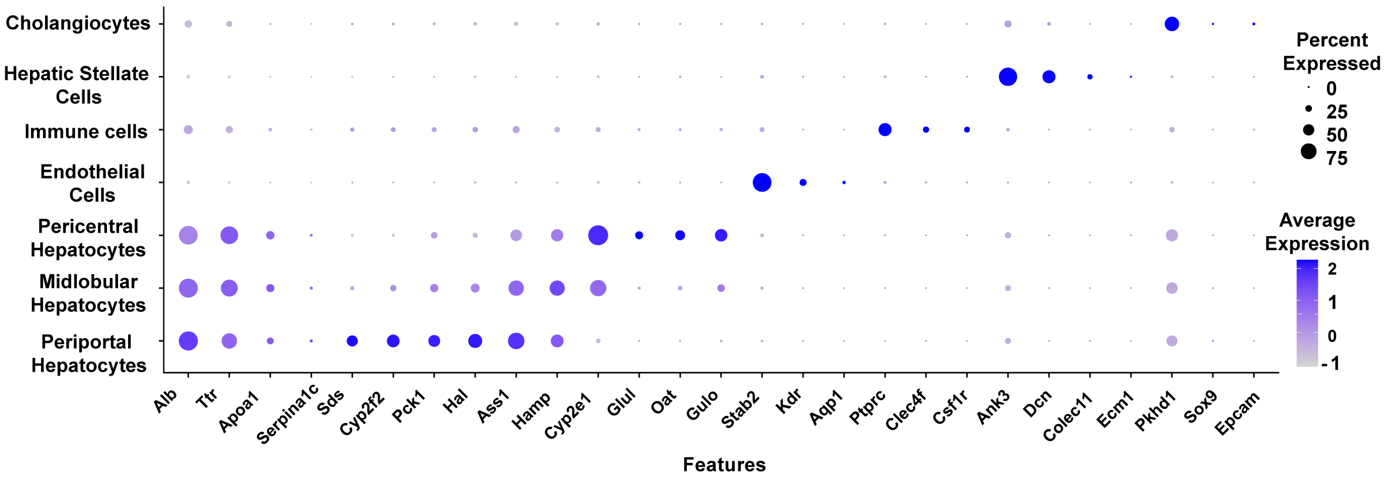
Cholangiocytes
Hepatic Stellate Cells
Immune cells
Endothelial Cells
Pericentral Hepatocytes
Midlobular Hepatocytes
Periportal Hepatocytes
Alb
Ttr
Apoa1
Serpina1c
Sds
Cyp2f2
Pck1
Hal
Ass1
Hamp
Cyp2e1
Glul
Oat
Gulo
Stab2
Kdr
Aqp1
Ptprc
Clec4f
Csf1r
Ank3
Dcn
Colec11
Ecm1
Pkhd1
Sox9
Epcam
Features
Percent Expressed
0
25
50
75
Average Expression
2
1
0
-1

**Fig. S2**

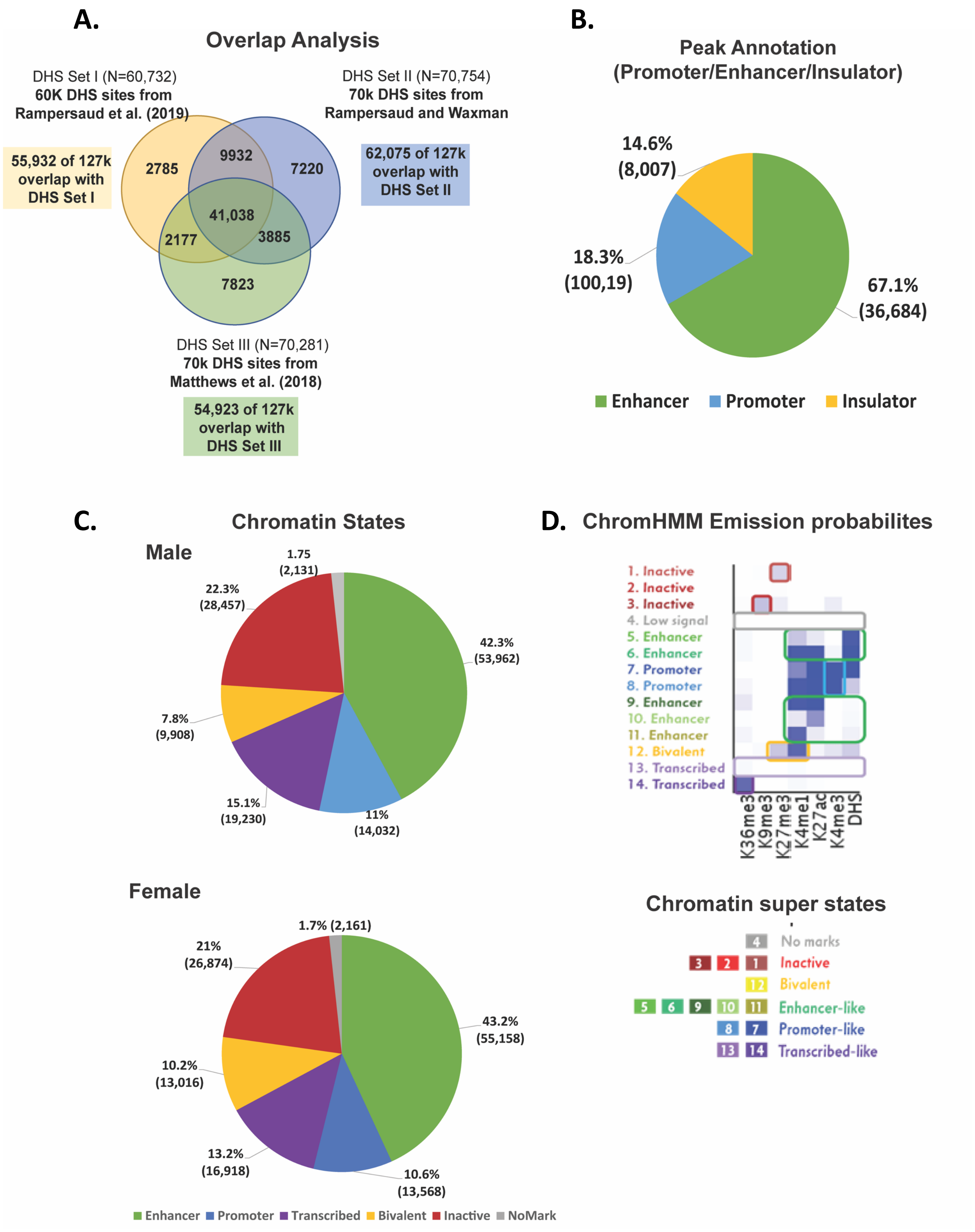
A.
Overlap Analysis
DHS Set I (N=60,732)
60K DHS sites from
Rampersaud et al. (2019)
DHS Set II (N=70,754)
70k DHS sites from
Rampersaud and Waxman
55,932 of 127k
overlap with
DHS Set I
62,075 of 127k
overlap with
DHS Set II
2785
9932
7220
41,038
2177
3885
7823
DHS Set III (N=70,281)
70k DHS sites from
Matthews et al. (2018)
54,923 of 127k
overlap with
DHS Set III
B.
Peak Annotation
(Promoter/Enhancer/Insulator)
14.6%
(8,007)
18.3%
(100,19)
67.1%
(36,684)
Enhancer
Promoter
Insulator
C.
Chromatin States
Male
1.75
(2,131)
22.3%
(28,457)
42.3%
(53,962)
7.8%
(9,908)
15.1%
(19,230)
11%
(14,032)
Female
1.7% (2,161)
21%
(26,874)
43.2%
(55,158)
10.2%
(13,016)
13.2%
(16,918)
10.6%
(13,568)
Enhancer
Promoter
Transcribed
Bivalent
Inactive
NoMark
D.
ChromHMM Emission probabilites
1. Inactive
2. Inactive
3. Inactive
4. Low signal
5. Enhancer
6. Enhancer
7. Promoter
8. Promoter
9. Enhancer
10. Enhancer
11. Enhancer
12. Bivalent
13. Transcribed
14. Transcribed
K36me3
K9me3
K27me3
K4me1
K27ac
K4me3
DHS
Chromatin super states
4 No marks
3 2 1 Inactive
12 Bivalent
5 6 9 10 11 Enhancer-like
8 7 Promoter-like
13 14 Transcribed-like

### Fig. S3

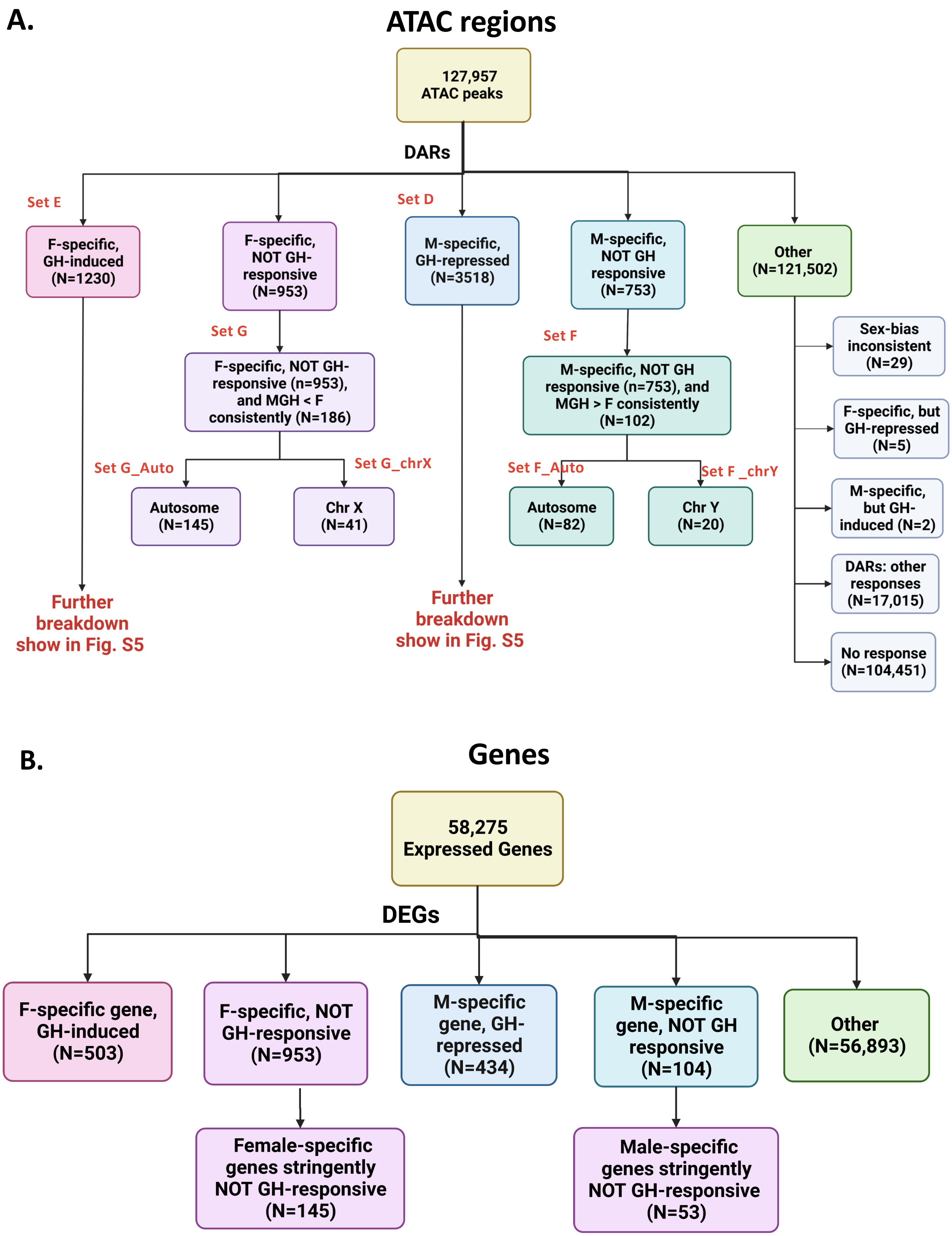
A.
ATAC regions
127,957 ATAC peaks
DARs
Set E
F-specific, GH-induced (N=1230)
F-specific, NOT GH-responsive (N=953)
Set D
M-specific, GH-repressed (N=3518)
M-specific, NOT GH responsive (N=753)
Other (N=121,502)
Set G
F-specific, NOT GH-responsive (n=953), and MGH < F consistently (N=186)
Set F
M-specific, NOT GH responsive (n=753), and MGH > F consistently (N=102)
Set G_Auto
Set G_chrX
Autosome (N=145)
Chr X (N=41)
Set F_Auto
Set F _chrY
Autosome (N=82)
Chr Y (N=20)
Sex-bias inconsistent (N=29)
F-specific, but GH-repressed (N=5)
M-specific, but GH-induced (N=2)
DARs: other responses (N=17,015)
No response (N=104,451)
Further breakdown show in Fig. S5
Further breakdown show in Fig. S5
B.
Genes
58,275 Expressed Genes
DEGs
F-specific gene, GH-induced (N=503)
F-specific, NOT GH-responsive (N=953)
M-specific gene, GH-repressed (N=434)
M-specific gene, NOT GH responsive (N=104)
Other (N=56,893)
Female-specific genes stringently NOT GH-responsive (N=145)
Male-specific genes stringently NOT GH-responsive (N=53)

Fig. S4

#### Hypox genes

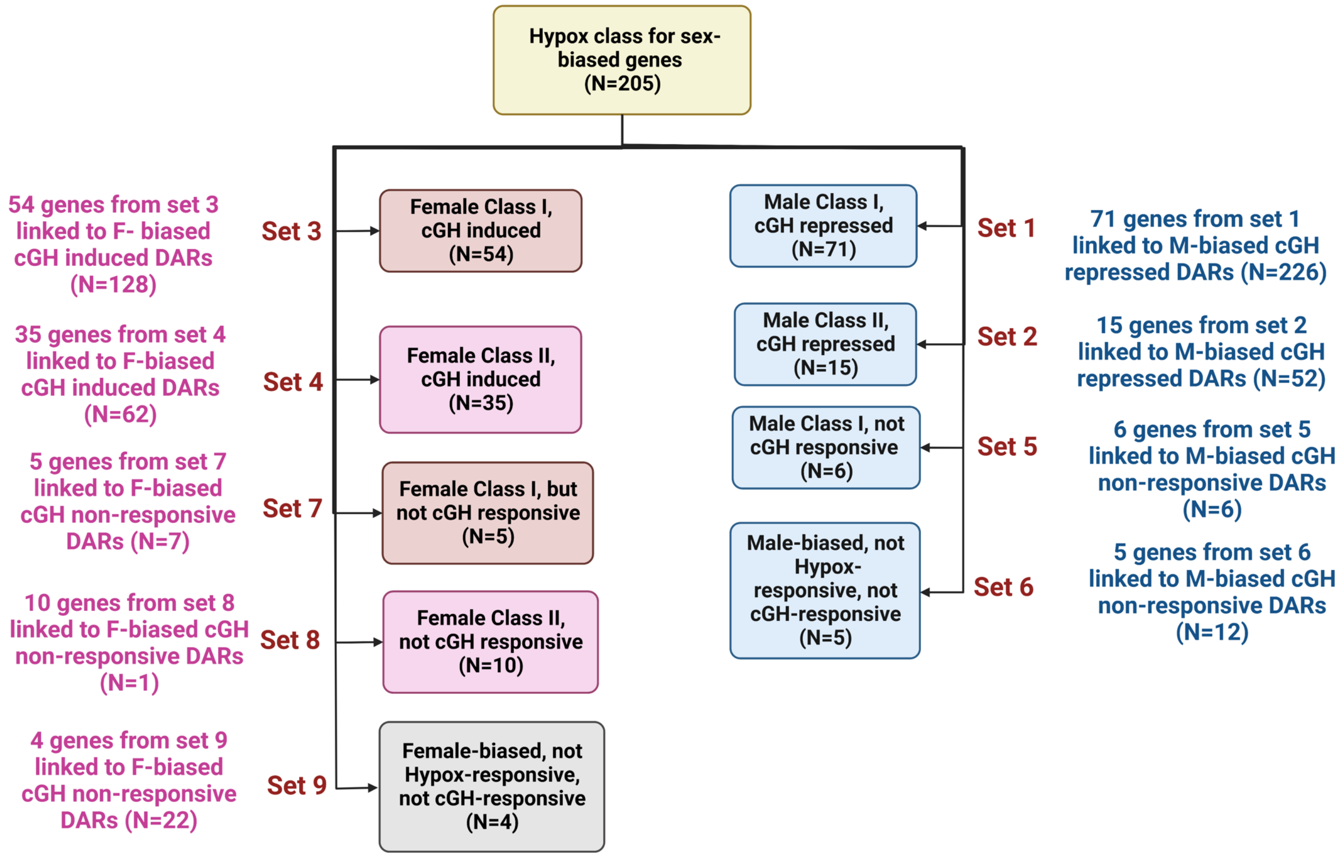

### Fig. S5A.1

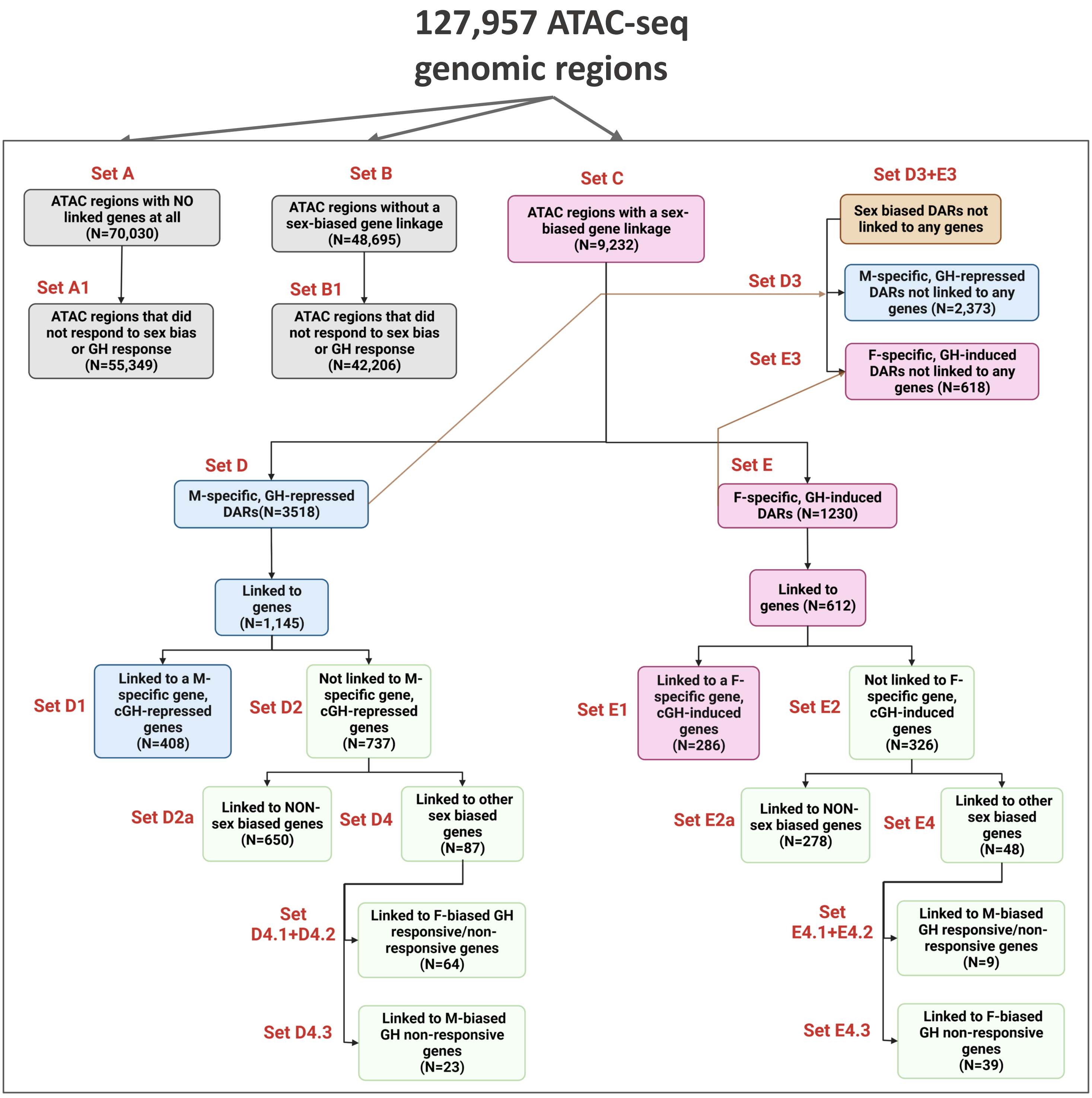

127,957 ATAC-seq genomic regions
Set A
ATAC regions with NO linked genes at all (N=70,030)
Set A1
ATAC regions that did not respond to sex bias or GH response (N=55,349)
Set B
ATAC regions without a sex-biased gene linkage (N=48,695)
Set B1
ATAC regions that did not respond to sex bias or GH response (N=42,206)
Set C
ATAC regions with a sex-biased gene linkage (N=9,232)
Set D3+E3
Sex biased DARs not linked to any genes
Set D3
M-specific, GH-repressed DARs not linked to any genes (N=2,373)
Set E3
F-specific, GH-induced DARs not linked to any genes (N=618)
Set D
M-specific, GH-repressed DARs(N=3518)
Set E
F-specific, GH-induced DARs (N=1230)
Linked to genes (N=1,145)
Linked to genes (N=612)
Set D1
Linked to a M-specific gene, cGH-repressed genes (N=408)
Set D2
Not linked to M-specific gene, cGH-repressed genes (N=737)
Set E1
Linked to a F-specific gene, cGH-induced genes (N=286)
Set E2
Not linked to F-specific gene, cGH-induced genes (N=326)
Set D2a
Linked to NON-sex biased genes (N=650)
Set D4
Linked to other sex biased genes (N=87)
Set E2a
Linked to NON-sex biased genes (N=278)
Set E4
Linked to other sex biased genes (N=48)
Set D4.1+D4.2
Linked to F-biased GH responsive/non-responsive genes (N=64)
Set D4.3
Linked to M-biased GH non-responsive genes (N=23)
Set E4.1+E4.2
Linked to M-biased GH responsive/non-responsive genes (N=9)
Set E4.3
Linked to F-biased GH non-responsive genes (N=39)

### Fig. S5A.2

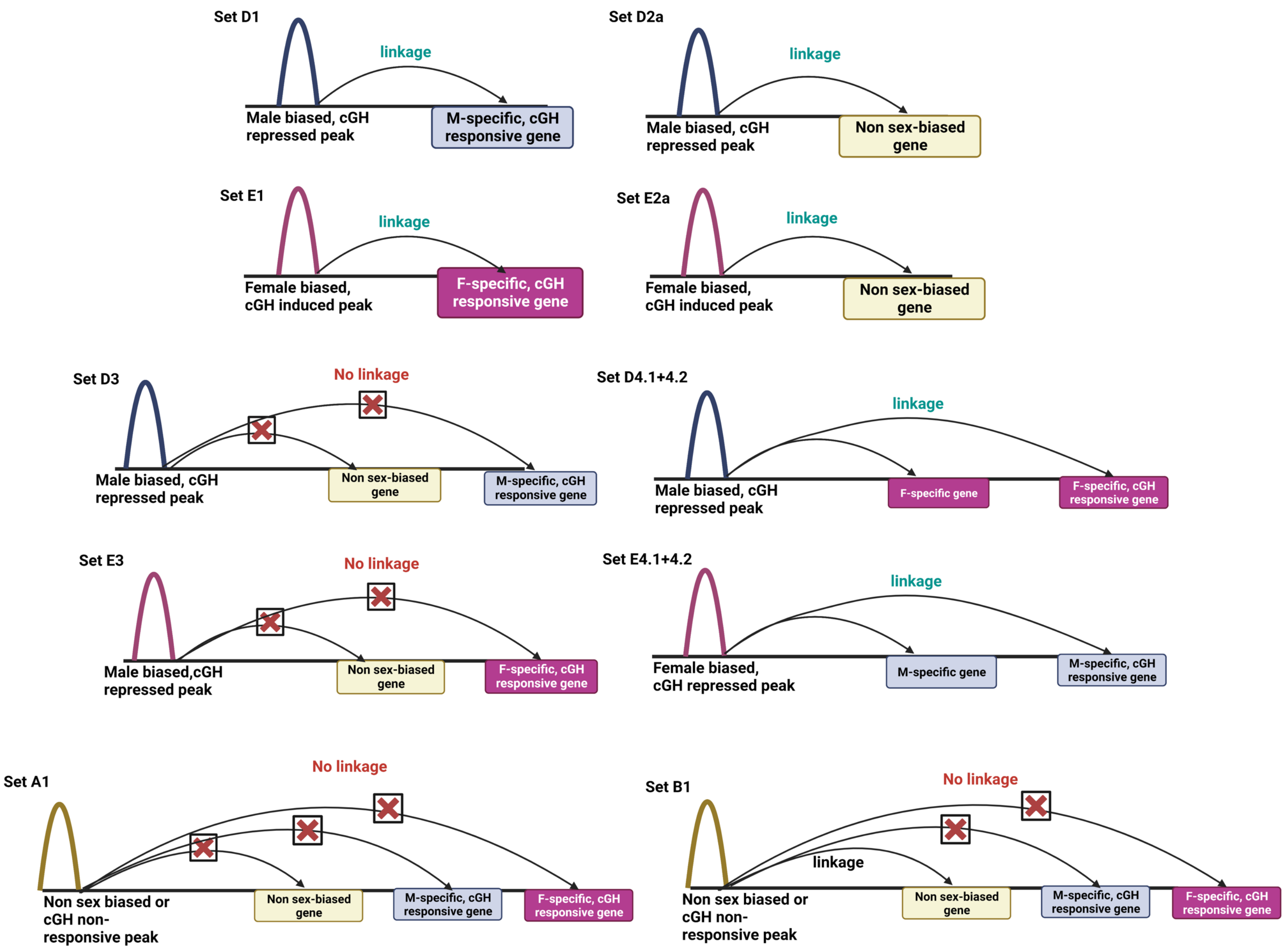
Set D1
linkage
Male biased, cGH repressed peak
M-specific, cGH responsive gene
Set D2a
linkage
Male biased, cGH repressed peak
Non sex-biased gene
Set E1
linkage
Female biased, cGH induced peak
F-specific, cGH responsive gene
Set E2a
linkage
Female biased, cGH induced peak
Non sex-biased gene
Set D3
No linkage
Male biased, cGH repressed peak
Non sex-biased gene
M-specific, cGH responsive gene
Set D4.1+4.2
linkage
Male biased, cGH repressed peak
F-specific gene
F-specific, cGH responsive gene
Set E3
No linkage
Male biased,cGH repressed peak
Non sex-biased gene
F-specific, cGH responsive gene
Set E4.1+4.2
linkage
Female biased, cGH repressed peak
M-specific gene
M-specific, cGH responsive gene
Set A1
No linkage
Non sex biased or cGH non-responsive peak
Non sex-biased gene
M-specific, cGH responsive gene
F-specific, cGH responsive gene
Set B1
No linkage
linkage
Non sex biased or cGH non-responsive peak
Non sex-biased gene
M-specific, cGH responsive gene
F-specific, cGH responsive gene

Fig. S5B

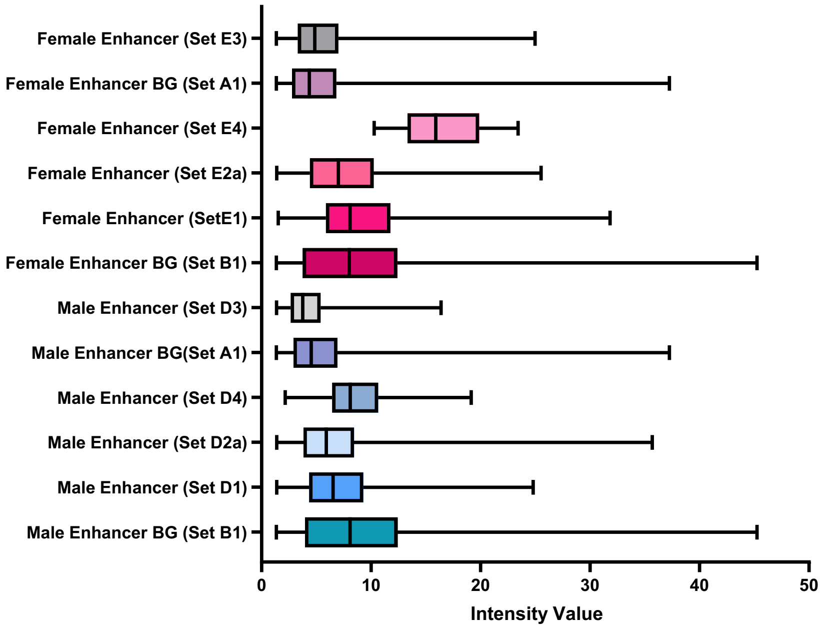


| Sets | Peaks in enhancer state | Total number of peaks | Median intensity values |
|---|---|---|---|
| **Male BG Set B1** | 12,051 | 42,206 | 8.07 |
| **SetD1** | 308 | 408 | 6.53 |
| **SetD2a** | 485 | 650 | 5.92 |
| **SetD4** | 48 | 64 | 8.09 |
| **Male BG Set A1** | 20,675 | 55,349 | 4.53 |
| **SetD3** | 1402 | 2373 | 3.76 |
| **Female BG SetB1** | 12,270 | 42,206 | 8.02 |
| **SetE1** | 211 | 286 | 8.07 |
| **SetE2a** | 206 | 278 | 7.02 |
| **SetE4** | 6 | 9 | 15.91 |
| **Female BG SetA1** | 21,411 | 55,349 | 4.37 |
| **SetE3** | 436 | 618 | 4.87 |

Fig. S5C

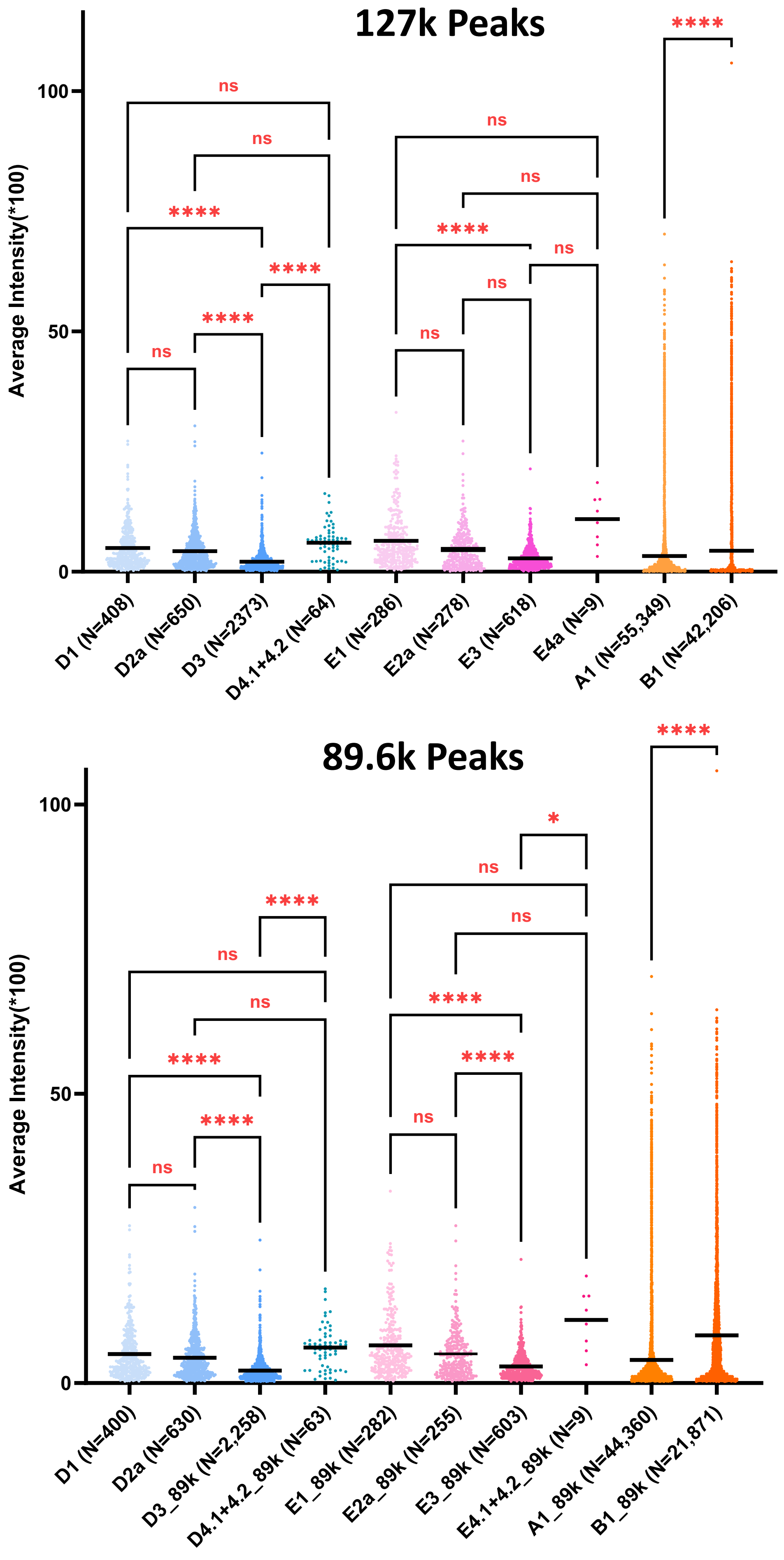
127k Peaks
Average Intensity(*100)
100
50
0
ns
ns
****
****
****
ns
ns
ns
****
ns
ns
ns
****
D1 (N=408)
D2a (N=650)
D3 (N=2373)
D4.1+4.2 (N=64)
E1 (N=286)
E2a (N=278)
E3 (N=618)
E4a (N=9)
A1 (N=55,349)
B1 (N=42,206)
89.6k Peaks
Average Intensity(*100)
100
50
0
****
ns
ns
****
****
ns
*
ns
ns
****
****
ns
****
D1 (N=400)
D2a (N=630)
D3_89k (N=2,258)
D4.1+4.2_89k (N=63)
E1_89k (N=282)
E2a_89k (N=255)
E3_89k (N=603)
E4.1+4.2_89k (N=9)
A1_89k (N=44,360)
B1_89k (N=21,871)

### Fig. S5D

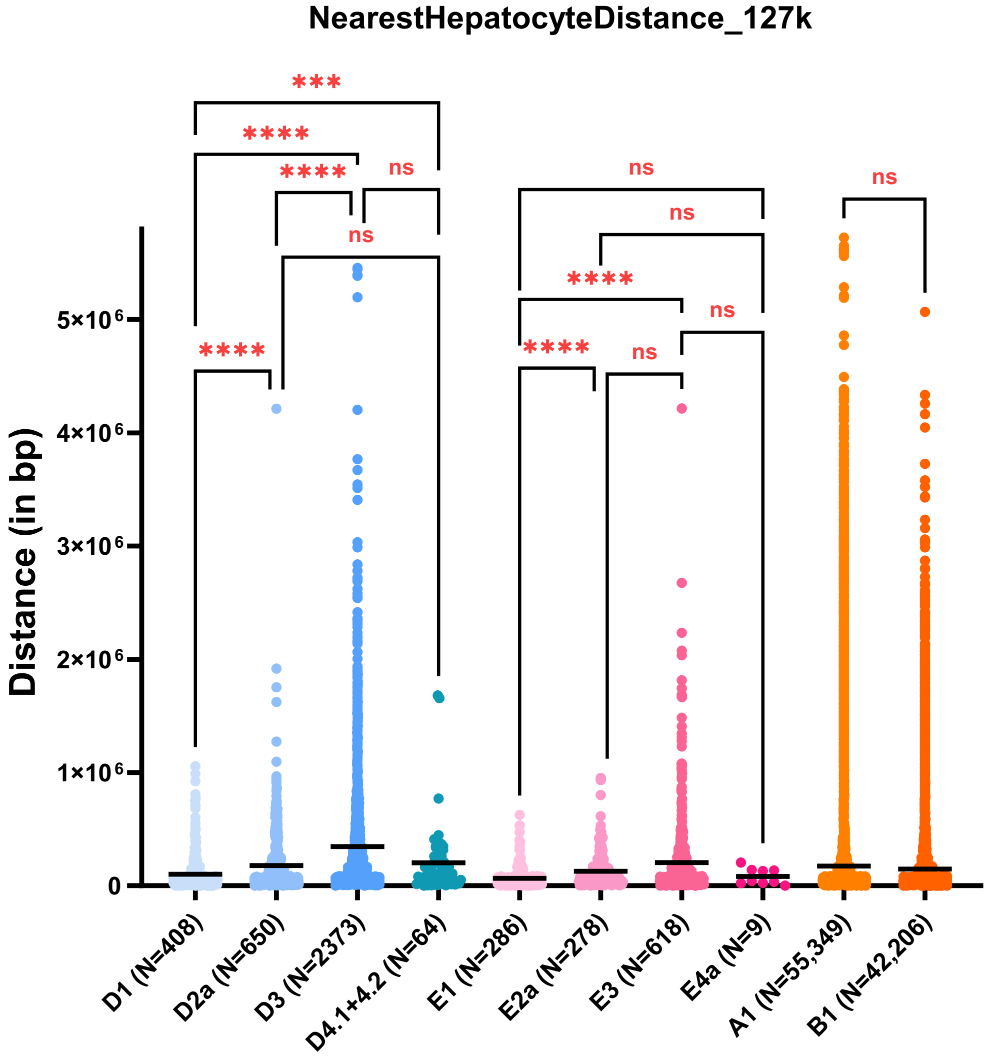


**Median Distance**

| D1 (N=408) | D2a (N=650) | D3 (N=2373) | D4.1+4.2 (N=64) | E1 (N=286) | E2a (N=278) | E3 (N=618) | E4a (N=9) | A1 (N=55,349) | B1 (N=42,206) |
|---|---|---|---|---|---|---|---|---|---|
| 39,673.5 | 81,301 | 143,460 | 104,648 | 32,577 | 76,513 | 94,912 | 42,055 | 65,461 | 69,035 |

Fig. S6

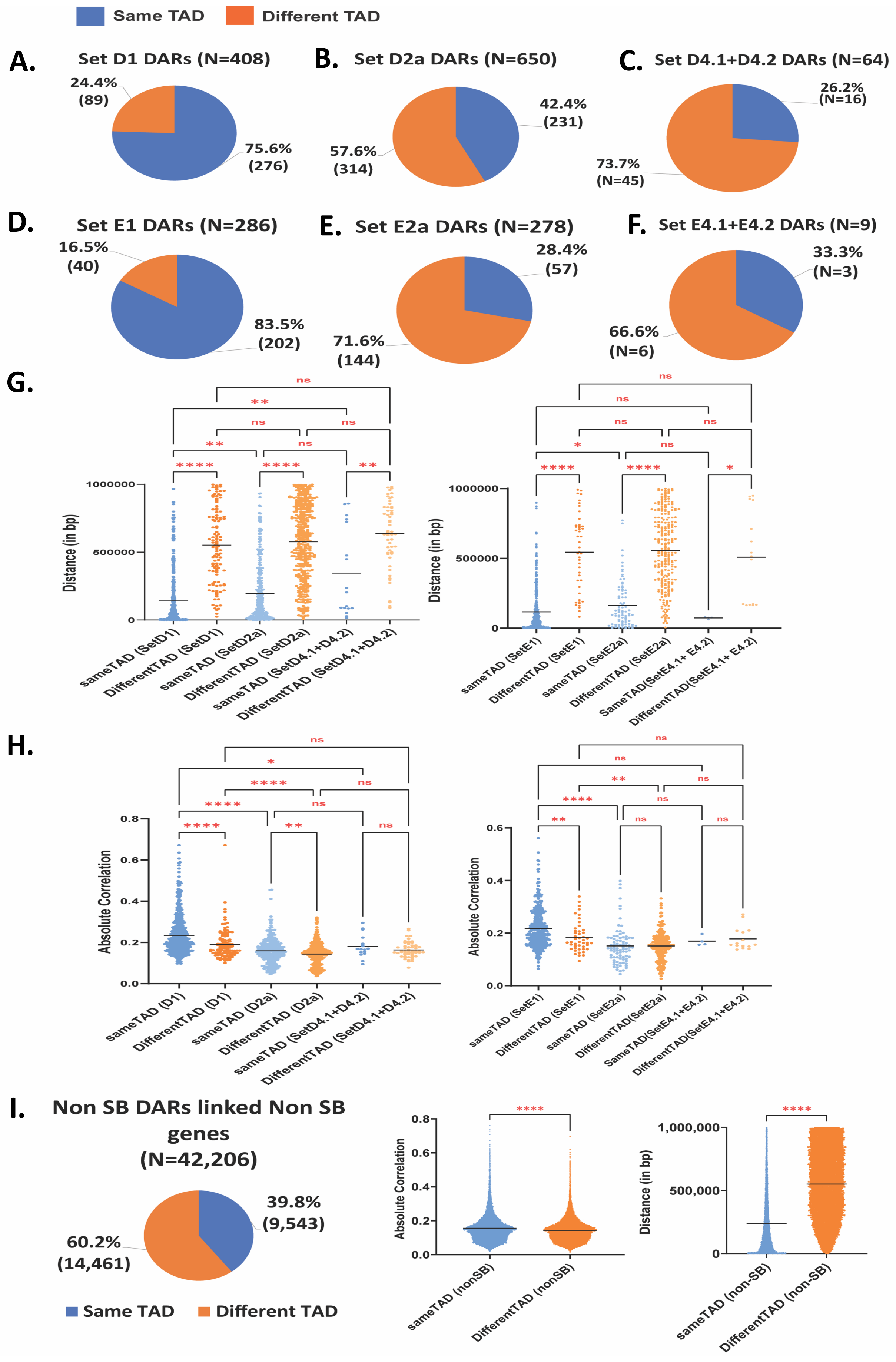

Same TAD
Different TAD
A.
Set D1 DARs (N=408)
24.4% (89)
75.6% (276)
B.
Set D2a DARs (N=650)
42.4% (231)
57.6% (314)
C.
Set D4.1+D4.2 DARs (N=64)
26.2% (N=16)
73.7% (N=45)
D.
Set E1 DARs (N=286)
16.5% (40)
83.5% (202)
E.
Set E2a DARs (N=278)
28.4% (57)
71.6% (144)
F.
Set E4.1+E4.2 DARs (N=9)
33.3% (N=3)
66.6% (N=6)
G.
Distance (in bp)
sameTAD (SetD1)
DifferentTAD (SetD1)
sameTAD (SetD2a)
DifferentTAD (SetD2a)
sameTAD (SetD4.1+D4.2)
DifferentTAD (SetD4.1+D4.2)
sameTAD (SetE1)
DifferentTAD (SetE1)
sameTAD (SetE2a)
DifferentTAD (SetE2a)
SameTAD(SetE4.1+ E4.2)
DifferentTAD(SetE4.1+ E4.2)
H.
Absolute Correlation
sameTAD (D1)
DifferentTAD (D1)
sameTAD (D2a)
DifferentTAD (D2a)
sameTAD (SetD4.1+D4.2)
DifferentTAD (SetD4.1+D4.2)
sameTAD (SetE1)
DifferentTAD (SetE1)
sameTAD (SetE2a)
DifferentTAD(SetE2a)
SameTAD(SetE4.1+E4.2)
DifferentTAD(SetE4.1+E4.2)
I.
Non SB DARs linked Non SB genes (N=42,206)
39.8% (9,543)
60.2% (14,461)
Same TAD
Different TAD
Absolute Correlation
sameTAD (nonSB)
DifferentTAD (nonSB)
Distance (in bp)
sameTAD (non-SB)
DifferentTAD (non-SB)

**Fig. S7**

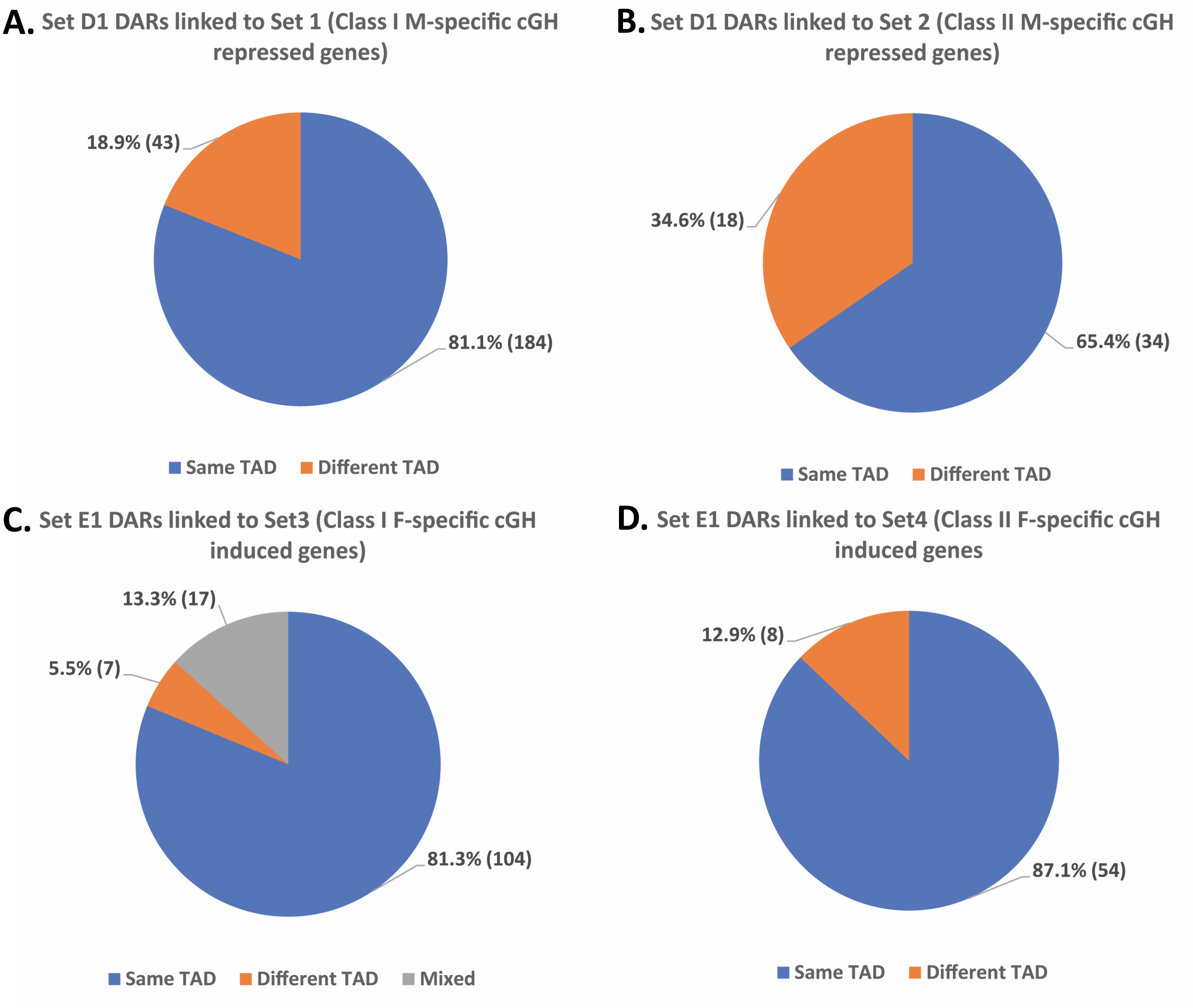
A. Set D1 DARs linked to Set 1 (Class I M-specific cGH repressed genes)
18.9% (43)
81.1% (184)
Same TAD
Different TAD
B. Set D1 DARs linked to Set 2 (Class II M-specific cGH repressed genes)
34.6% (18)
65.4% (34)
Same TAD
Different TAD
C. Set E1 DARs linked to Set3 (Class I F-specific cGH induced genes)
13.3% (17)
5.5% (7)
81.3% (104)
Same TAD
Different TAD
Mixed
D. Set E1 DARs linked to Set4 (Class II F-specific cGH induced genes
12.9% (8)
87.1% (54)
Same TAD
Different TAD

**Fig. S8**

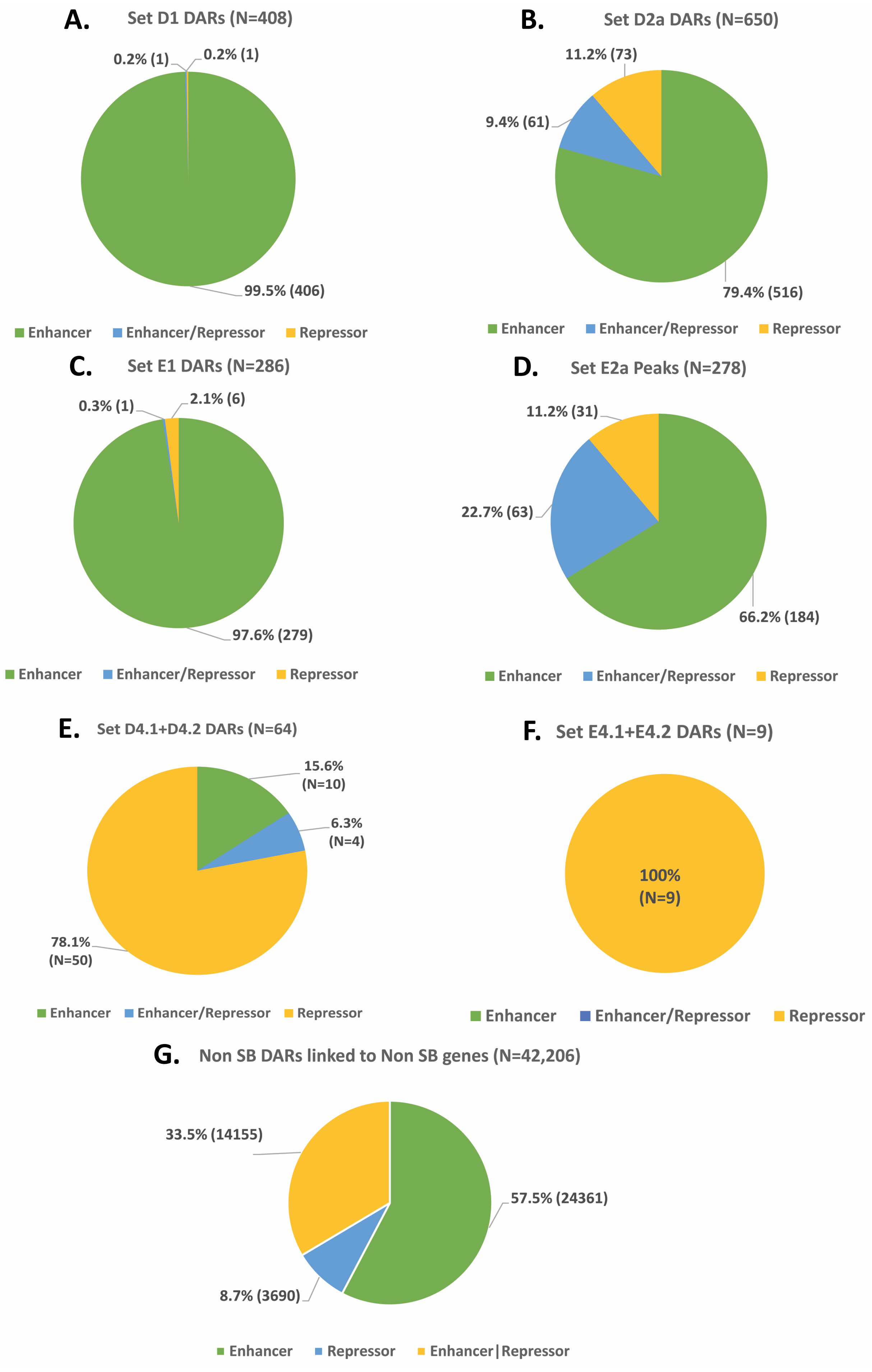
A.
Set D1 DARs (N=408)
0.2% (1)
0.2% (1)
99.5% (406)
Enhancer
Enhancer/Repressor
Repressor
B.
Set D2a DARs (N=650)
11.2% (73)
9.4% (61)
79.4% (516)
Enhancer
Enhancer/Repressor
Repressor
C.
Set E1 DARs (N=286)
0.3% (1)
2.1% (6)
97.6% (279)
Enhancer
Enhancer/Repressor
Repressor
D.
Set E2a Peaks (N=278)
11.2% (31)
22.7% (63)
66.2% (184)
Enhancer
Enhancer/Repressor
Repressor
E.
Set D4.1+D4.2 DARs (N=64)
15.6% (N=10)
6.3% (N=4)
78.1% (N=50)
Enhancer
Enhancer/Repressor
Repressor
F.
Set E4.1+E4.2 DARs (N=9)
100% (N=9)
Enhancer
Enhancer/Repressor
Repressor
G.
Non SB DARs linked to Non SB genes (N=42,206)
33.5% (14155)
57.5% (24361)
8.7% (3690)
Enhancer
Repressor
Enhancer|Repressor

**Fig. S9**

**A.**

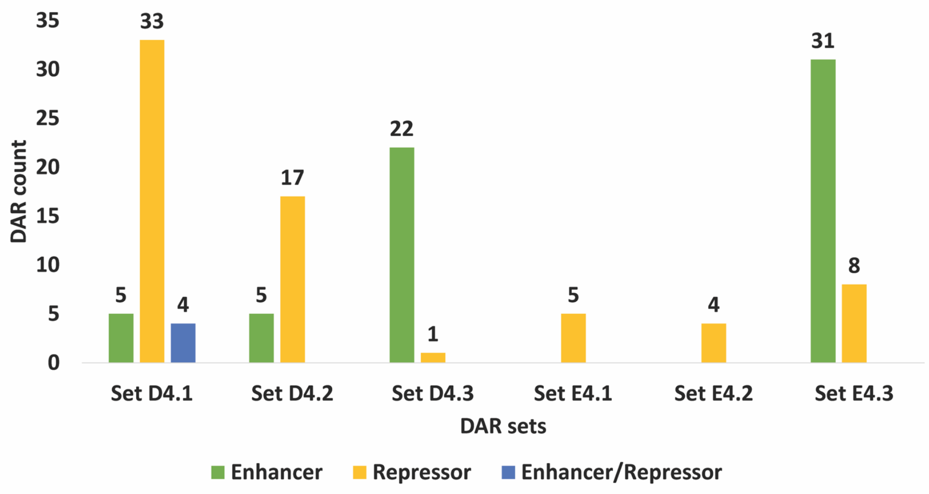


Set D4.1: Male biased DAR linked to F-specific gene, GH-induced gene
Set D4.2: Male biased DAR linked to F-specific gene, NOT GH-responsive gene
Set D4.3: Male biased DAR linked to M-specific gene, NOT GH responsive gene
Set E4.1: Female biased DAR linked to M-specific,GH-repressed gene
Set E4.2: Female biased DAR linked to M-specific, NOT GH responsive gene
Set E4.3: Female biased DAR linked to F-specific, NOT GH-responsive gene

**B.**

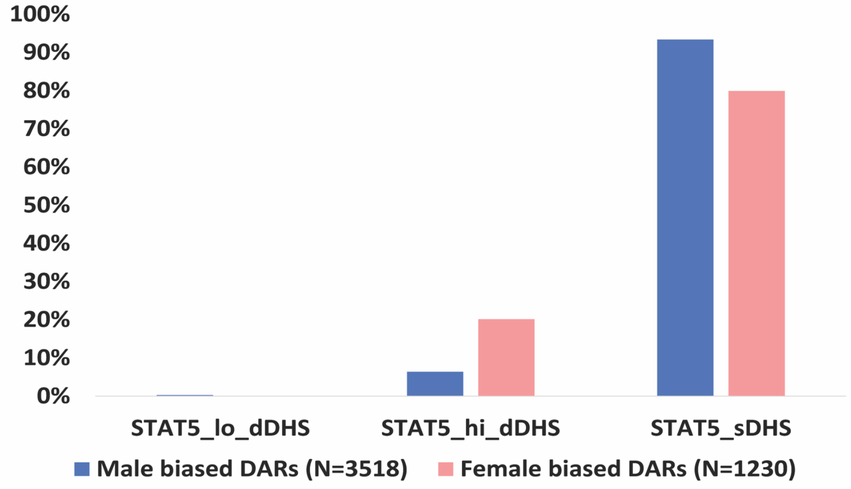


**C.**

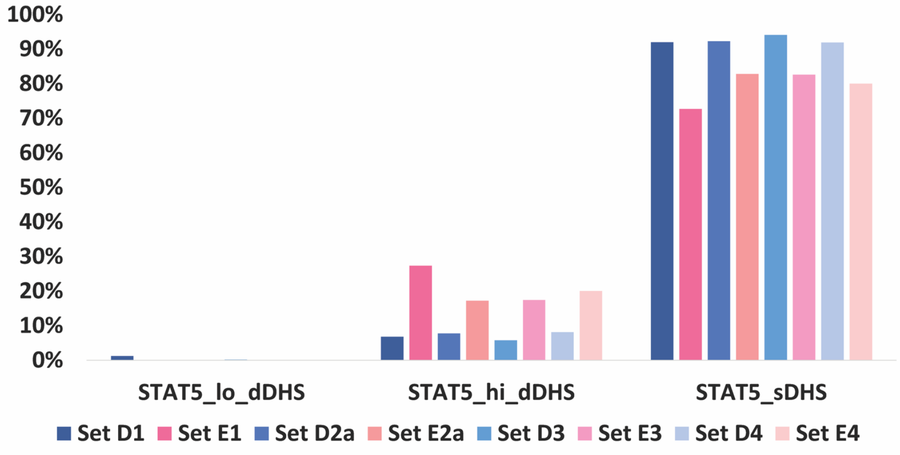

**Fig. S10**

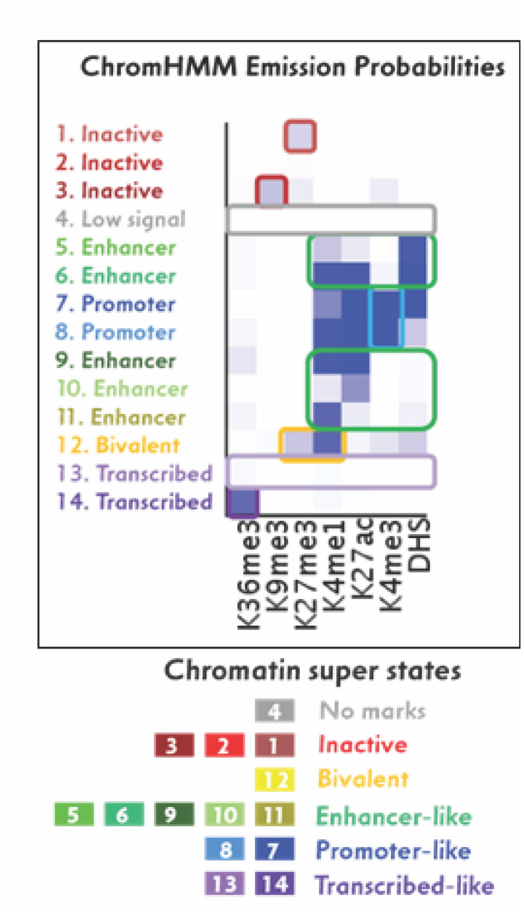
ChromHMM Emission Probabilities
1. Inactive
2. Inactive
3. Inactive
4. Low signal
5. Enhancer
6. Enhancer
7. Promoter
8. Promoter
9. Enhancer
10. Enhancer
11. Enhancer
12. Bivalent
13. Transcribed
14. Transcribed
K36me3
K9me3
K27me3
K4me1
K27ac
K4me3
DHS
Chromatin super states
4 No marks
3 2 1 Inactive
12 Bivalent
5 6 9 10 11 Enhancer-like
8 7 Promoter-like
13 14 Transcribed-like

### Fig. S11

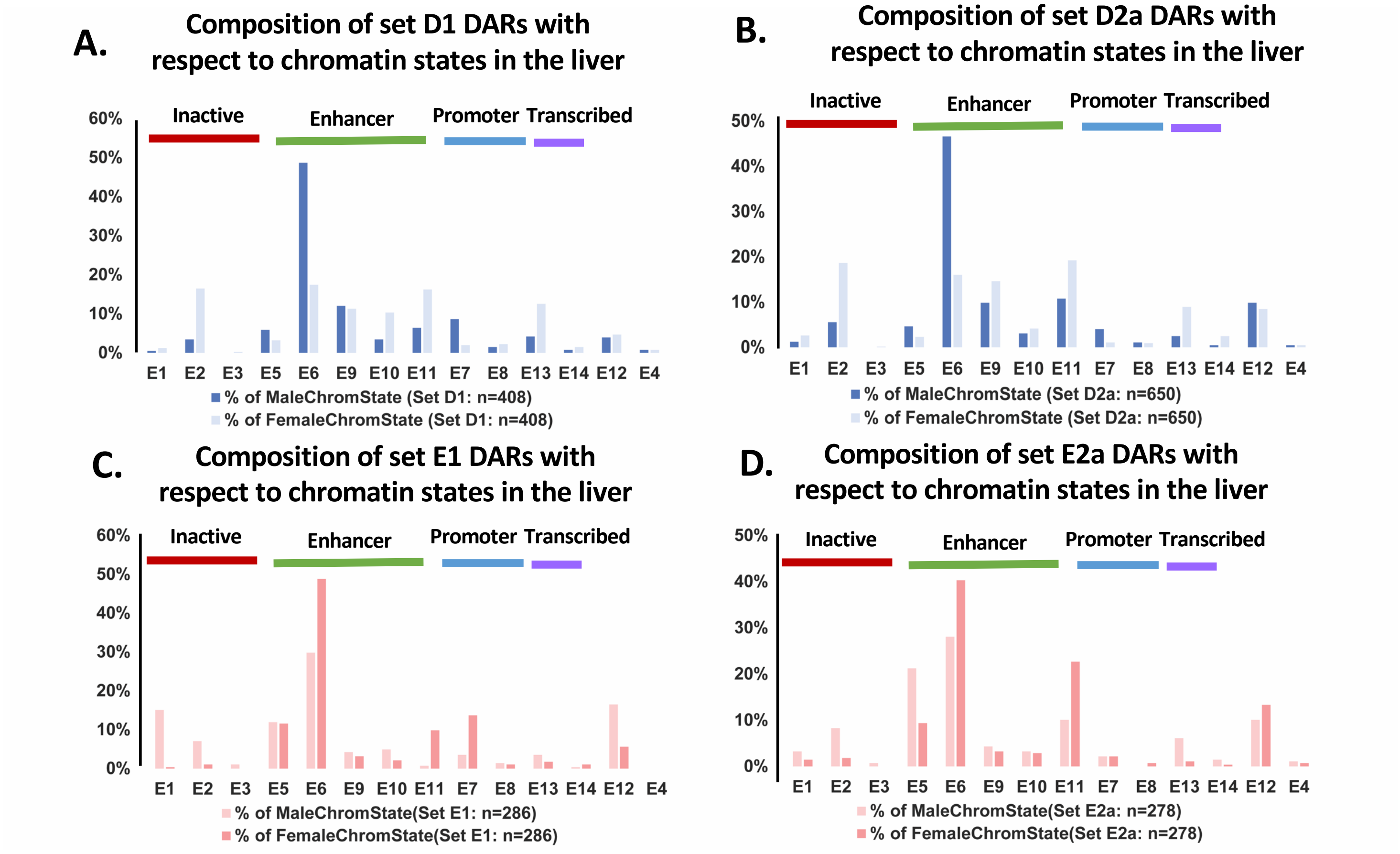
A.
Composition of set D1 DARs with respect to chromatin states in the liver
Inactive
Enhancer
Promoter
Transcribed
60%
50%
40%
30%
20%
10%
0%
E1 E2 E3 E5 E6 E9 E10 E11 E7 E8 E13 E14 E12 E4
% of MaleChromState (Set D1: n=408)
% of FemaleChromState (Set D1: n=408)
B.
Composition of set D2a DARs with respect to chromatin states in the liver
Inactive
Enhancer
Promoter
Transcribed
50%
40%
30%
20%
10%
0%
E1 E2 E3 E5 E6 E9 E10 E11 E7 E8 E13 E14 E12 E4
% of MaleChromState (Set D2a: n=650)
% of FemaleChromState (Set D2a: n=650)
C.
Composition of set E1 DARs with respect to chromatin states in the liver
Inactive
Enhancer
Promoter
Transcribed
60%
50%
40%
30%
20%
10%
0%
E1 E2 E3 E5 E6 E9 E10 E11 E7 E8 E13 E14 E12 E4
% of MaleChromState(Set E1: n=286)
% of FemaleChromState(Set E1: n=286)
D.
Composition of set E2a DARs with respect to chromatin states in the liver
Inactive
Enhancer
Promoter
Transcribed
50%
40%
30%
20%
10%
0%
E1 E2 E3 E5 E6 E9 E10 E11 E7 E8 E13 E14 E12 E4
% of MaleChromState(Set E2a: n=278)
% of FemaleChromState(Set E2a: n=278)

### Fig. S12

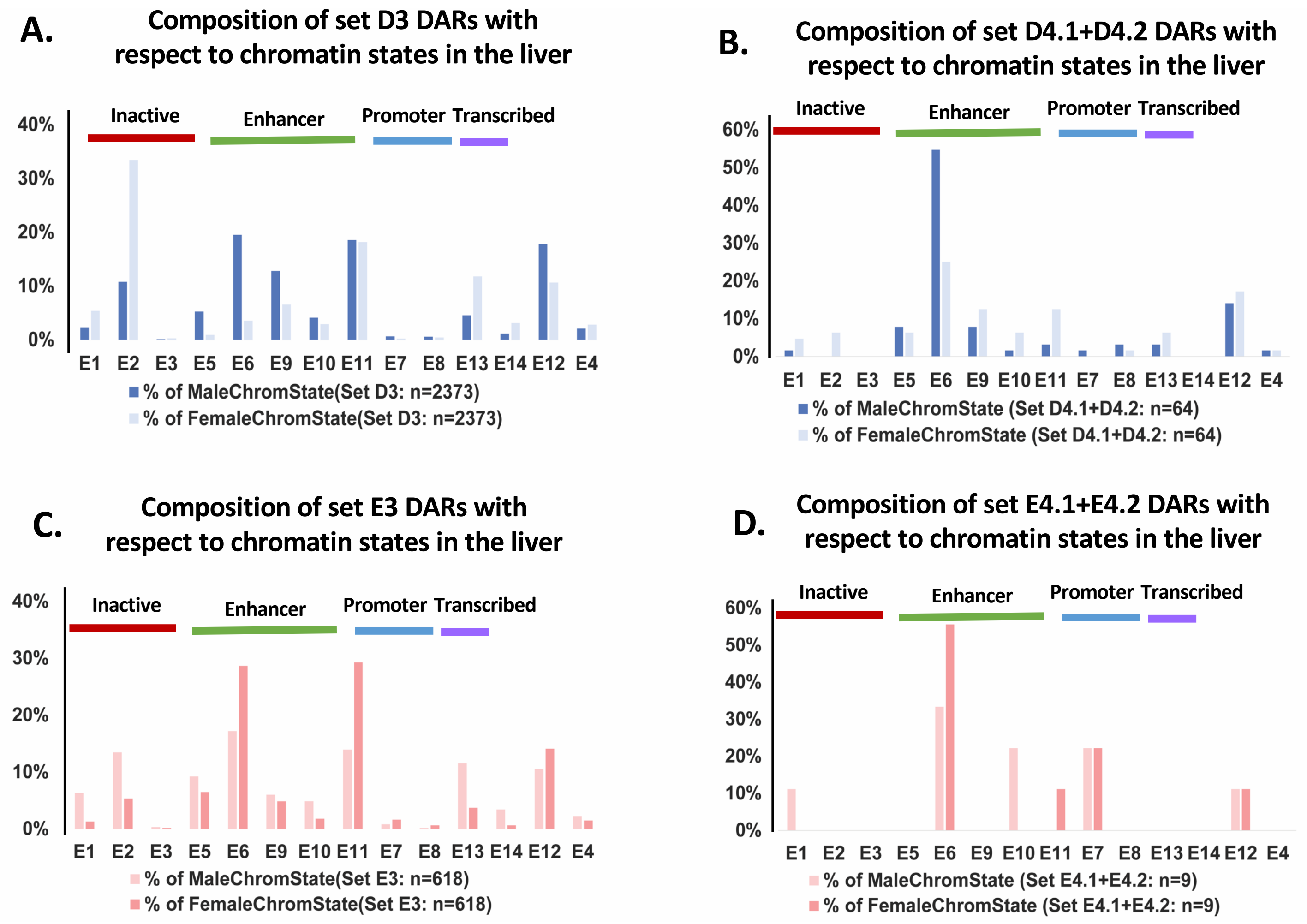
A.
Composition of set D3 DARs with respect to chromatin states in the liver
Inactive
Enhancer
Promoter
Transcribed
40%
30%
20%
10%
0%
E1 E2 E3 E5 E6 E9 E10 E11 E7 E8 E13 E14 E12 E4
% of MaleChromState(Set D3: n=2373)
% of FemaleChromState(Set D3: n=2373)
B.
Composition of set D4.1+D4.2 DARs with respect to chromatin states in the liver
Inactive
Enhancer
Promoter
Transcribed
60%
50%
40%
30%
20%
10%
0%
E1 E2 E3 E5 E6 E9 E10 E11 E7 E8 E13 E14 E12 E4
% of MaleChromState (Set D4.1+D4.2: n=64)
% of FemaleChromState (Set D4.1+D4.2: n=64)
C.
Composition of set E3 DARs with respect to chromatin states in the liver
Inactive
Enhancer
Promoter
Transcribed
40%
30%
20%
10%
0%
E1 E2 E3 E5 E6 E9 E10 E11 E7 E8 E13 E14 E12 E4
% of MaleChromState(Set E3: n=618)
% of FemaleChromState(Set E3: n=618)
D.
Composition of set E4.1+E4.2 DARs with respect to chromatin states in the liver
Inactive
Enhancer
Promoter
Transcribed
60%
50%
40%
30%
20%
10%
0%
E1 E2 E3 E5 E6 E9 E10 E11 E7 E8 E13 E14 E12 E4
% of MaleChromState (Set E4.1+E4.2: n=9)
% of FemaleChromState (Set E4.1+E4.2: n=9)

**Fig. S13A**

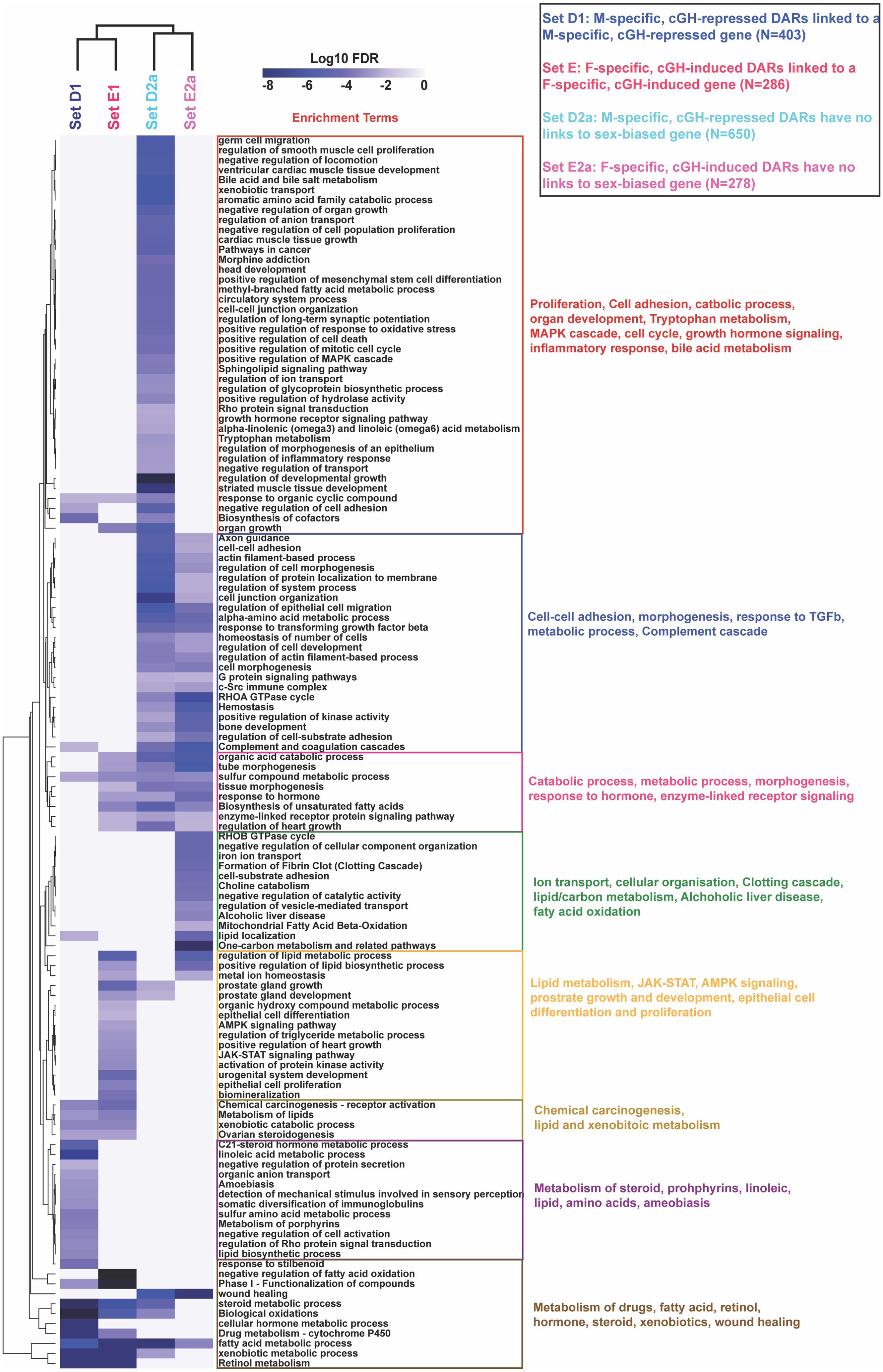
Set D1: M-specific, cGH-repressed DARs linked to a M-specific, cGH-repressed gene (N=403)
Set E: F-specific, cGH-induced DARs linked to a F-specific, cGH-induced gene (N=286)
Set D2a: M-specific, cGH-repressed DARs have no links to sex-biased gene (N=650)
Set E2a: F-specific, cGH-induced DARs have no links to sex-biased gene (N=278)
Log10 FDR
-8
-6
-4
-2
0
Set D1
Set E1
Set D2a
Set E2a
Enrichment Terms
germ cell migration
regulation of smooth muscle cell proliferation
negative regulation of locomotion
ventricular cardiac muscle tissue development
Bile acid and bile salt metabolism
xenobiotic transport
aromatic amino acid family catabolic process
negative regulation of organ growth
regulation of anion transport
negative regulation of cell population proliferation
cardiac muscle tissue growth
Pathways in cancer
Morphine addiction
head development
positive regulation of mesenchymal stem cell differentiation
methyl-branched fatty acid metabolic process
circulatory system process
cell-cell junction organization
regulation of long-term synaptic potentiation
positive regulation of response to oxidative stress
positive regulation of cell death
positive regulation of mitotic cell cycle
positive regulation of MAPK cascade
Sphingolipid signaling pathway
regulation of ion transport
regulation of glycoprotein biosynthetic process
positive regulation of hydrolase activity
Rho protein signal transduction
growth hormone receptor signaling pathway
alpha-linolenic (omega3) and linoleic (omega6) acid metabolism
Tryptophan metabolism
regulation of morphogenesis of an epithelium
regulation of inflammatory response
negative regulation of transport
regulation of developmental growth
striated muscle tissue development
response to organic cyclic compound
negative regulation of cell adhesion
Biosynthesis of cofactors
organ growth
Proliferation, Cell adhesion, catbolic process, organ development, Tryptophan metabolism, MAPK cascade, cell cycle, growth hormone signaling, inflammatory response, bile acid metabolism
Axon guidance
cell-cell adhesion
actin filament-based process
regulation of cell morphogenesis
regulation of protein localization to membrane
regulation of system process
cell junction organization
regulation of epithelial cell migration
alpha-amino acid metabolic process
response to transforming growth factor beta
homeostasis of number of cells
regulation of cell development
regulation of actin filament-based process
cell morphogenesis
G protein signaling pathways
c-Src immune complex
RHOA GTPase cycle
Hemostasis
positive regulation of kinase activity
bone development
regulation of cell-substrate adhesion
Complement and coagulation cascades
Cell-cell adhesion, morphogenesis, response to TGFb, metabolic process, Complement cascade
organic acid catabolic process
tube morphogenesis
sulfur compound metabolic process
tissue morphogenesis
response to hormone
Biosynthesis of unsaturated fatty acids
enzyme-linked receptor protein signaling pathway
regulation of heart growth
Catabolic process, metabolic process, morphogenesis, response to hormone, enzyme-linked receptor signaling
RHOB GTPase cycle
negative regulation of cellular component organization
iron ion transport
Formation of Fibrin Clot (Clotting Cascade)
cell-substrate adhesion
Choline catabolism
negative regulation of catalytic activity
regulation of vesicle-mediated transport
Alcoholic liver disease
Mitochondrial Fatty Acid Beta-Oxidation
lipid localization
One-carbon metabolism and related pathways
Ion transport, cellular organisation, Clotting cascade, lipid/carbon metabolism, Alchoholic liver disease, faty acid oxidation
regulation of lipid metabolic process
positive regulation of lipid biosynthetic process
metal ion homeostasis
prostate gland growth
prostate gland development
organic hydroxy compound metabolic process
epithelial cell differentiation
AMPK signaling pathway
regulation of triglyceride metabolic process
positive regulation of heart growth
JAK-STAT signaling pathway
activation of protein kinase activity
urogenital system development
epithelial cell proliferation
biomineralization
Lipid metabolism, JAK-STAT, AMPK signaling, prostrate growth and development, epithelial cell differentiation and proliferation
Chemical carcinogenesis - receptor activation
Metabolism of lipids
xenobiotic catabolic process
Ovarian steroidogenesis
Chemical carcinogenesis, lipid and xenobitoic metabolism
C21-steroid hormone metabolic process
linoleic acid metabolic process
negative regulation of protein secretion
organic anion transport
Amoebiasis
detection of mechanical stimulus involved in sensory perception
somatic diversification of immunoglobulins
sulfur amino acid metabolic process
Metabolism of porphyrins
negative regulation of cell activation
regulation of Rho protein signal transduction
lipid biosynthetic process
Metabolism of steroid, prohphyrins, linoleic, lipid, amino acids, ameobiasis
response to stilbenoid
negative regulation of fatty acid oxidation
Phase I - Functionalization of compounds
wound healing
steroid metabolic process
Biological oxidations
cellular hormone metabolic process
Drug metabolism - cytochrome P450
fatty acid metabolic process
xenobiotic metabolic process
Retinol metabolism
Metabolism of drugs, fatty acid, retinol, hormone, steroid, xenobiotics, wound healing

Fig. S13B

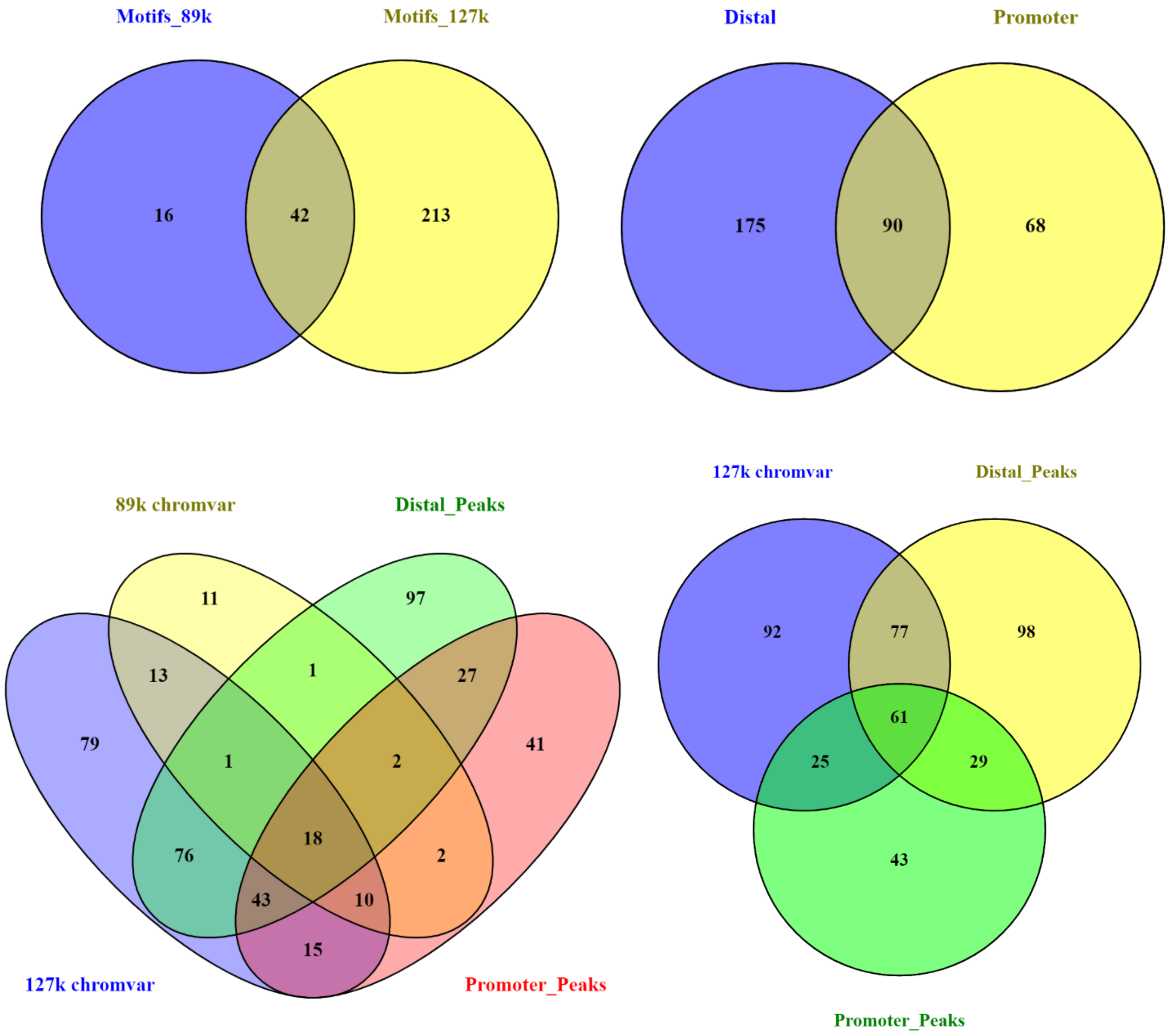
Motifs_89k
Motifs_127k
16
42
213
Distal
Promoter
175
90
68
89k chromvar
Distal_Peaks
11
97
13
1
27
79
1
2
41
18
76
2
43
10
15
127k chromvar
Promoter_Peaks
127k chromvar
Distal_Peaks
92
77
98
61
25
29
43
Promoter_Peaks

**Fig. S14**

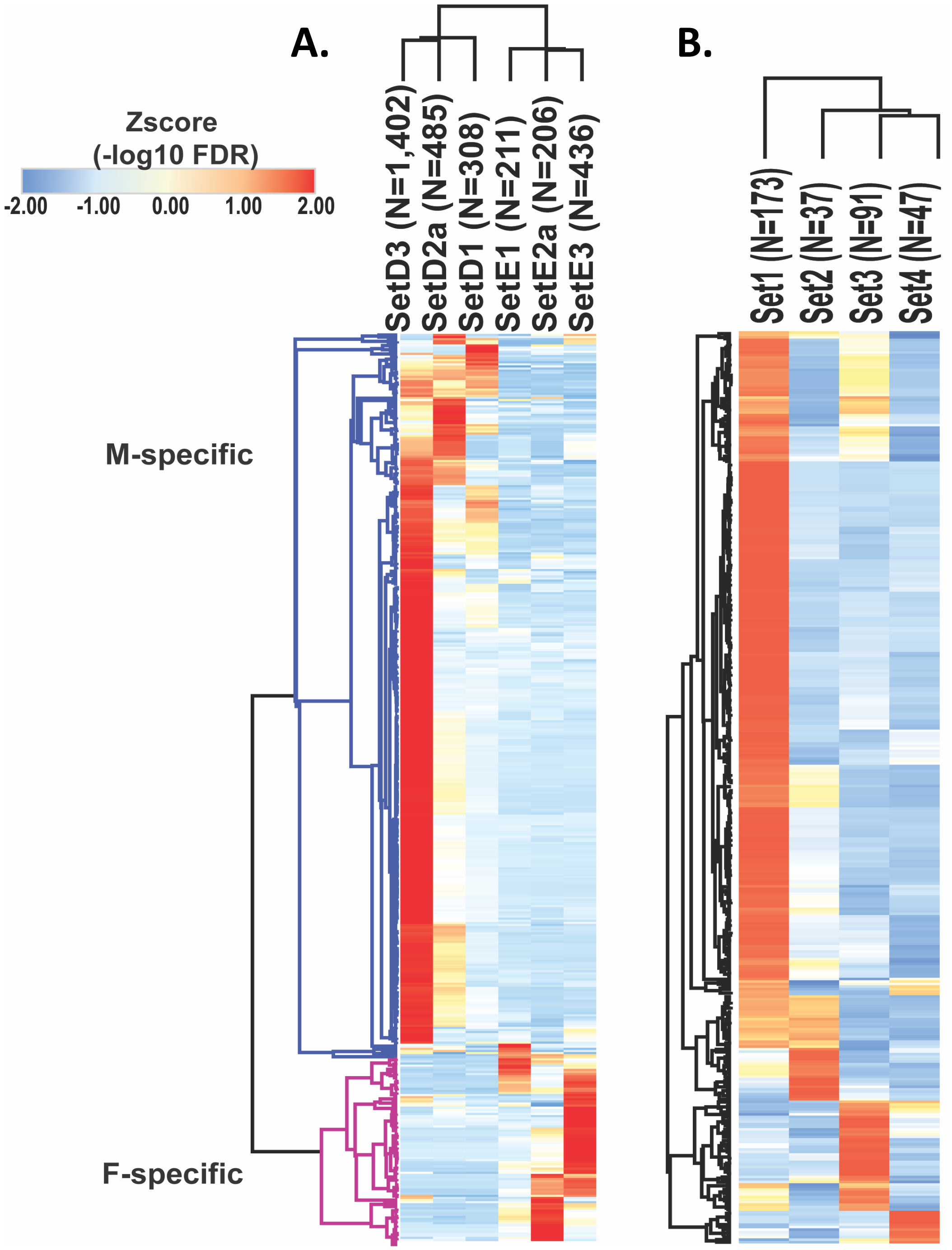


432 motifs enriched in hypox classes of sex-biased DAR sets

364 motifs enriched in hypox classes of sex-biased DAR sets

### Fig. S15

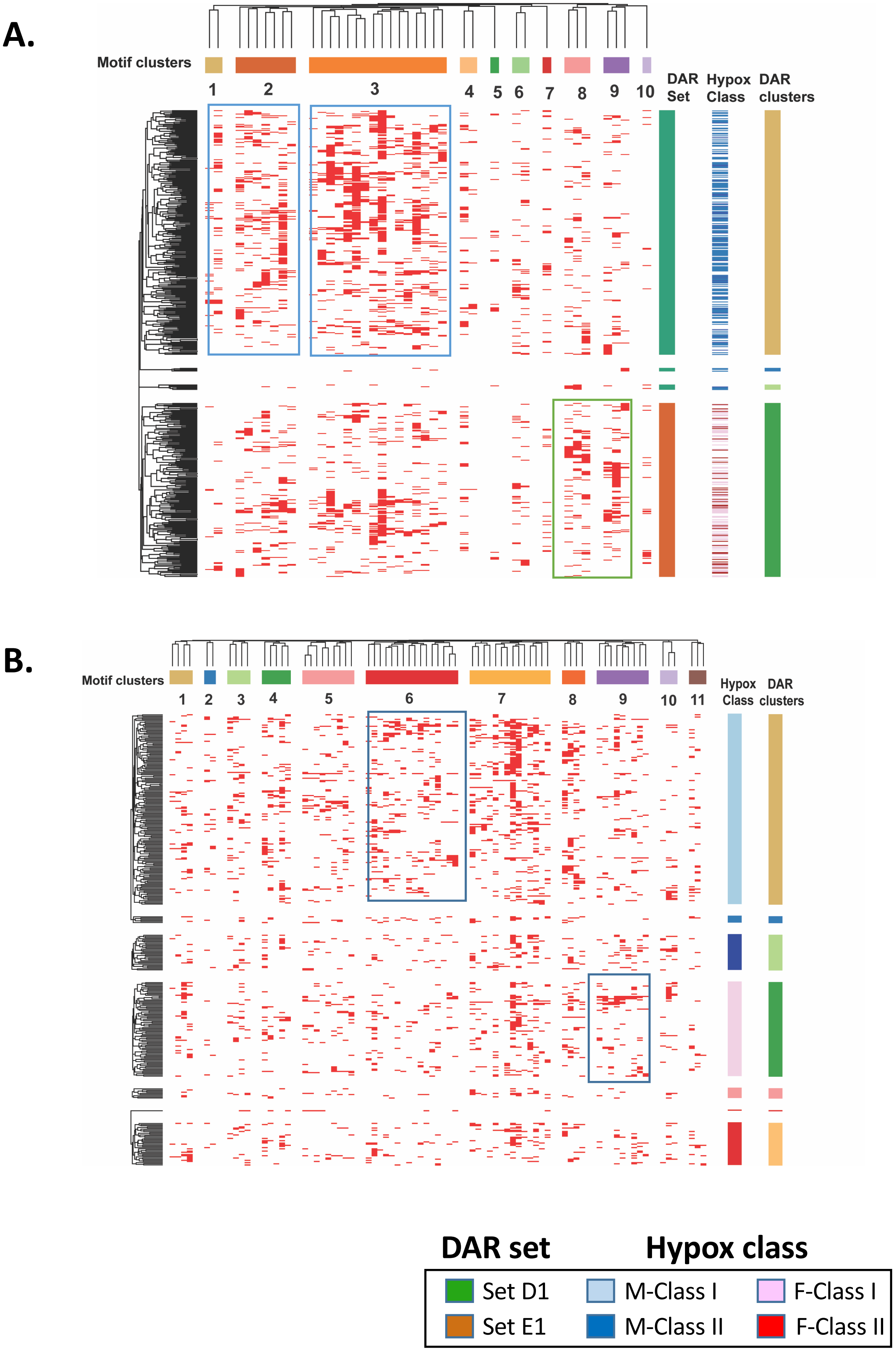
A.
Motif clusters
1
2
3
4
5
6
7
8
9
10
DAR Set
Hypox Class
DAR clusters
B.
Motif clusters
1
2
3
4
5
6
7
8
9
10
11
Hypox Class
DAR clusters
DAR set
Hypox class
Set D1
Set E1
M-Class I
M-Class II
F-Class I
F-Class II

### Fig. S16

#### A. Motif from M-Specific peaks linked to M-specific genes (Set D1)

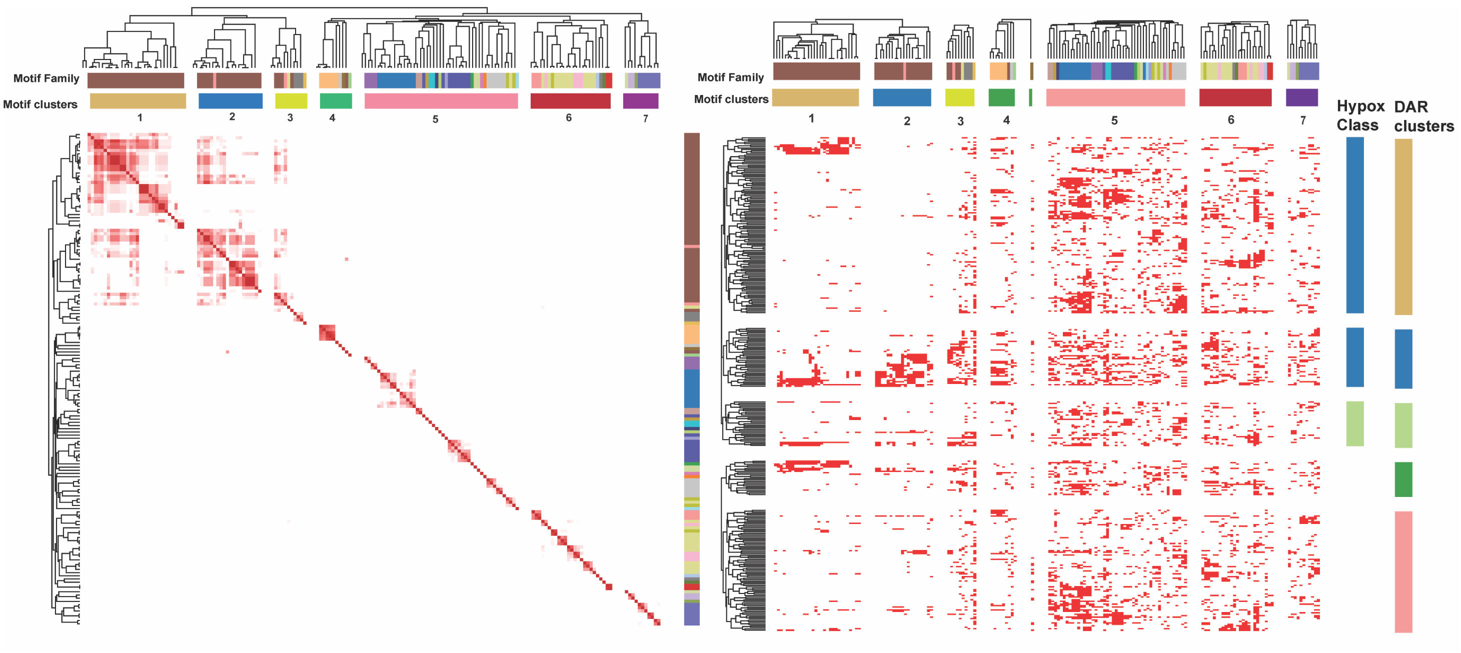


#### B. Motifs from F-Specific peaks linked to F-specific genes (Set E1)

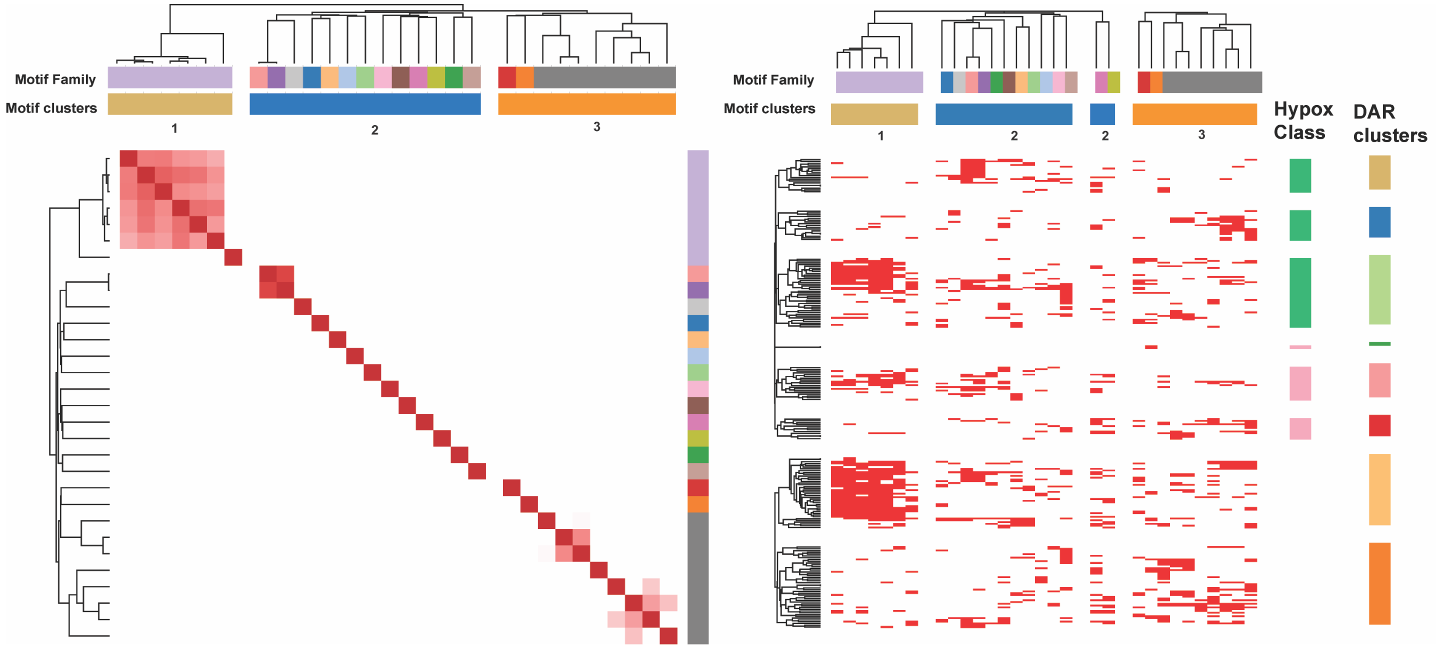


**Hypox class**

- M Class I
- M Class II
- F Class I
- F Class II

Fig. S16

C. Motif from M-Specific peaks linked to Male Hypox class I genes (Set1)

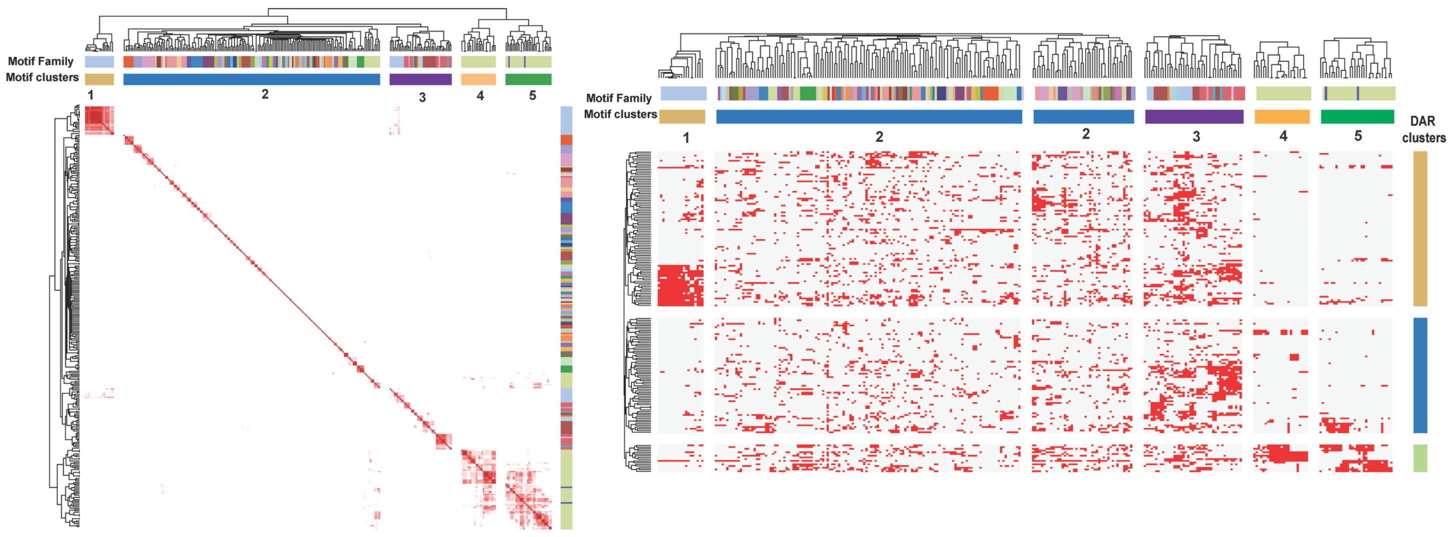


D. Motif from M-Specific peaks linked to Male Hypox class II genes (Set2)

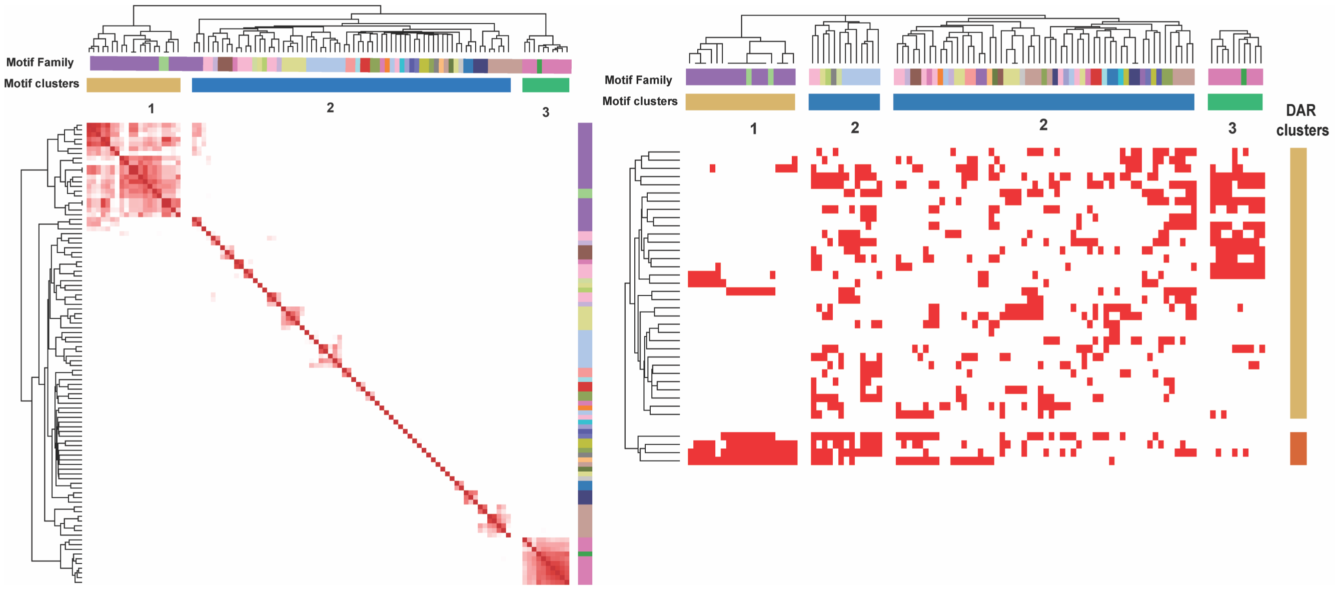

### Fig. S17

#### A. Motifs from F-Specific peaks linked to Female Hypox class I genes (Set3)

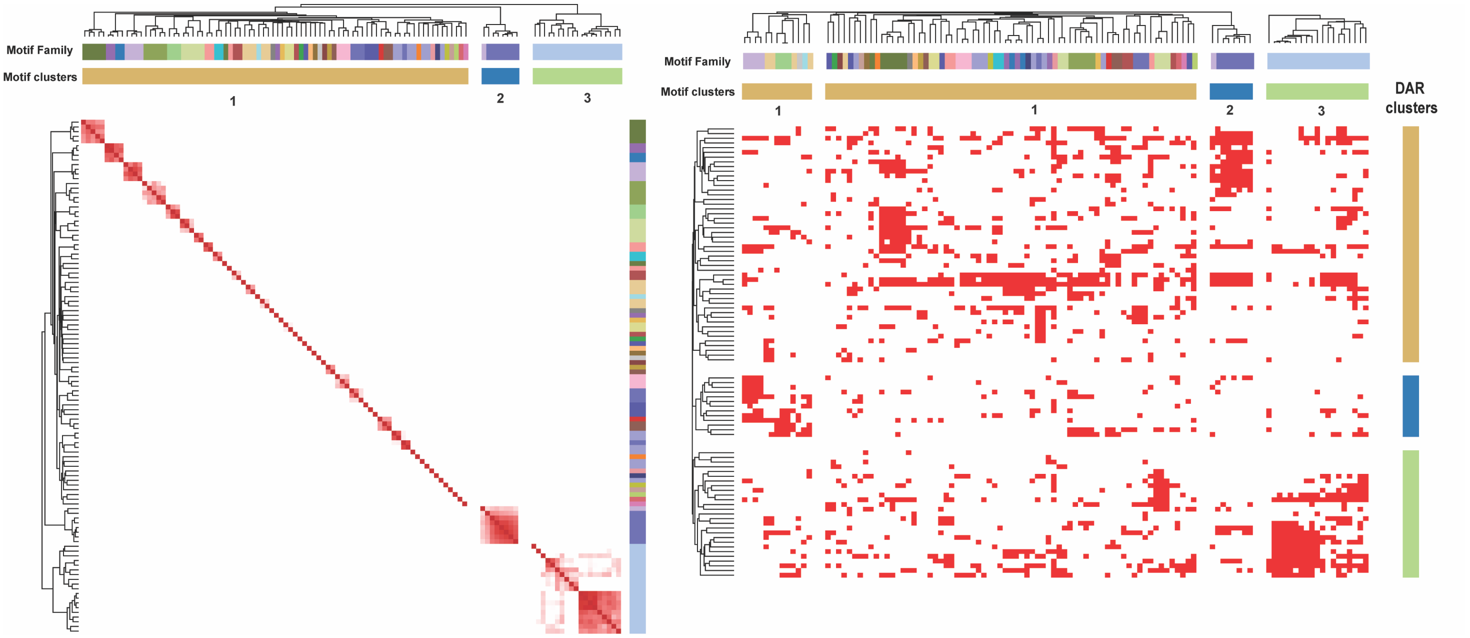


#### B. Motifs from F-Specific peaks linked to Female Hypox class II genes (Set4)

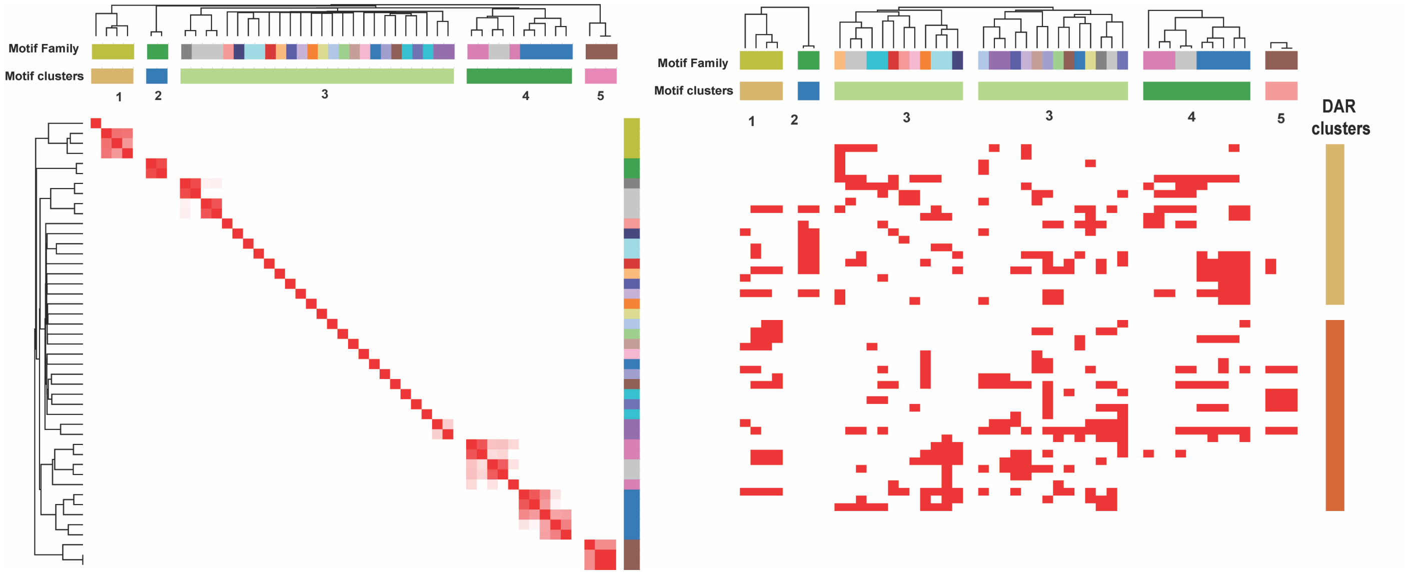

Fig. S18

A.

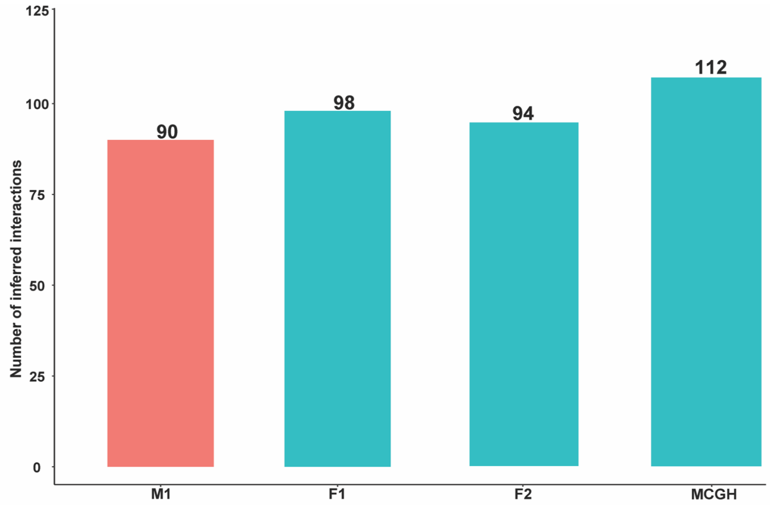

Number of inferred interactions
125
100
75
50
25
0
90
98
94
112
M1
F1
F2
MCGH


B.

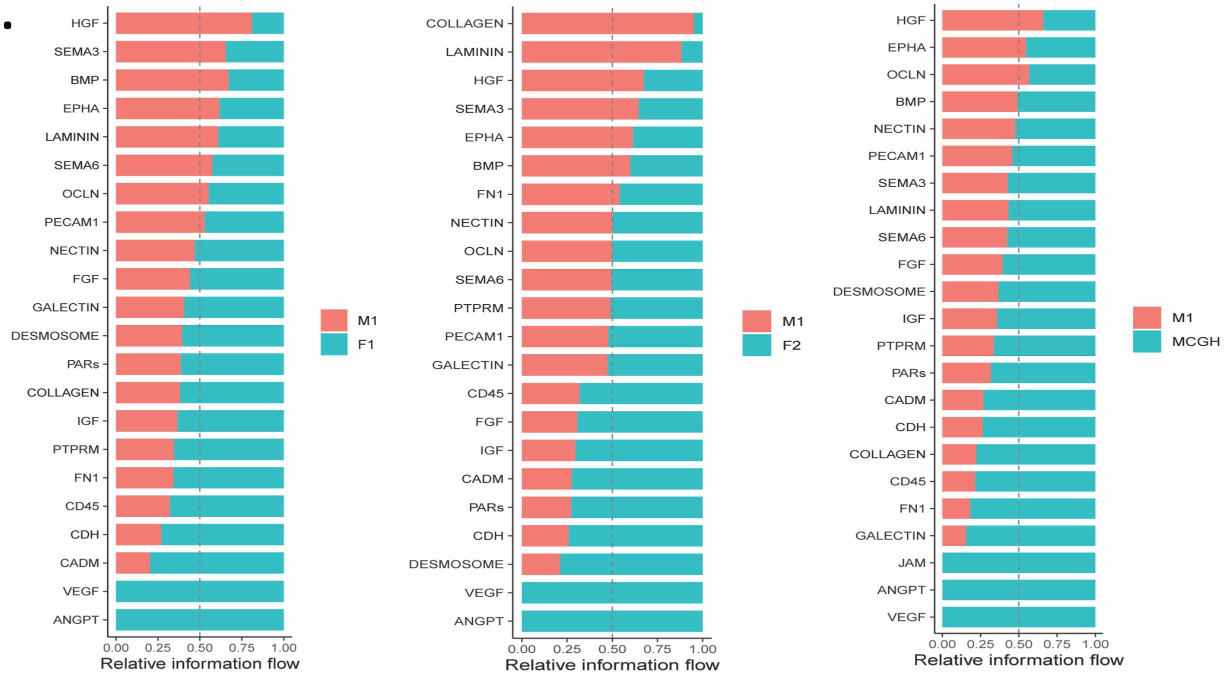

HGF
SEMA3
BMP
EPHA
LAMININ
SEMA6
OCLN
PECAM1
NECTIN
FGF
GALECTIN
DESMOSOME
PARs
COLLAGEN
IGF
PTPRM
FN1
CD45
CDH
CADM
VEGF
ANGPT
0.00 0.25 0.50 0.75 1.00
Relative information flow
M1
F1
COLLAGEN
LAMININ
HGF
SEMA3
EPHA
BMP
FN1
NECTIN
OCLN
SEMA6
PTPRM
PECAM1
GALECTIN
CD45
FGF
IGF
CADM
PARs
CDH
DESMOSOME
VEGF
ANGPT
0.00 0.25 0.50 0.75 1.00
Relative information flow
M1
F2
HGF
EPHA
OCLN
BMP
NECTIN
PECAM1
SEMA3
LAMININ
SEMA6
FGF
DESMOSOME
IGF
PTPRM
PARs
CADM
CDH
COLLAGEN
CD45
FN1
GALECTIN
JAM
ANGPT
VEGF
0.00 0.25 0.50 0.75 1.00
Relative information flow
M1
MCGH


C.

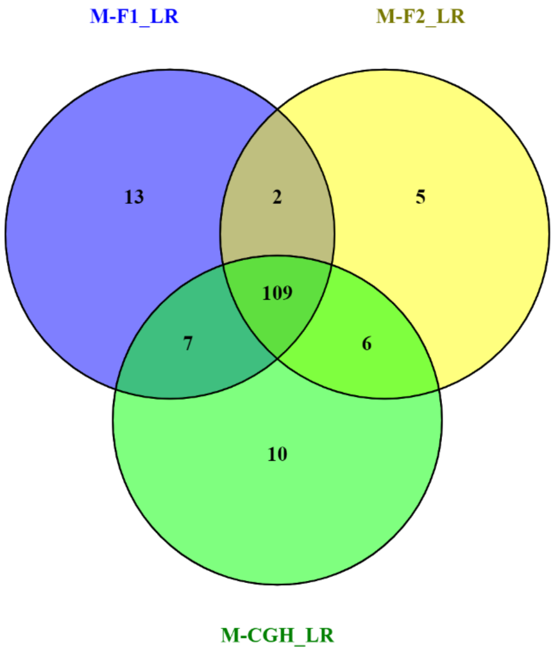

M-F1_LR
M-F2_LR
13
2
5
109
7
6
10
M-CGH_LR

**Fig. S19**

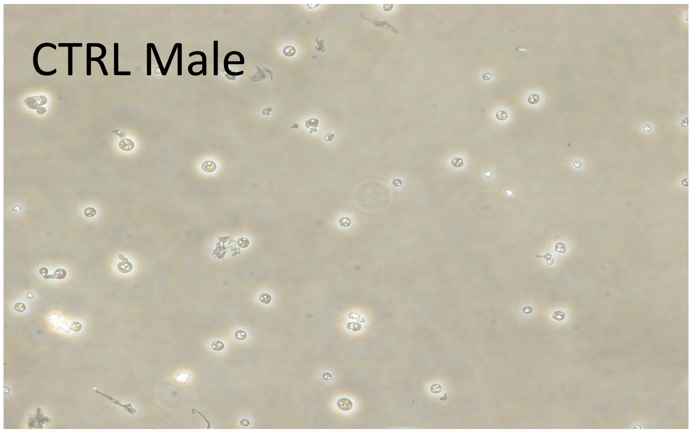


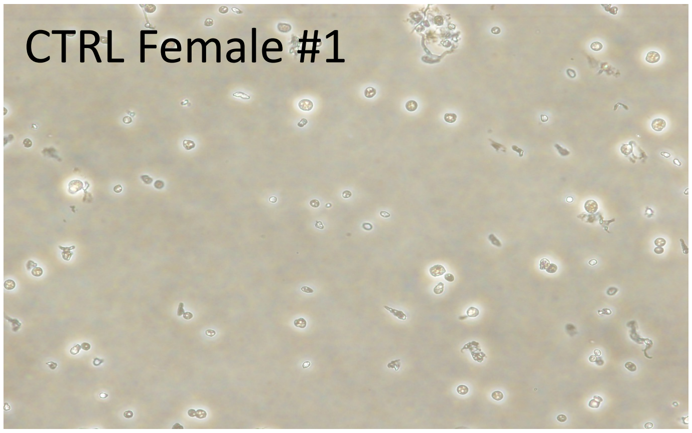

### Fig. S20

Control Male

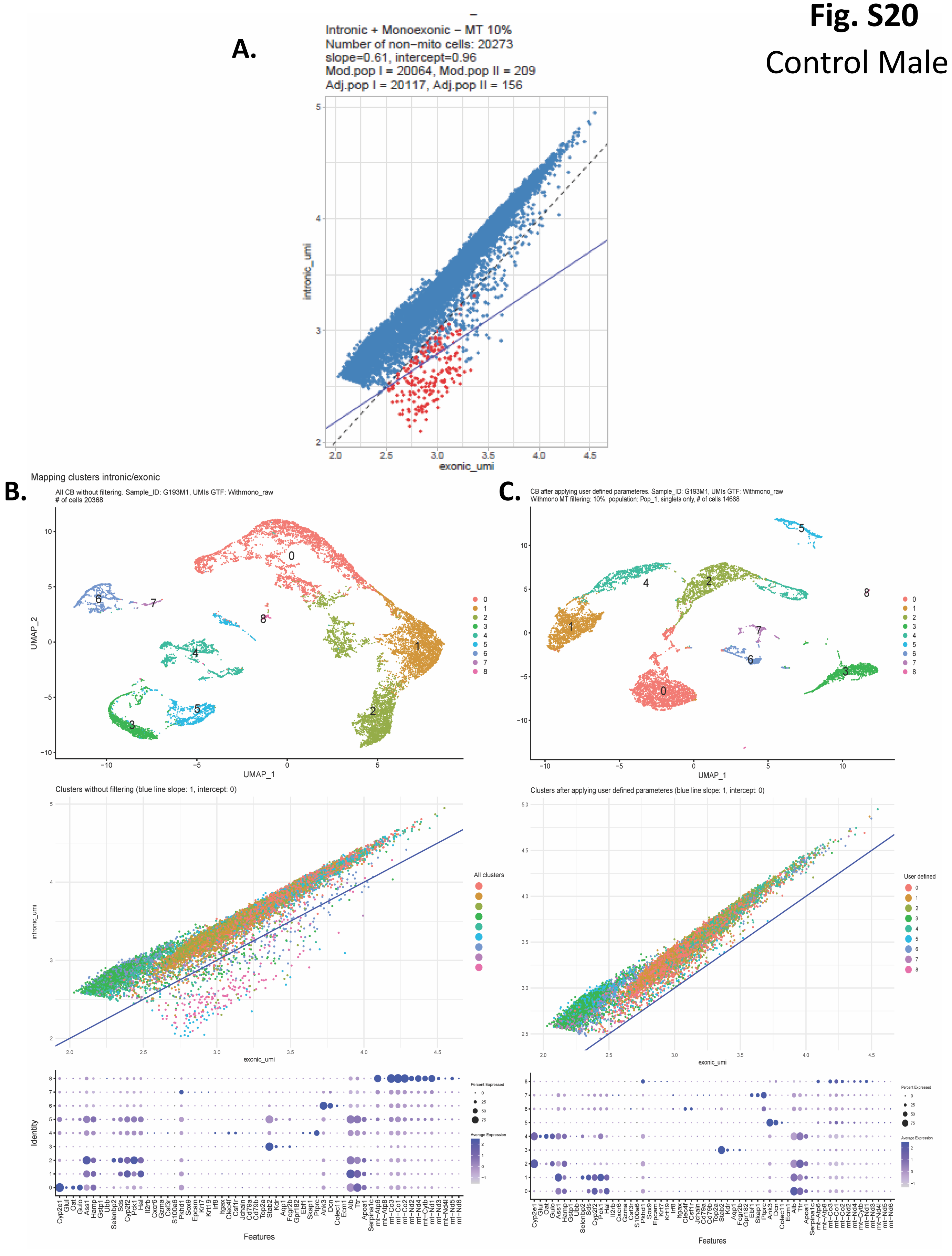

### Fig. S21

Control female (F1)

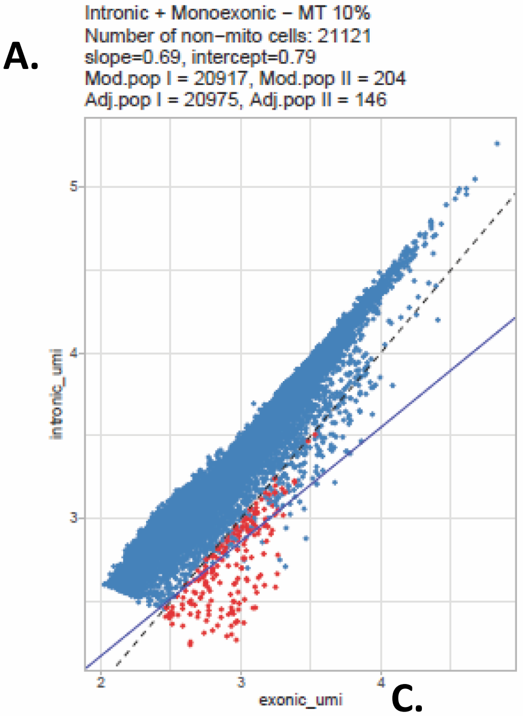


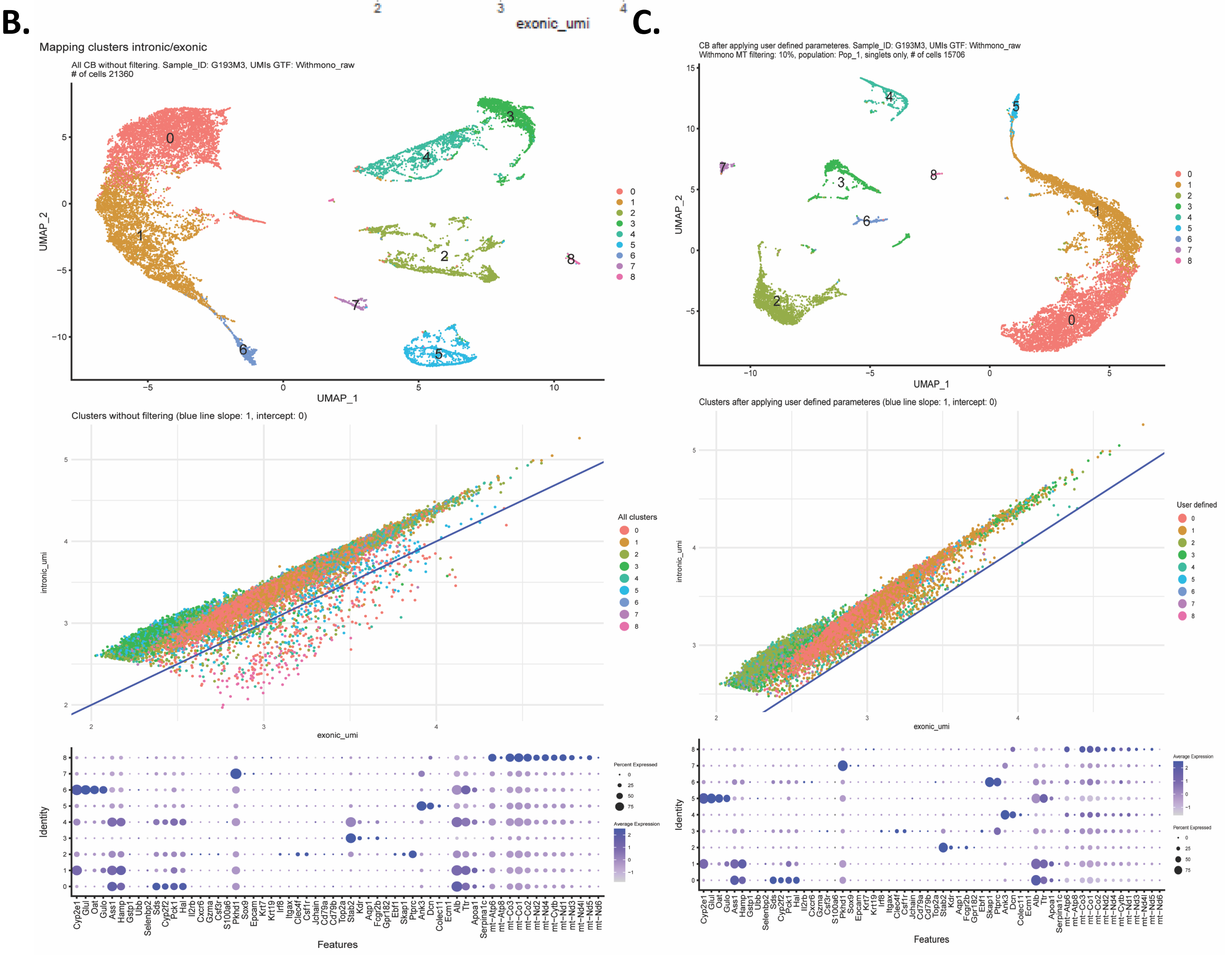

### Fig. S22

Control female (F2)

A.

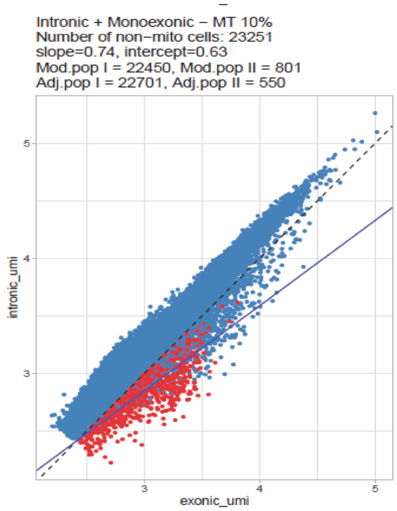


B.

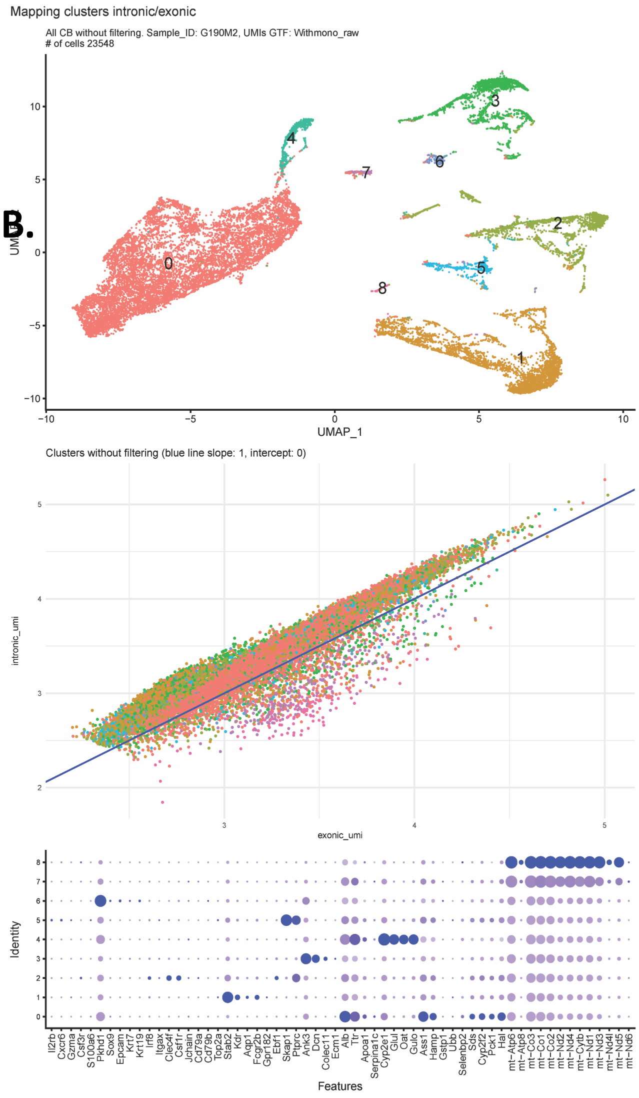


C.

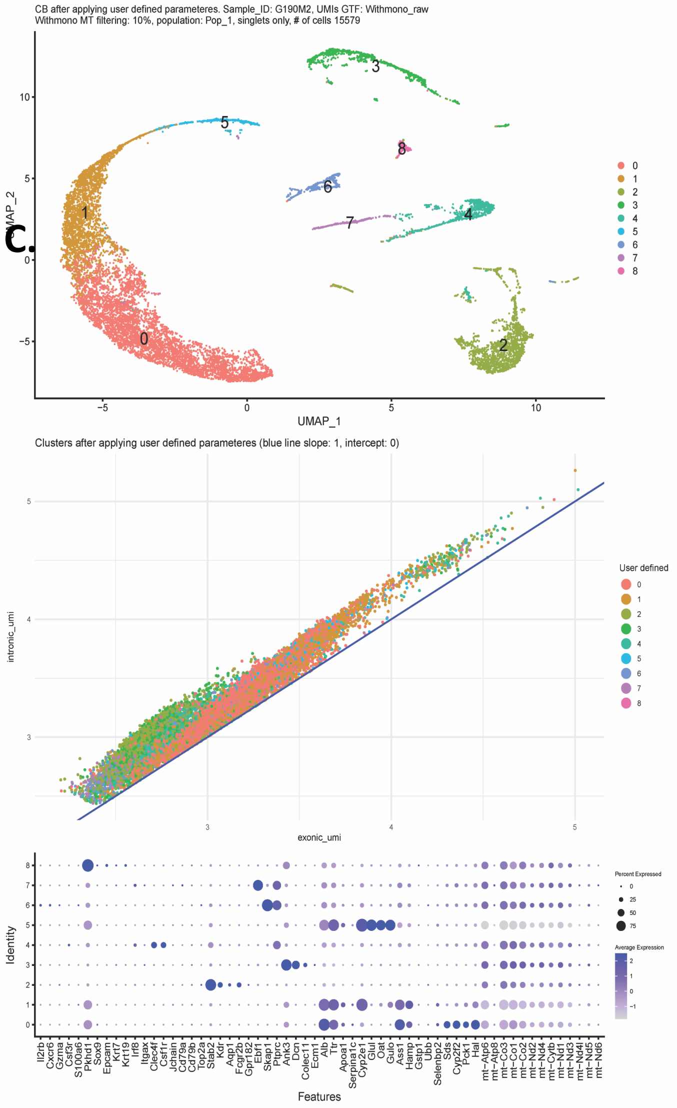

### Fig. S23

Male treated with Continuous growth hormone (MCGH)

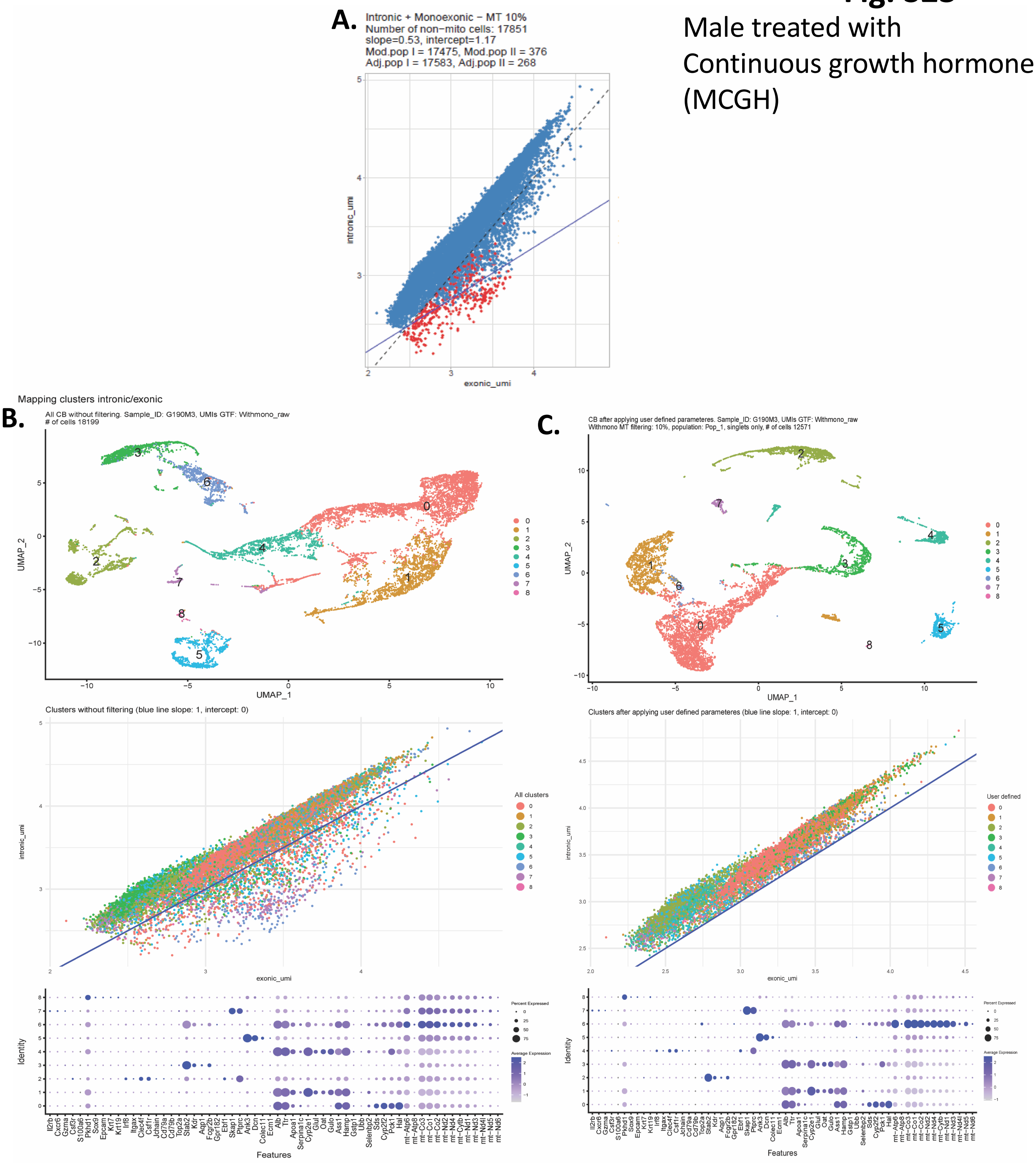

**Fig. S24**

A.

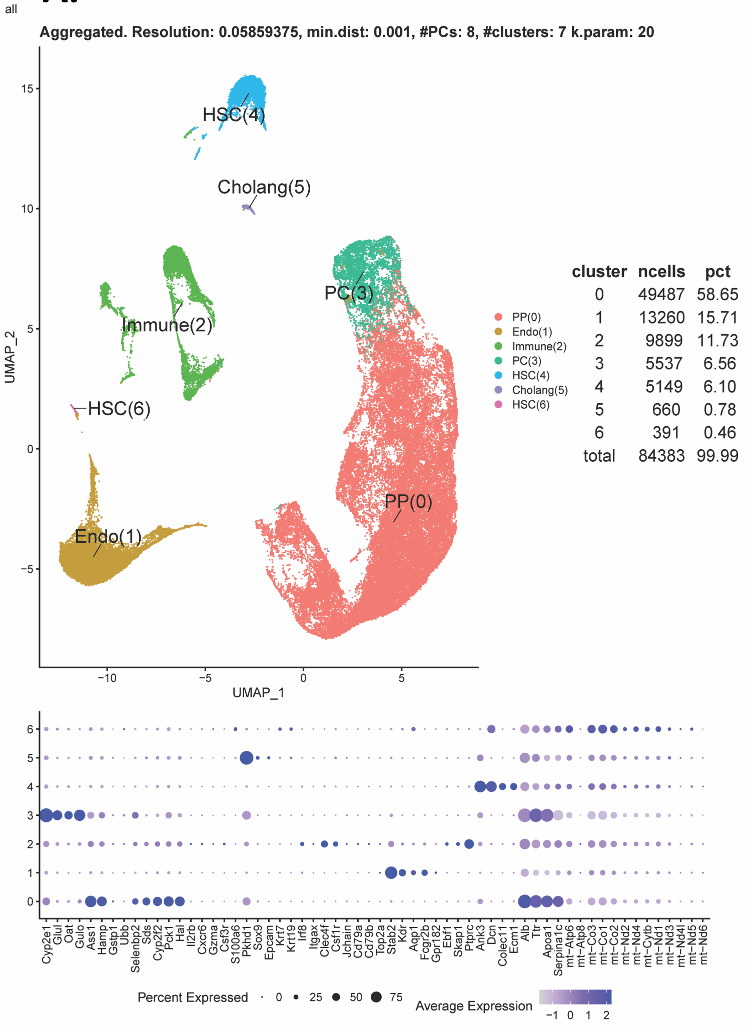
all
Aggregated. Resolution: 0.05859375, min.dist: 0.001, #PCs: 8, #clusters: 7 k.param: 20
HSC(4)
Cholang(5)
PC(3)
Immune(2)
HSC(6)
Endo(1)
PP(0)
UMAP_1
UMAP_2
PP(0)
Endo(1)
Immune(2)
PC(3)
HSC(4)
Cholang(5)
HSC(6)
cluster ncells pct
0 49487 58.65
1 13260 15.71
2 9899 11.73
3 5537 6.56
4 5149 6.10
5 660 0.78
6 391 0.46
total 84383 99.99
Cyp2e1 Glul Oat Gulo Ass1 Hamp Gstp1 Ubb Selenbp2 Sds Cyp2f2 Pck1 Hal Il2rb Cxcr6 Gzma Csf3r S100a6 Pkhd1 Sox9 Epcam Krt7 Krt19 Irf8 Itgax Clec4f Csf1r Jchain Cd79a Cd79b Top2a Stab2 Kdr Aqp1 Fcgr2b Gpr182 Ebf1 Skap1 Ptprc Ank3 Dcn Colec11 Ecm1 Alb Ttr Apoa1 Serpina1c mt−Atp6 mt−Atp8 mt−Co3 mt−Co1 mt−Co2 mt−Nd2 mt−Nd4 mt−Cytb mt−Nd1 mt−Nd3 mt−Nd4l mt−Nd5 mt−Nd6
Percent Expressed 0 25 50 75
Average Expression −1 0 1 2

B.

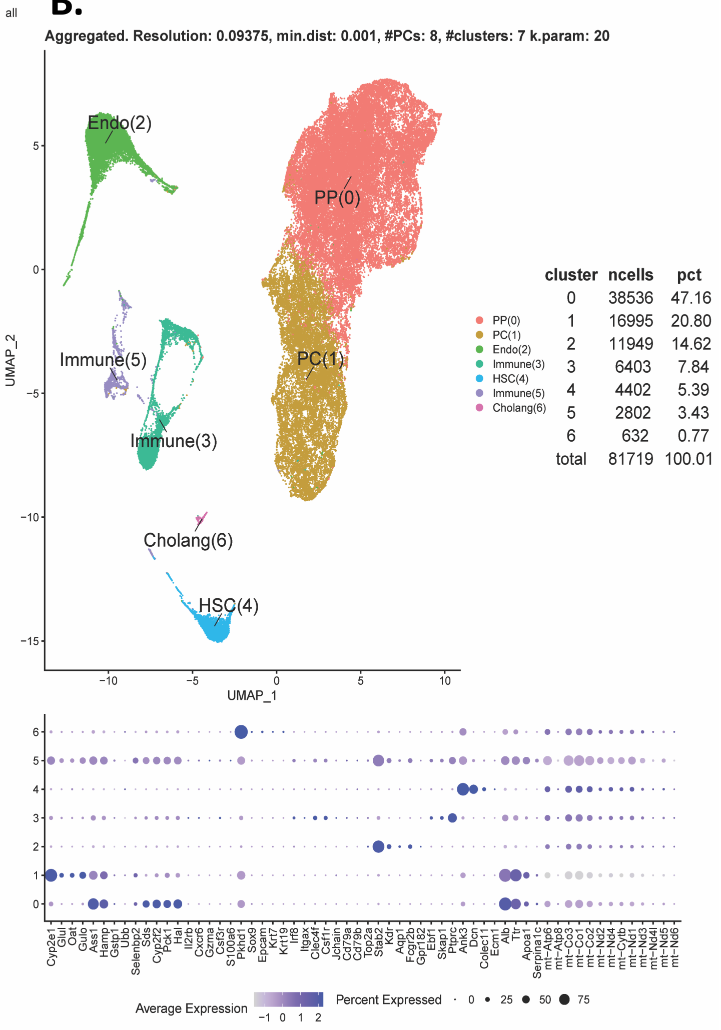
all
Aggregated. Resolution: 0.09375, min.dist: 0.001, #PCs: 8, #clusters: 7 k.param: 20
Endo(2)
PP(0)
Immune(5)
PC(1)
Immune(3)
Cholang(6)
HSC(4)
UMAP_1
UMAP_2
PP(0)
PC(1)
Endo(2)
Immune(3)
HSC(4)
Immune(5)
Cholang(6)
cluster ncells pct
0 38536 47.16
1 16995 20.80
2 11949 14.62
3 6403 7.84
4 4402 5.39
5 2802 3.43
6 632 0.77
total 81719 100.01
Cyp2e1 Glul Oat Gulo Ass1 Hamp Gstp1 Ubb Selenbp2 Sds Cyp2f2 Pck1 Hal Il2rb Cxcr6 Gzma Csf3r S100a6 Pkhd1 Sox9 Epcam Krt7 Krt19 Irf8 Itgax Clec4f Csf1r Jchain Cd79a Cd79b Top2a Stab2 Kdr Aqp1 Fcgr2b Gpr182 Ebf1 Skap1 Ptprc Ank3 Dcn Colec11 Ecm1 Alb Ttr Apoa1 Serpina1c mt−Atp6 mt−Atp8 mt−Co3 mt−Co1 mt−Co2 mt−Nd2 mt−Nd4 mt−Cytb mt−Nd1 mt−Nd3 mt−Nd4l mt−Nd5 mt−Nd6
Average Expression −1 0 1 2
Percent Expressed 0 25 50 75
